# Storing long-lived memories via molecular error correction: a minimal mathematical model of Crick’s memory switch

**DOI:** 10.1101/2025.11.13.688304

**Authors:** John J. Vastola, Tejas Ramdas, Samuel J. Gershman

## Abstract

Cells store information in part by attaching molecular marks to proteins and DNA. But because marks can be randomly removed (e.g., due to ambient phosphatase activity) and added (e.g., due to ambient kinase activity), information encoding may be poor, and in the worst case marks may only store information on the time scale of molecular turnover. We identify a high-level strategy cells use to maintain encoding fidelity beyond the time scale of turnover— which we call molecular error correction—and find that it closely resembles an underexplored theoretical proposal by Francis Crick. To assess the effectiveness of molecular error correction, we construct and analyze several minimal mathematical models of molecular memory switches. We find that Crick-like error correction provides an efficient way to improve the lifetime of stored memories, especially compared to redundantly encoding information using a large number of molecules, but that it can yield false positives (and hence low information fidelity) when there is ambient marking activity; given these two competing concerns, the optimal level of error correction is moderate rather than arbitrarily high. We also find that combining error correction with redundant encoding can efficiently and robustly produce memories that last between ten and one hundred times longer than the turnover time scale. Our work provides insight regarding how to model and interrogate noisy molecular memory systems, suggests that error correction is a design principle of molecular memory, and quantitatively connects error correction to observable memory time scales.

## Introduction

One way cells store information is by attaching molecular marks to proteins and DNA. Phosphate groups and other post-translational modifications can alter protein function in response to a transient environmental stimulus [1]; for example, the calcium-dependent protein kinase CaMKII, which participates in cellular responses to calcium influx, becomes phosphorylated after a sufficiently strong calcium signal [2, 3]. Modifications to chromatin, including histone modifications (e.g., acetylation and methylation) and DNA methylation, can produce long-lasting changes in gene expression [4, 5] and help stably maintain cell fate [6]. Because these marks can produce long-lasting changes in intracellular dynamics, they can be viewed as a form of cellular memory.

Storing memory via molecular marks faces a fundamental problem not necessarily shared by other forms of memory: molecular components are continually replaced. Marks can be randomly removed by enzymes, and marked proteins can be degraded and replaced. In the absence of any process which combats molecular turnover, information could only be stored on the time scale of turnover, which may be short. Most human and mouse proteins, for example, turn over on a time scale of hours to days [7–9], which makes it unlikely that any single protein molecule is sufficient for long-term memory storage.

Prior work has identified different mechanisms by which cells combat turnover: DNA methylation is maintained by DNA methyltransferases [10, 11]; modified histones recruit ‘readers’ which cause their modifications to be ‘written’ to nearby histones [12, 13]; and proteins like CaMKII can maintain their phosphorylation state via autophosphorylation [2, 3, 14]. While these mechanisms are clearly distinct in their details and biological purpose (e.g., the type of memory stored), we claim that they reflect the same high-level strategy, which we call *molecular error correction*. Molecular error correction involves two key ingredients: using three states (*unmarked, partially marked*, and *fully marked*) rather than only two (*unmarked* and *marked*), and a mechanism that quickly drives a partially marked system into its fully marked state. If the system stores information by being placed into its fully marked state, but molecular turnover erroneously pushes it into its partially marked state, the mechanism ensures the system usually returns to its fully marked state (Fig. 1a). Because ‘errors’ are rapidly corrected, information persists longer than the turnover time scale.

**Figure 1.**
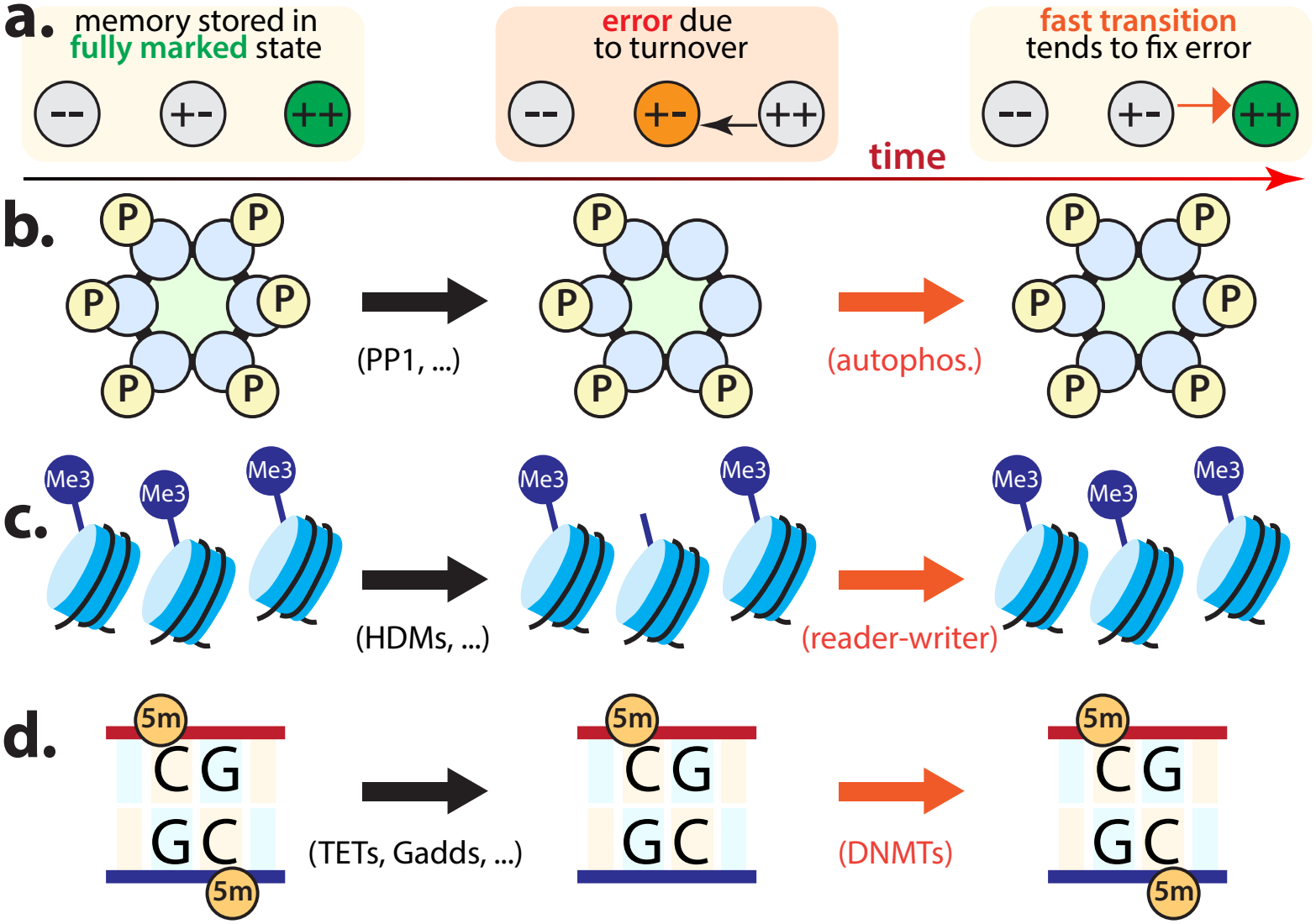
Different processes share an error correction motif. (a) Schematic of molecular error correction. If turnover erroneously drives the system into its partially marked state, it tends to quickly return to its fully marked state. (b) CaMKII autophosphorylation as molecular error correction. (c) Histone mark maintenance as molecular error correction. (d) DNA (cytosine) methylation maintenance as molecular error correction.

In some biological contexts, like gene regulatory networks, dynamical motifs like attractors prevent information degradation [15, 16]: the turnover of individual proteins matters less, since regulatory interactions between genes prevent protein concentrations from deviating too much from a specific value. But the attractor concept, which is typically used to describe systems with continuous state spaces (e.g., protein concentrations), is not necessarily appropriate for noisy systems whose state space is small and discrete, like a single protein with two phosphorylation states. One reason molecular error correction is interesting is that it adapts the concept of ‘slow’ regions of state space [17–19]—i.e., sets of states which are ‘difficult to leave’—to noisy, few-state systems.

The idea of molecular error correction is not new, but in the context of molecular memory it is curiously undertheorized. Although Francis Crick described it in an influential and widely cited 1984 letter to *Nature* [20], it does not appear to have ever been explicitly modeled. There are obvious questions concerning its effectiveness at maintaining information beyond the time scale of turnover, and in particular regarding how it fares relative to the simpler strategy of using a large population of molecules to redundantly encode information [21]. The answers to quantitative questions, like how much error correction is ideal, are also unclear. In the absence of clarifying experiments, a useful way to get at these questions is via mathematical modeling [22, 23].

In what follows, our goal is to study the effectiveness of molecular error correction via a detailed analysis of a minimal mathematical model of it. We call our model a *Crick switch* because its precise form follows Crick’s recommendation. Our approach has two primary advantages over previous modeling work on similar questions: first, our model is simple enough that we can solve it exactly, which yields interesting quantitative insights about how error correction for memory storage works; and second, the abstract nature of our model allows us to connect it to a variety of systems that store information via molecular marks, including CaMKII and epigenetic systems.

## Models and metrics

### Different processes share an error correction motif

Our modeling goal is to capture the high-level phenomenology of several examples of molecular error correction—CaMKII autophosphorylation, histone mark maintenance via reader-writer enzymes, and DNA methylation maintenance via DNA methyltransferases—each of which is associated with very different biological substrates and time scales. Common to these examples are (i) three high-level marking states; (ii) a *turnover time scale τ* on which information is retained in the absence of error correction; (iii) an *error correction time scale τ*_fast_ < *τ*; and (iv) a *memory time scale t*_*\**_ > *τ*. Before describing our model, we review the relevant biology and provide rough numbers for *τ*, *τ*_fast_, and *t*_*\**_ below. Because these time scales are measured differently in each case, comparisons should be understood as approximate and mainly at the level of orders of magnitude.

#### CaMKII autophosphorylation

Calcium/calmodulin-dependent protein kinase II (CaMKII) modulates cellular responses to calcium influx and is abundant in postsynaptic densities [3]. A strong calcium signal causes calcium to bind to calmodulin (CaM), which then binds to CaMKII and causes it to phosphorylate itself, allowing it to phosphorylate downstream targets. CaMKII has a dodecameric structure and consists of two stacked rings, each with six subunits. Each of these 12 subunits can be phosphorylated at T286, and these marks are vulnerable to both ambient phosphatase (e.g., PP1) activity and subunit degradation.

One way CaMKII maintains a short-term memory of a transient calcium signal is through *autophosphorylation*: subunits can phosphorylate their neighbors within the same ring, and hence ‘correct errors’ due to turnover (Fig. 1b). Although this property caused CaMKII to receive sub-stantial attention as a potential memory molecule that stores information for years or decades [2, 14, 24], modern experiments revealed that only subunits with bound Ca^2+^-CaM can be autophosphorylated, and hence that the time scale of CaMKII memory is constrained by Ca^2+^-CaM availability [25]. Fluorescence measurements indicate that kinases with no error correction (i.e., mutants which cannot undergo T286 autophosphorylation) retain information about a calcium signal for *τ* ~ 1 s, while the wild-type kinase retains information for *t*_*\**_ ~ 60 s [3, 26, 27]. Reported rates of T286 autophosphorylation under strongly activating conditions are approximately 1 *–* 10 s^−1^, corresponding to an error-correction time scale *τ*_fast_ ~ 0.1 − 1 s [28, 29].

#### Reader-writer-mediated histone mark restoration

Histones carry marks—most prominently, acetylation and methylation—that regulate gene expression and store information [30, 31]. While most classic work views marks as encoding information about developmental state, stimulusdependent changes in histone marking have been implicated in learning and memory, especially via contextual fear conditioning experiments [32, 33]. Marks are vulnerable to turnover due to demodifying enzymes, histone subunit exchange, and especially dilution after DNA replication.

For repressive methylation marks like H3K27me3 and H3K9me3, reader-writer enzymes [12, 13] like PRC2 oppose this turnover: ‘reading’ the mark on one histone tends to cause it to be ‘written’ to neighboring histones. The histones within a local chromatin domain, then, tend to be either *mostly unmarked, mostly marked*, or in a less stable mixed state that reader-writer enzymes drive into the mostly marked state. Time scales of histone marking dynamics vary, but the main source of turnover in rapidly dividing cells is replication-coupled dilution, which happens every *τ* ~ 12−24 h for mammalian cells. H3K27me3-rich regions recover over approximately *τ*_fast_ ~ 2−6 h [34], and marks can persist over many cell divisions, with *t*_*\**_ ≳ 8 days in an inducible Polycomb system [35].

#### DNA maintenance methylation

The classic example of molecular error correction, which Crick explicitly cited as a motivation for his proposal [20], is the maintenance of DNA (cytosine) methylation. In a CpG sequence context, cytosines on opposite strands are methylated, so there is a symmetrically methylated CpG dyad; turnover (or, less randomly, cell division) can remove one methyl group, producing a hemimethylated state. On a fast time scale, site-specific DNA methyl-transferases (DNMTs) can ‘read’ the methylated residue of one strand and ‘write’ a methyl group to the complementary cytosine, producing a fully-methylated pair [10, 11]. Due to rapid error correction, these marks last longer than the time scale on which any single mark is removed.

There is wide variation in measurements of epigenetic mark turnover, but because epigenetic marks survive dilution due to DNA replication and can even be heritable, it is reasonable to expect that the error correction mechanism is extremely effective. For mammalian cells with a cell cycle duration *τ* ~ 12 − 24 h, remethylation after cell division occurs on a time scale *τ*_fast_ ~0.3 − 4 h [36, 37]. Maintenance fidelity is locus-dependent, but measured failure probabilities of a few percent per CpG per division imply persistence over tens of cell generations, corresponding roughly to *t*_*\**_ ~ several weeks in many dividing cells [38].

### Markov chain models of molecular memory switches

As the previous examples indicate, the minimal ingredients of a Crick-like molecular memory switch are three states and transitions between them due to turnover and error correction. To study the effectiveness of error correction while minimizing unnecessary detail, we model molecular memory switches as small, continuous-time Markov chains. Note that by treating transitions as stochastic, we account for observed randomness associated with molecular events like turnover, which becomes important when the number of molecules or states in the system is small [39–45]. In addition to studying the switches below, we also consider *N*-switch model variants consisting of *N* independent and identical copies of them. This allows us to explicitly compare the effectiveness of error correction with the ‘population-averaging’ strategy that simply uses more molecules.

#### The two-state model

As a baseline, we consider arguably the simplest molecular memory switch. The **two-state model** has two states: *unmarked* (−) and *marked* (+). We assume that turnover produces random transitions from the marked to unmarked state, and that marking noise (e.g., ambient kinase activity) produces transitions from the unmarked to marked state. Model behavior is fully characterized by the probability *p*_0_(*t*) of being unmarked and the probability *p*_1_(*t*) of being marked at time *t*. These probabilities evolve in time according to a **master equation**:

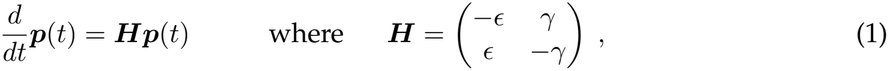

where ***p***(*t*) := (*p*_0_(*t*), *p*_1_(*t*))^*T*^, and the 2 × 2 matrix ***H*** is the transition matrix, or *Hamiltonian* [46, 47]. *ϵ* ≥ 0 is the probability per unit time of transitioning to the marked state and *γ* > 0 is the turnover rate. Note that the off-diagonal entries of ***H*** are the small-time transition rates between different states, and that the diagonal entries are chosen so that the columns of ***H*** sum to zero (which ensures that *p*_0_(*t*) and *p*_1_(*t*) always sum to one [46]).

#### The Crick switch

Crick’s original proposal involves proteins with two marking (e.g., phosphorylation) states, which we will call *unmarked* (−) and *marked* (+). He imagines that two such proteins can bind together to form a dimer with three possible states: *unmarked* (−−), *partially marked* (+−), and *fully marked* (++). (He assumes that the dimer exhibits two-fold rotational symmetry, so that the −+ state is equivalent to the +− state.) The proposal also supposes that there exists an enzyme that tends to quickly drive the system from the partially marked state into the fully marked state. The enzyme and additional state work together to stabilize the fully marked state, and hence extend the time scale on which a one-bit memory can be stored: if molecular turnover pushes the system into the partially marked state, the enzyme tends to ‘correct the error’ by pushing it back.

Following Crick, we define the **Crick switch** as a continuous-time Markov chain on three states. Like the two-state model, its behavior is characterized by time-dependent state occupancy probabilities ***p***(*t*) := (*p*_0_(*t*), *p*_1_(*t*), *p*_2_(*t*))^*T*^ and a master equation. Its Hamiltonian is defined as

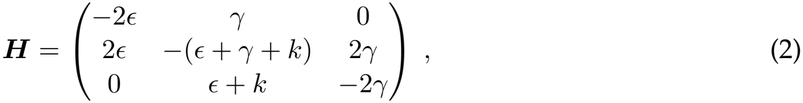

where *k* ≥ 0 is the error correction rate, and *ϵ* and *γ* are again the marking noise and turnover rate.

### Memory can be quantified in terms of mutual information

#### One-bit memory task

For simplicity and tractability, we will only study how well molecular memory switches store **one bit of information** regarding whether there *was* (*u* = 1) or *was not* (*u* = 0) a stimulus recently. In what follows, we remain agnostic about the nature of the stimulus and are mainly interested in determining how much longer the time scale of information loss is than the turnover time scale as a function of various choices (e.g., whether there is error correction). We particularly want to know whether error correction can produce the ten-to hundredfold time scale disparity that appears in our motivating biological examples. Because only relative time scales matter, we almost always plot functions of the system time *t* in units where *γ* = 1.

We assume the system encodes the presence or absence of the stimulus via its initial marking state: if there *was* a stimulus recently, we initialize it in its fully marked state; if there *was not*, we initialize it according to its *steady state distribution* ***π*** := lim_*t→∞*_ ***p***(*t*) (which generally corresponds to the unmarked state for the models we consider). This memory task is easy to understand for the two-state model. A recent stimulus (*u* = 1) initializes the switch in its marked state; if there was no stimulus (*u* = 0), or enough time has passed, the switch is usually unmarked (Fig. 2a).

**Figure 2.**
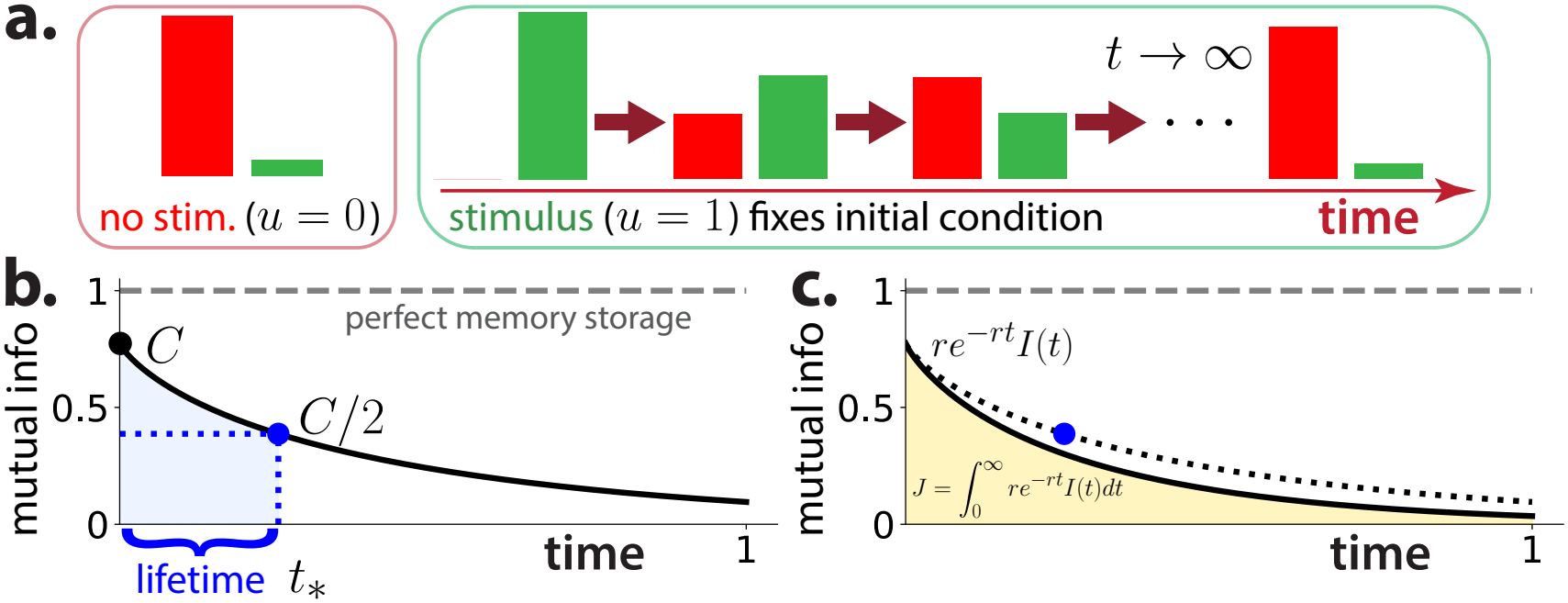
Memory can be quantified in terms of mutual information. (a) Red: unmarked probability *p*_0_(*t*); green: marked probability *p*_1_(*t*). Left: histogram visualization of two-state model probabilities given no stimulus (*u* = 0). Right: histograms over time given a stimulus (*u* = 1). (b) Mutual information *I*(*t*) vs time for the two-state model (*γ* = 1, *ϵ* = 0.1). It is bounded between zero (*no memory*) and one (*perfect memory*, dashed gray line). We define capacity as *C* := *I*(0) and lifetime as the time *t*_*\**_ at which *I*(*t*) equals half of its initial value. (c) Performance (*J*) is defined as the discounted area under *I*(*t*) (shaded yellow region). *r >* 0 is the discounting rate.

#### Memory lifetime, capacity, and performance

To ensure fair comparisons between different systems, we formally measure memory in terms of the mutual information (MI) [48, 49] *I*(*t*) between the binary stimulus *u* ∈ {0, 1} and the system’s state, assuming *u* = 0 and *u* = 1 are equally likely. MI has been used to understand memory in neural systems, and is appealing partly because it offers a general way of quantifying memory across many scales and substrates [50–52]. *I*(*t*) is bounded between zero (*no memory*) and one (*perfect memory*), and is a strictly decreasing function of time with lim_*t→∞*_ *I*(*t*) = 0 (Fig. 2b). To extract interpretable numbers from MI curves, we also define **memory lifetime** as the time at which *I*(*t*) equals half its original (*t* = 0) value (Fig. 2b, *t*_*\**_), which reflects the marked-to-unmarked first-passage time; **memory capacity** as the maximum value of *I*(*t*) (Fig. 2b, *C*), which reflects the false-positive probability; and **memory performance** as the (discounted) area under *I*(*t*) (Fig. 2c). This performance definition implies it is better to encode a stimulus accurately, and then suddenly forget it, than to poorly encode it for longer.

## Results

### Turnover determines memory of two-state model

We begin by analyzing the **two-state model**, which illustrates how memory is limited by turnover and marking noise in the absence of strategies like error correction. The two-state model’s behavior is simple and easy to mathematically characterize. When initialized in its marked state, in the absence of marking noise (*ϵ* = 0) it tends to enter its unmarked state on the time scale *τ* = 1*/γ* of turnover (Fig. 3a, b). One way to quantify this is by computing the average first-passage time associated with the marked-to-unmarked transition, which is precisely *t*_*F*_ = 1*/γ* (Fig. 3c). Its behavior is similar when there is marking noise (*ϵ* > 0), except that (i) there is a nonzero probability of a switch being marked, even in the long-time limit (Fig. 3d, e); and (ii) the time scale of the system’s approach to steady state equals *τ* = 1*/*(*γ* + *ϵ*), which depends on the marking noise level *ϵ*. Nonzero marking noise produces false positives (i.e., the switch may be marked without a stimulus), with the false-positive probability increasing as *ϵ/γ* increases (Fig. 3f).

**Figure 3.**
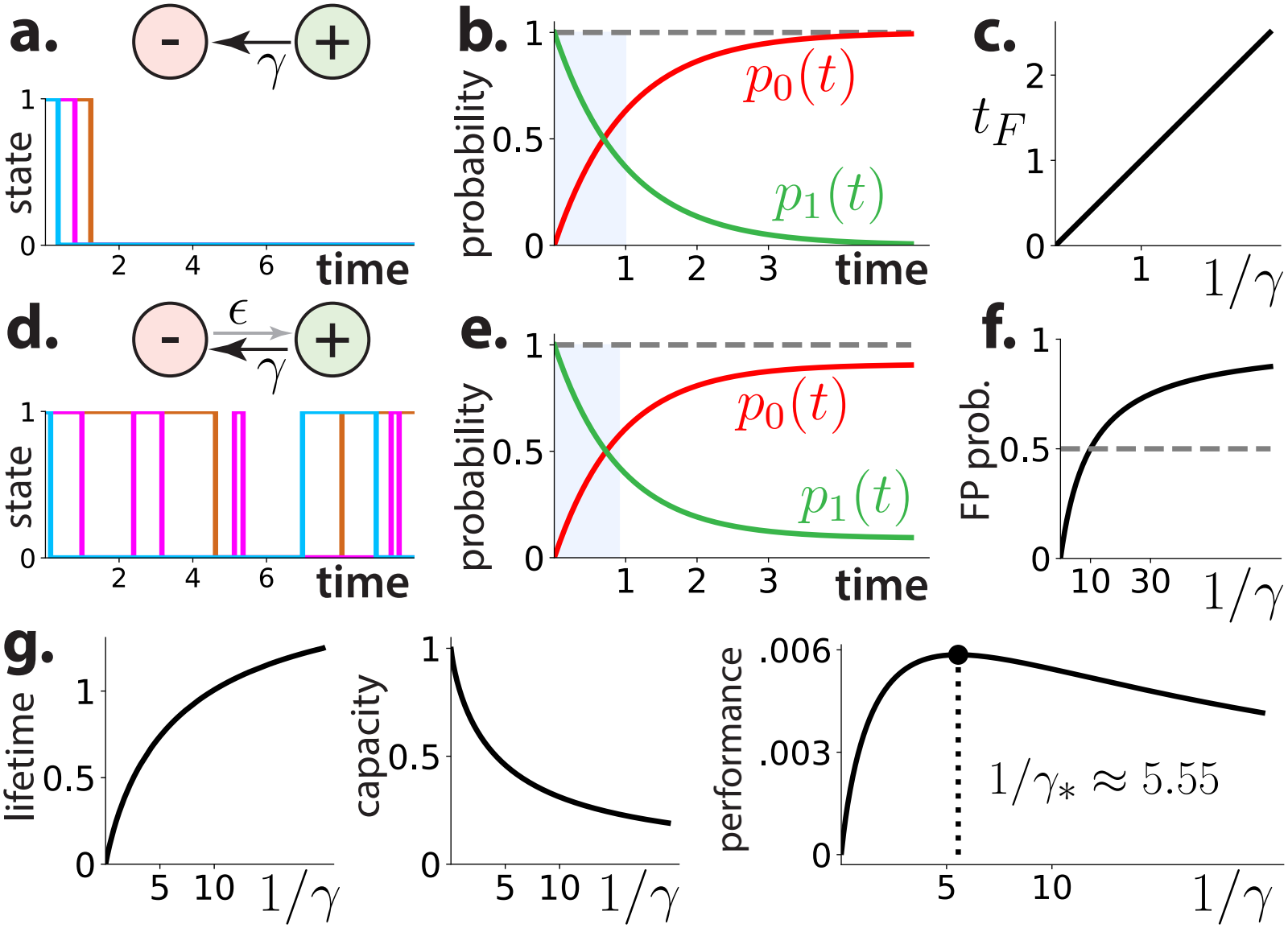
Turnover determines memory of two-state model. (a) Schematic of pure turnover model and three example stochastic simulations (*γ* = 1, *ϵ* = 0). (b) State occupancy probabilities over time (*γ* = 1, *ϵ* = 0). Shaded region: time before *τ* = 1*/γ*. (c) Average first-passage time vs 1*/γ*. (d) Schematic of the two-state model and three example stochastic simulations (*γ* = 1, *ϵ* = 0.1). (e) State occupancy probabilities over time (*γ* = 1, *ϵ* = 0.1). Shaded region: time before *τ* = 1*/*(*ϵ* + *γ*). (f) Average first-passage time 1*/γ* vs false-positive probability *ϵ/*(*ϵ* + *γ*) (*ϵ* = 0.1). Horizontal dashed line: *p*_FP_ = 0.5. (g) Lifetime, capacity, and performance vs 1*/γ* (*ϵ* = 0.1, *r* = 1*/*100).

We will find two types of analyses particularly useful in assessing how well switches like the two-state model store memories. The first is to analyze the eigenvalues and (right) eigenvectors of the Hamiltonian ***H***. The nonzero eigenvalues of ***H*** control how quickly the system equilibrates, and hence loses all information about its initial condition; the (right) eigenvectors describe the associated flow of probability. The two-state model’s Hamiltonian has two eigenvalues, *λ*_0_ = 0 and *λ*_1_ = −(*ϵ* + *γ*), and corresponding eigenvectors:

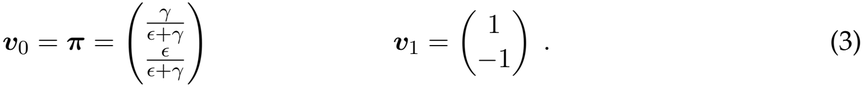

The first eigenvalue-eigenvector pair corresponds to the system’s steady state distribution, while the second determines both the manner and time scale of its relaxation to steady state. Since there are only two states, ***v***_1_ reflects a trivial fact: as the system relaxes, every unit of probability that leaves the marked state must enter the unmarked state. The overall solution to the master equation can be written in terms of these eigenvalues and eigenvectors:

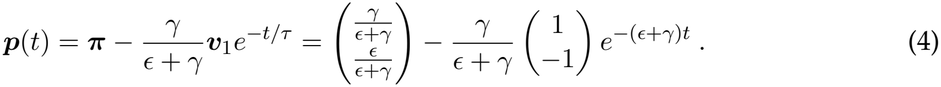

This picture is consistent with an information-theoretic analysis, which indicates that lifetime (which is related to the marked-to-unmarked first-passage time) increases with increasing 1*/γ* (Fig. 3g, left); capacity (which is related to the false-positive probability) is a decreasing function of 1*/γ* (Fig. 3g, center); and performance (which depends on both the memory lifetime and the false-positive probability) peaks at a specific value of *γ* (Fig. 3g, right).

### A pair of two-state switches slightly outperforms a single switch

We now turn to strategies for improving the two-state model. The most obvious is to redundantly encode information using multiple switches—two, in the simplest case—instead of one. Intuitively, one expects better performance because information may remain present at the population level even if any single switch loses information due to noise or turnover. Motivated by this idea, we define the **pair model** as two independent and identical two-state models; it can also be viewed as the Crick switch with no error correction (*k* = 0), as it is defined by the same Hamiltonian (Eq. 2). See Fig. 4a for a schematic and example stochastic simulations.

**Figure 4.**
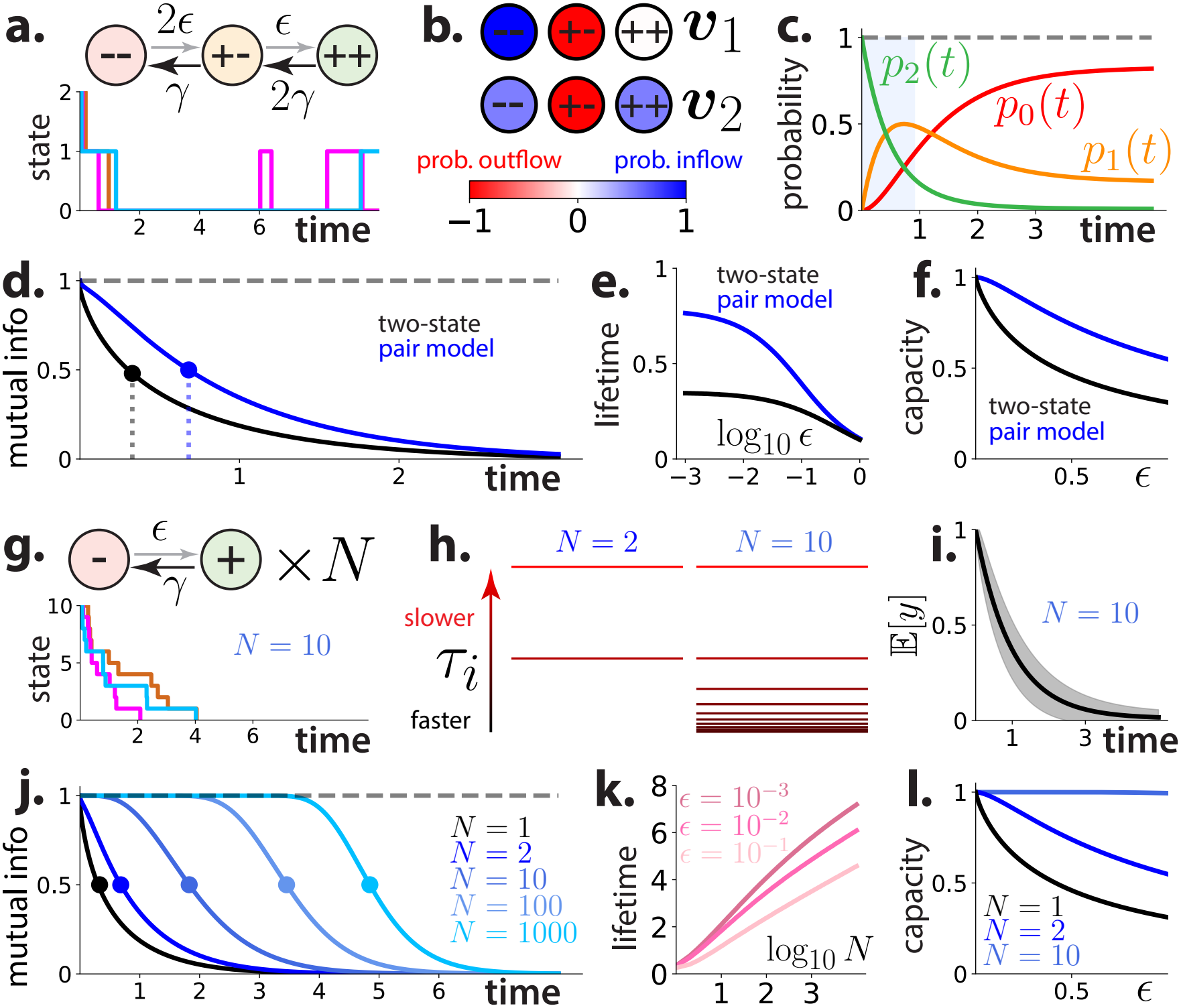
Increasing the number of switches inefficiently improves performance. Time is measured in units where *γ* = 1. (a) Schematic of the pair model and three example stochastic simulations (*ϵ* = 0.1). (b) First and second right eigenvectors of the pair model’s Hamiltonian, which indicate how probability flows through the system. Blue: probability inflow, red: probability out-flow. (c) State occupancy probabilities over time for the pair model (*ϵ* = 0.1). Shaded region: time before *τ* = 1*/*(*ϵ* + *γ*). (d) *I*(*t*) for the two-state (black) and pair model (blue) (*ϵ* = 0.01). Dots and dashed lines indicate lifetimes. (e) Lifetime vs log_10_ *ϵ* for two-state and pair model. (f) Capacity vs *ϵ* for two-state and pair model. (g) Schematic of the multi-switch model and three example stochastic simulations (*N* = 10, *ϵ* = 0.1). (h) Time scales *τ*_*i*_ = −1*/λ*_*i*_ of the multi-switch model for *N* = 2 and *N* = 10. As *N* increases, only faster time scales are added. (i) Mean fraction of switches that are marked over time (*N* = 10, *ϵ* = 0.01). Shaded region: *±*1 standard deviation. (j) *I*(*t*) for the multi-switch model for different values of *N* (*ϵ* = 0.01). (k) Lifetime vs log_10_ *N* for different values of *ϵ*. (l) Capacity vs *ϵ* for different values of *N*.

Its Hamiltonian has three eigenvalues: *λ*_0_ = 0, *λ*_1_ = −(*ϵ* + *γ*), and *λ*_2_ = −2(*ϵ* + *γ*). The first two eigenvalues are the same as those of the two-state model, and *λ*_2_ introduces a *faster* (rather than slower) new time scale. The three corresponding eigenvectors of ***H*** are

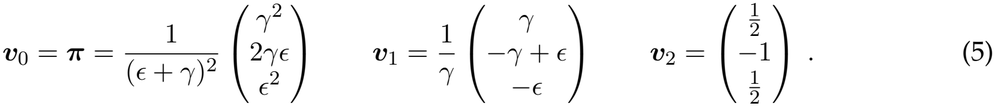

These eigenvectors indicate that probability moves through the system in two ways, at least assuming *γ* > *ϵ*: in the ***v***_1_ mode, it tends to move from the partially marked state to the unmarked state; and in the ***v***_2_ mode, it tends to move from the partially marked state to the unmarked and fully marked states (Fig. 4b). The solution to this model’s master equation is

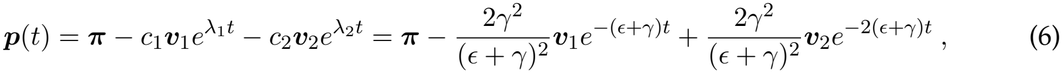

so ***p***(*t*) relaxes to its steady state on a time scale *τ* = −1*/λ*_1_ = 1*/*(*ϵ* + *γ*) (Fig. 4c). From an information perspective, there is no downside to adding a second switch: this model’s *I*(*t*) curve is consistently higher than that of the two-state model (Fig. 4d), and so are lifetime (Fig. 4e) and capacity (Fig. 4f). Still, because memory lifetime remains around the turnover time scale (Fig. 4d, vertical dashed lines), performance has not substantially improved beyond the two-state model.

### Increasing the number of switches inefficiently improves performance

We can generalize one or two identical and indistinguishable two-state models to *N* ≥ 1 such models (see Fig. 4g for a schematic and example stochastic simulations), which allows us to formally assess the population-averaging strategy mentioned in the introduction. Note that the meaning of *N* depends strongly on the specific biological context: for protein-marking-based switches like CaMKII, *N* corresponds to the number of proteins (although each protein may have multiple subunits); for histone marking, *N* corresponds to the number of approximately independent local chromatin domains; and for DNA methylation, *N* corresponds to the number of approximately independent CpG sequence contexts.

The *N*-switch model has *N* +1 states (zero switches marked, one is marked, …, all are marked), and its Hamiltonian is implicitly defined via the chemical master equation (CME) [53–55]:

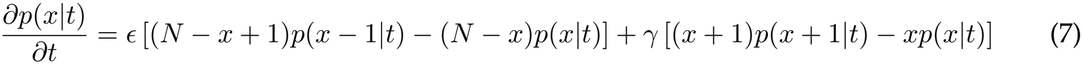

where *x* ∈ {0, 1, …, *N*} is the number of marked switches. We can exploit the relationship between this model and the two-state model (see *Supporting Information*) to find that its eigenvalues are

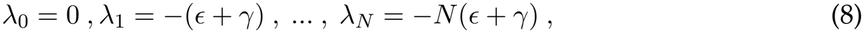

which implies that introducing additional switches yields only additional faster (rather than slower) time scales (Fig. 4h). To get intuition for how this model behaves, it is instructive to compute the expected fraction of marked switches over time (Fig. 4i):

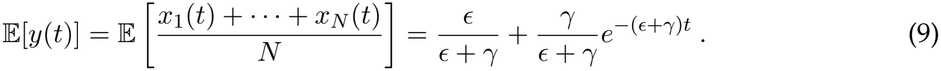

This quantity has exactly the same time course as the probability of the two-state model being marked (see Fig. 3e). What makes the multi-protein system different is that it has smaller fluctuations about this value; the variance of *y*(*t*) equals

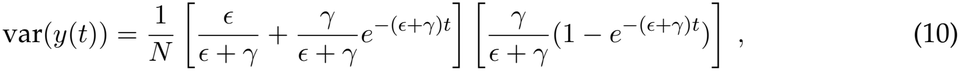

and scales like 1*/N*. There is no information-theoretic downside to making *N* larger: *I*(*t*) becomes uniformly higher, more sigmoidal-looking, and shifts to the right (Fig. 4j). These increases occur not because increasing *N* produces new, slower time scales, but because fluctuations about the average trajectory get smaller. As variance decreases, it becomes easier to distinguish the changing system from its steady state, for example by comparing Eq. 9 to its steady state value. On the other hand, increasing *N* by several orders of magnitude does not *substantially* improve the system’s memory storage capabilities, since lifetime depends logarithmically on *N* (Fig. 4k). Increasing *N* especially improves capacity, which becomes near-maximal at some *N* < 10 (Fig. 4l).

We conclude that the ‘population-averaging’ strategy is useful but does not greatly extend the lifetime of stored memories. For *ϵ* > 0.001 and *N* ≈ 10, 000, lifetime improves by less than an order of magnitude over the two-state model. Hence, redundancy alone cannot explain the memory-to-turnover ratios of *t*_*\**_*/τ* ~ 60 (CaMKII) and *t*_*\**_*/τ* ~ 10 − 100 (epigenetic marks) of our motivating examples. High *N* may also be unrealistic; there are only 80 CaMKII holoenzymes in a typical postsynaptic density, for example [56].

### Crick-like error correction produces a long time scale

We are now ready to study the **Crick switch**, which can be viewed as a generalization of the pair model that includes error correction (see Fig. 5a for a schematic and example stochastic simulations). Error correction tends to drive it from its partially marked state to its fully marked state, thereby ‘correcting’ putative errors (Fig. 5b). Its Hamiltonian (Eq. 2) has three eigenvalues: *λ*_0_ = 0, *λ*_1_ = −[*ϵ* + *γ* − Δ], and *λ*_2_ = −[2(*ϵ* + *γ*) + Δ + *k*], which depend on the *eigenvalue gap*

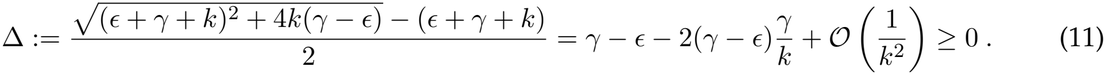

**Figure 5.**
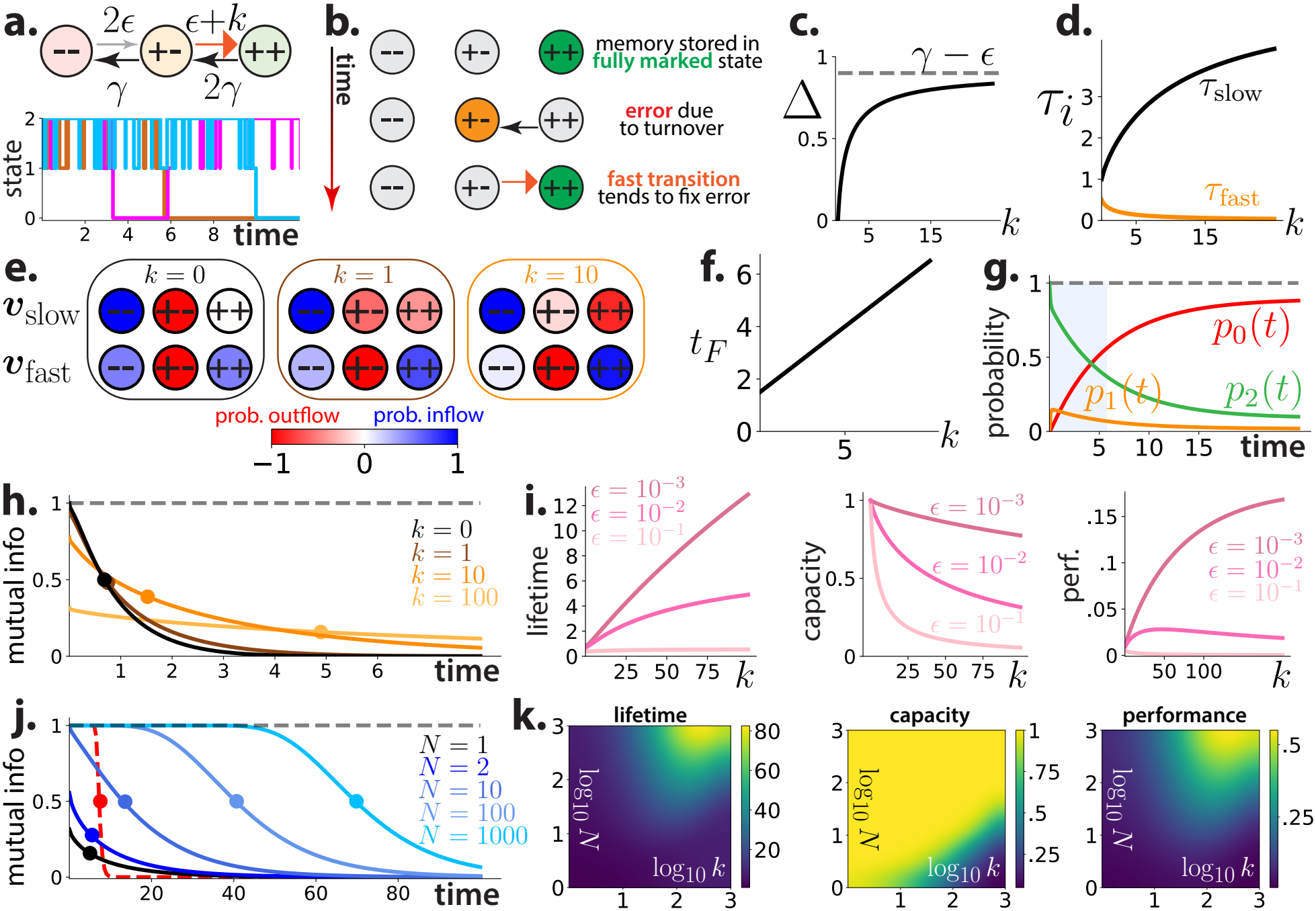
Error correction efficiently extends memory lifetime by introducing a slow time scale. Time is measured in units where *γ* = 1. (a) Schematic of Crick switch and three example stochastic simulations (*ϵ* = 0.01, *k* = 10). (b) When *k* ≫ *γ*, if turnover moves the system into its partially marked state, it tends to quickly transition back into the fully marked state. (c) Eigenvalue gap vs *k* (*ϵ* = 0.1). (d) Time scales *τ*_*i*_ = −1*/λ*_*i*_ vs *k* (*ϵ* = 0.1). (e) First (***v***_slow_) and second (***v***_fast_) right eigenvectors for different values of *k*. Blue: probability inflow, red: probability outflow. As *k* increases, ***v***_slow_ increasingly corresponds to probability moving from the fully marked to unmarked state (*memory loss*); ***v***_fast_ increasingly corresponds to probability moving from the partially marked to fully marked state (*error correction*). (f) Average first-passage time vs *k* (*ϵ* = 0.01). (g) State occupancy probabilities over time (*ϵ* = 0.01, *k* = 10). Shaded region: time before *τ*_slow_ = −1*/λ*_1_. (h) *I*(*t*) for different values of *k* (*ϵ* = 0.01). (i) Lifetime, capacity, and performance vs *k* for different values of the ambient marking rate *ϵ*. (j) *I*(*t*) for different values of *N* (*ϵ* = 0.01, *k* = 100). Red line: *I*(*t*) for *N* = 100, 000 two-state models for comparison. (k) Lifetime, capacity, and performance vs log_10_ *k* (horizontal) and log_10_ *N* (vertical) (*ϵ* = 0.01).

When *k* = 0, the gap equals zero, and the system reduces to the pair model. As *k* → ∞, Δ asymptotically approaches *γ* − *ϵ* (Fig. 5c). We call Δ a ‘gap’ because it controls the difference between the eigenvalues of the Crick switch and pair model; when *k* is large, the eigenvalues of the Crick switch become very different, since we approximately have *λ*_1_ ≈ 2*ϵ* and *λ*_2_ ≈ (3*γ* +*ϵ*+*k*). Equivalently, the eigenvalue difference |*λ*_2_ − 2*λ*_1_| = 3Δ + *k* depends strongly on Δ.

Importantly, unlike the *N* > 1 generalization of the two-state model considered thus far, this model involves a time scale *slower* than the turnover time scale *τ* = 1*/*(*ϵ* + *γ*): the *memory time scale τ*_slow_ := −1*/λ*_1_ = 1*/*(*ϵ* + *γ* − Δ). It becomes slower as *k* increases (Fig. 5d, black line), and

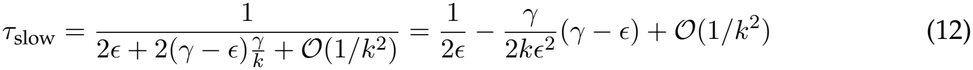

in the large *k* limit. Meanwhile, the model’s other time scale (the time scale of *error correction*) *τ*_fast_ := −1*/λ*_2_ = 1*/*[2(*ϵ* + *γ*) + Δ + *k*] grows faster as *k* increases (Fig. 5d, orange line), since

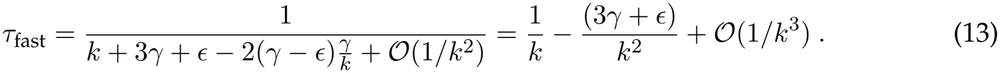

The eigenvectors of ***H*** provide a more precise picture of how these two time scales relate to the system’s dynamics. The three right eigenvectors of ***H*** are

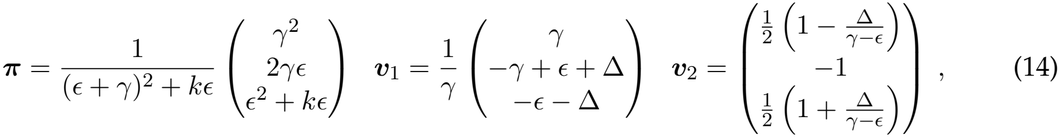

with ***v***_slow_ := ***v***_1_ corresponding to *τ*_slow_ and ***v***_fast_ := ***v***_2_ corresponding to *τ*_fast_. As *k* becomes larger, the eigenvector ***v***_slow_ increasingly looks like it represents probability flowing directly from the fully marked state to the unmarked state, and hence corresponds to *memory loss* (Fig. 5e, top). The eigenvector ***v***_fast_ increasingly looks like it represents probability flowing from the partially marked state to the fully marked state, and hence corresponds to *error correction* (Fig. 5e, bottom). This intuition becomes exactly correct in the infinite *k* limit, since

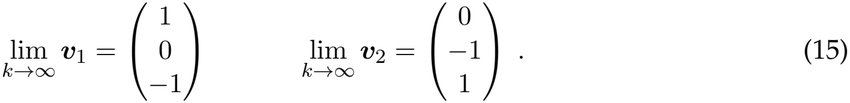

Another way to quantitatively understand error correction is to compute how long it takes until the first transition between the fully marked state and the unmarked state. The corresponding first-passage time distribution is a mixture of exponentials, and its mean and variance are

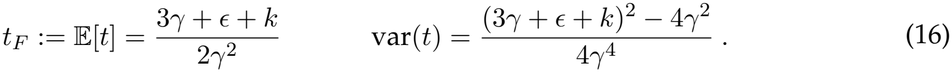

This means that the mean first-passage time (or equivalently, time until a ‘false negative’) can be arbitrarily large as long as *k* is large enough (Fig. 5f). Hence, the error correction mechanism can in this sense be arbitrarily effective. But increasing *k* also increases the false-positive probability

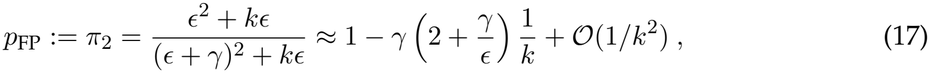

which leads to a tradeoff between capacity and lifetime. This increase in false positives is the major drawback of Crick-like error correction: noise-induced transitions from the unmarked to fully marked state become significantly more likely. It is interesting to note that it is only a problem if there is ambient marking noise (i.e., *ϵ* > 0). If *ϵ* = 0, there are no false positives because it is impossible to reach the fully marked state from the unmarked state.

When *k* is large enough to make *τ*_slow_ substantially different from *τ*, but not so large that the false-positive probability is high, the Crick switch functions like a two-state model with a smaller *effective* turnover rate. The visual similarity between the Crick switch’s state occupancy probabilities (Fig. 5g) and the two-state model’s (Fig. 3e) in this regime illustrates this.

### Moderate error correction greatly improves performance

Since increasing *k* both lengthens a memory-related time scale and increases the false-positive probability, we expect a moderate value of *k* to be ideal for memory storage. Indeed, a moderately large *k* pushes the curve to the right, and a *k* which is too large yields a very low capacity (Fig. 5h). Across a range of marking noise levels, increasing *k* improves lifetime but reduces capacity (Fig. 5i). Generally, performance is maximized at a moderate value of *k*, but when marking noise is high, it may be maximized when *k* = 0 (Fig. 5i, right). Note that increasing *k* is different from increasing *N*, which *always* improves performance since *I*(*t*) becomes larger for every *t* > 0.

Crick-like error correction improves lifetime by around tenfold or less and achieves a performance level similar to the population-averaging strategy using many fewer switches. In that sense it is much more resource-efficient. Combining both strategies by using a population of *N* > 1 independent and indistinguishable Crick switches yields MI curves which are strikingly better than those of either strategy alone (Fig. 5j). The midpoint of these curves still depends logarithmically on *N*, but for *N* = 1000 they occur at around *t*_*\**_ ≈ 80 rather than around *t*_*\**_ ≈ 5. Lifetime and performance both increase with *N* but peak around *k* ≈ 150. Capacity remains near one as long as log_10_ *N* ≳ log_10_ *k* − 1 (Fig. 5k).

We conclude that combining population-averaging with Crick-like error correction yields memory lifetimes almost *two* orders of magnitude longer than the turnover time scale, which is consistent with *t*_*\**_*/τ* ~ 60 (CaMKII) and *t*_*\**_*/τ* ~ 10 − 100 (epigenetic marks). Hence, our model supports the idea that error correction and redundancy are both required for CaMKII and epigenetic marks to store memories significantly beyond the time scale of turnover.

### Error correction also improves performance of more realistic switches

Next, we show that the behavior of the Crick switch does not substantially change when we make it more realistic. First, in line with Crick’s original proposal, we explicitly model both monomers and dimers: two monomers can bind to one another to form a dimer, and a dimer can spontaneously break apart to form two monomers. Both monomers and dimers can be marked or unmarked, with the parameters *ϵ, γ*, and *k* having the same meaning as before (Fig. 6a, right). Second, we will assume that there are fluctuations in the number of available monomers and dimers. We model this by including a ‘reservoir’ species: monomers can spontaneously convert to this species, and this species can convert to an unmarked monomer (Fig. 6a, left).

**Figure 6.**
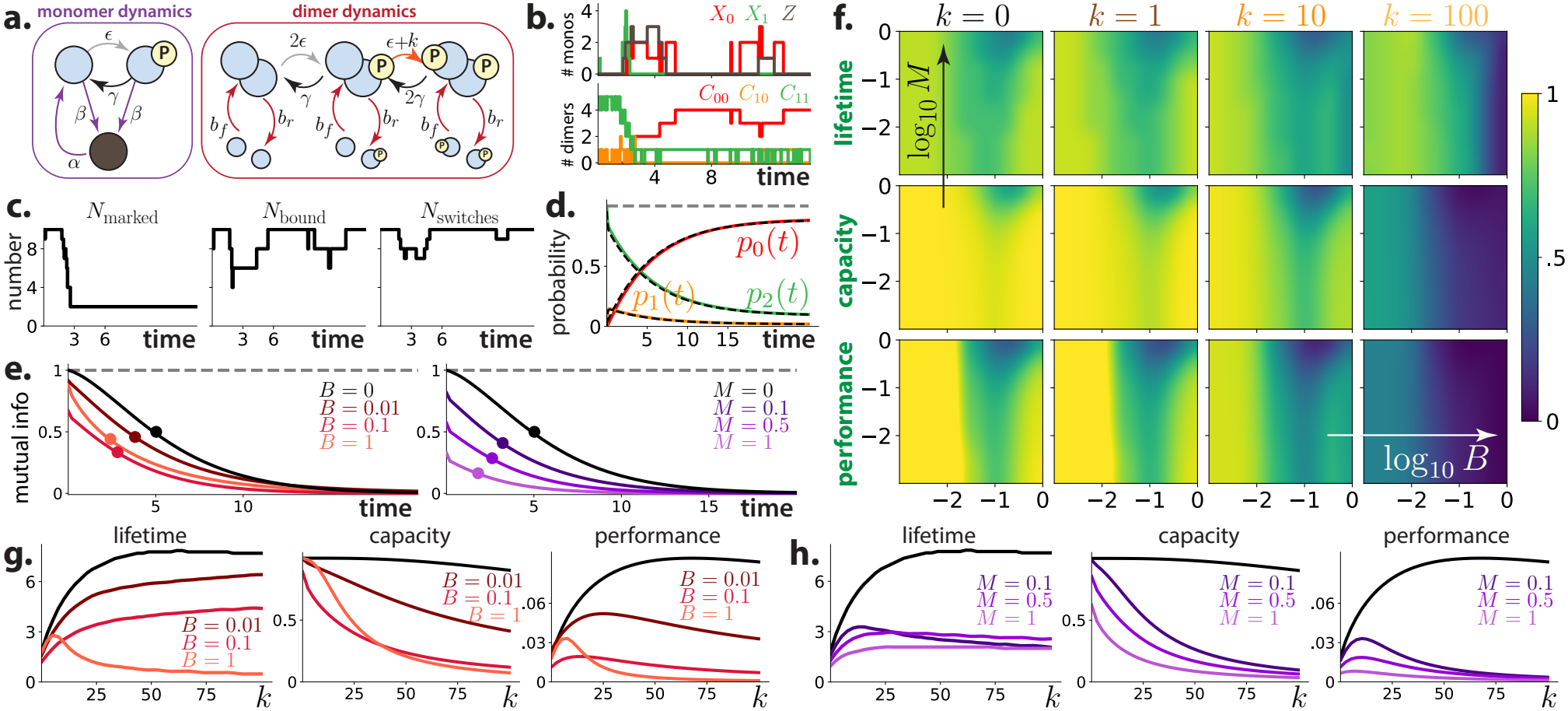
Error correction also improves more realistic switches. This model has six species: unmarked (*X*_0_) and marked (*X*_1_) monomers, a reservoir species *Z*, and dimers with three different marking states (*C*_00_, *C*_10_, *C*_11_). (a) Schematic of reaction list. Left: monomer dynamics, which includes conversion between ‘active’ and ‘reservoir’ molecules. Right: dimer dynamics, which includes binding and unbinding. (b) Example stochastic simulation (*γ* = 1, *ϵ* = 0.01, *k* = 10, *b*_*f*_ = 1, *b*_*r*_ = 0.1, *α* = 0.7, *β* = 0.3, *N* = 10). Top: monomer counts over time. Bottom: dimer counts over time. (c) Number of marked molecules over time, number of bound molecules over time, and number of non-reservoir molecules over time. Trajectory data same as in (b). (d) Probabilities of realistic model (black dashed lines) reduce to those of the Crick switch when *b*_*f*_ ≫ *b*_*r*_, *α* ≫ *β*, and *b*_*f*_ and *α* are very small. (e) Left: *I*(*t*) for different values of *B*. Right: *I*(*t*) for different values of *M*. (f) Lifetime, capacity, and performance as functions of log_10_ *B* (left to right) and log_10_ *M* (bottom to top) for different values of *k*. All values are normalized relative to their Crick switch value, so color intensity quantifies loss of fidelity due to new model elements (binding and unbinding, monomer fluctuations). (g) Lifetime, capacity, and performance vs *k* for different values of *B* (*M* = 0.1). Black: curves for same number of Crick switches. (h) Lifetime, capacity, and performance vs *k* for different values of *M* (*B* = 0.5). Black: curves for same number of Crick switches.

In a typical stochastic simulation of this system, the counts of all of model species fluctuate (Fig. 6b), and the number of bound and non-reservoir molecules also fluctuate (Fig. 6c). But this model reduces to *N* Crick switches in the limit that the binding rate is much larger than the unbinding rate (*b*_*f*_ ≫ *b*_*r*_), the monomer gain rate is much larger than the monomer loss rate (*α* ≫ *β*), and each of those parameters is small (Fig. 6d).

Let 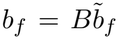 and 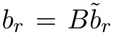, so that *B* is the time scale of binding and unbinding dynamics. Similarly, let 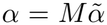 and 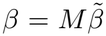, so that *M* is the time scale of monomer gain and loss. Faster binding and unbinding dynamics generally reduces available stimulus information (Fig. 6e, left), albeit not necessarily in a monotonic fashion (compare *B* = 0.1 and *B* = 1). This occurs because the error correction mechanism, which only works for dimers, becomes less effective when the dimer population size fluctuates. It fluctuates most for intermediate *B* (here, for *B* ≈ 0.1); small *B* yields stable single dimers, while large *B* yields fast equilibration of the monomer and dimer populations. Fast fluctuations in monomer number also reduce information (Fig. 6e, right).

Lifetime, capacity, and performance are more sensitive to *B* than *M* (Fig. 6f; note that the colorbar indicates fraction of the corresponding Crick switch value). For moderate values of *k*, lifetime becomes less than around half of its Crick switch value when *B* ≳ 0.1. Capacity and performance become less than half of their Crick switch values earlier, when *B* ≳ 0.01 (Fig. 6f, g). Changing *M* has a noticeable effect, with all quantities much less than their Crick switch values, only if *B* ≳ 0.5 (Fig. 6f, h). One consequence of this model’s additional fluctuations is that the optimal value of *k* is substantially lower than for the Crick switch (Fig. 6g, h). Moderate amounts of error correction still improve lifetime and performance while reducing capacity.

## Discussion

Our analysis of a minimal model of molecular error correction has shown that error correction improves memory by producing a slower time scale, has the downside of increasing false positives if there is ambient marking noise, and is significantly more resource-efficient than simply using a large number of switches. Moreover, our model quantitatively connects low-level rates (e.g., the error correction rate *k*) to observable time scales of memory (*t*_*\**_) and error correction (*τ*_fast_); we find that moderate error correction generally produces a memory time scale between one and two orders of magnitude longer than the time scale of turnover, which is consistent with measurements of CaMKII (*t*_*\**_*/τ* ~ 60) and epigenetic marks (*t*_*\**_*/τ* ~ 10 − 100). It is interesting that Crick’s proposal may provide a good model of molecular error correction even though it has largely failed as a model of synaptic memory [11].

### Theoretical contributions

The first major difference between our model of molecular error correction and others is its relative simplicity—each switch has only three states, and there are only three types of transitions (turnover, erroneous marking due to marking noise, and error correction). The abstract character of our model allows us to connect it to a variety of concrete examples of molecular error correction, and it also allows us to analytically compute many quantities of interest, including eigenvalues, transition probabilities, and first-passage times. Most existing models involve large numbers of states, species, and reactions, and their behavior can only be understood through simulations or by making strong approximations. This is true of recent models of CaMKII dynamics (~ 18^12^ states and ~40 reactions) [57] and epigenetic dynamics (7 species and 52 reactions) [58].

The second major difference is that our model accounts for intrinsic noise: state space trajectories are stochastic rather than deterministic. Classical models of CaMKII autophosphorylation [14] and epigenetic mark dynamics [59] were usually formulated in terms of ODEs, which involve no noise. (Although the master equation is a system of ODEs, and hence defines deterministic dynamics, it characterizes how the *distribution* over states evolves rather than how states evolve. A model which is deterministic at the level of trajectories would follow, e.g., ODEs like reaction rate equations [55].) Even modern models tend to incorporate noise in only some analyses [57, 58, 60] rather than treating it as a fundamental feature of the system. This is conceptually important because our conclusions change without it: more error correction would always be better in the absence of ambient marking noise (*ϵ* = 0), and even a two-state switch could store information for arbitrarily long in the absence of all noise (e.g., if we modeled it using reaction rate equations).

Although our specific formalization and analysis of molecular error correction appear to be new, there are interesting parallels between our work and other theoretical studies. Bruno et al.’s analysis of a model of chromatin modification dynamics [58] is particularly related to ours, as it indicates that robust memory requires a separation of time scales and an associated error-correction-like motif. In that work and follow-up work [60], they assume that the transitions from unmodified to modified chromatin tend to be much faster than the reverse transitions. But the specific transition topology they consider is different than that of the Crick switch in its details and tends to produce bistability rather than slow transitions out of the fully marked state.

Real systems do not necessarily remember all stimuli, or all aspects of all stimuli—real stimuli are multi-dimensional and vary nontrivially in time—and it may be preferable to forget a stimulus rather than retain information about it for arbitrarily long. Even if storing a long-lived memory is desirable, our MI analysis only quantifies how much information about the stimulus is *in principle* present, rather than how much information is accessible to a biologically plausible decoder. Also, as is well-known in other contexts, additional considerations become relevant when one would like information to be efficiently retrievable [61]. Finally, in this work we only consider a single type of stimulus which only ever appears once. Real memory storage devices are more useful when they can store many memories simultaneously, which leads to concerns regarding interference between memories [62, 63].

### Biological implications

When our error correction model is valid, one can use it to infer low-level reaction rates (e.g., the error correction rate *k*) from observable time scales of memory (*t*_*\**_) and turnover (*τ*). In cases where the biophysics of error correction are unknown or unclear, this connection may help generate plausible hypotheses about the underlying mechanism, since a possible mechanism’s rate may be inconsistent with what the model predicts given a known *τ* and *t*_*\**_. Consider, for example, the case of CaMKII. We observed earlier that *τ* ~ 1 s and *t*_*\**_*/τ* ~ 60 [3, 26, 27] within a postsynaptic density with *N* ~ 80 holoenzymes [56]. According to a numerical calculation based on our model, these numbers imply that the effective error correction rate should be *k* ~ 40 s^−1^ assuming a low ambient marking rate *ϵ/γ* = 0.001 (see Fig. 5k for a related heatmap). Interestingly, this is higher than the rate of T286 autophosphorylation under strongly activating conditions (1 − 10 s^−1^ [28, 29]). This discrepancy suggests that additional biological mechanisms may contribute to stabilizing CaMKII activity—by increasing the error correction rate *k*, decreasing the turnover rate *γ*, or through processes not captured by our minimal model. This lends credence to work that suggests other features of CaMKII dynamics, including CaMKII subunit exchange [64, 65], the ability for one holoenzyme to phosphorylate another without subunit exchange [66], and interactions with binding partners that compete with autoinhibition [67], may be critical for information retention. For histone marking, given moderate redundancy (*N* ~ 10) and a low ambient marking rate (*ϵ/γ* = 0.001), the estimates *τ* ~ 1 day [34] and *t*_*\**_ ~ 8 days [35] imply that *k/γ* ~ 5, which is consistent with the observed *τ*_fast_ ~ 2 − 6 h. For DNA methylation, similarly low rates of erroneous methylation, together with modest redundancy, can account for memory lifetimes of several weeks given fast remethylation kinetics (*τ*_fast_ ~ 0.3−4 h [36, 37]). For example, taking *τ* ~ 1 day, *ϵ/γ* = 0.001, *τ*_fast_ ~ 0.3 h, and *N* = 5 yields *t*_*\**_ ~ 35 days. In both cases, our model appears broadly consistent with suspected error correction mechanisms. However, it may be important to invoke other biology when marks are treated less abstractly. Owen et al., for example, find that histone mark fidelity depends on chromatin’s three-dimensional structure when mark locations are explicitly modeled [68].

Our error correction model should approximate the dynamics of molecules like CaMKII when they can be treated as having three biologically relevant macrostates (*unmarked, partially marked*, and *fully marked*), (ii) transitions between these macrostates are approximately Markovian, and (iii) individual ‘switches’ do not interact much. These assumptions, while crude, help us analyze error correction and let us make predictions about biologically distinct systems. It is possible to extend our work in a variety of ways, including by adding biological detail, allowing different switches to interact, and considering more complex storage and retrieval tasks. A potential extension we find particularly interesting is to relax our assumption that a single switch operates in the vicinity of one time scale (i.e., around *τ*), since real memory storage involves many time scales. Following Fusi et al.’s ideas about multiscale synaptic memory [69, 70], one way to model this would be to hierarchically connect many Crick-like switches with different intrinsic time scales.

Although it is increasingly accepted that learning and memory are possible even in organisms without nervous systems [71–74], like the single-celled ciliates *Paramecium* [75–77] and *Stentor* [78– 83], the molecular mechanisms that support these capabilities remain mysterious. The problem is arguably not a shortage of candidates—possibilities include information storage via RNA [84–86], post-translational modifications of proteins [1, 14, 87], and epigenetic modifications [33, 88, 89]— but rather that it is unclear when and why a given candidate might be preferred over others. In the absence of experiments that decisively support one or more candidates, theorizing about ‘design principles’ of molecular memory may provide insight regarding what future experiments ought to look for. Our analysis suggests that molecular error correction is one such principle.

## Materials and Methods

### Models and their exact solutions

Throughout this paper, we model molecular memory switches as continuous-time Markov chains with finitely many (*S* ≥ 2) states. Because these models evolve in time according to a master equation, they are completely defined by the choice of an *S* × *S* Hamiltonian matrix ***H***. See the *Supporting Information, Relevant facts about continuous-time Markov chains* for more discussion of this fact and its consequences.

We consider five models in this paper—the two-state model, the Crick switch, *N* two-state models, *N* Crick switches, and the realistic Crick-like switch—and exactly solve the master equations associated with all models except the realistic model. These results were used to efficiently generate mutual information curves and associated quantities (e.g., lifetime, capacity, and performance) without stochastic simulations. See the *Supporting Information* for detailed derivations of the solution of each model, including a characterization of their eigenvalues and eigenvectors. See the *Supporting Information, Realistic Crick-like switch* for the full chemical reaction list of the realistic model, which is depicted schematically in Fig. 6a.

### Numerical methods

We used an implementation of the Gillespie algorithm [53, 90] to simulate various models (e.g., in Fig. 3a), although stochastic simulations were only used for visualization, and were not used to generate any mutual information curves or associated quantities. To numerically solve the realistic model, which is analytically intractable, we solved the master equation by sparse matrix exponentiation. This approach is similar to finite state projection [91, 92], but no state space truncation was required since the realistic model has finitely many states.

## Data availability statement

Code that can be used to reproduce all figures is available on GitHub (https://github.com/john-vastola/crick-switch-model-26). It is written in Python and only uses standard libraries (e.g., NumPy, SciPy, Matplotlib). Detailed mathematical derivations of the solutions of all but the realistic model are in the *Supporting Information*.

## Acknowledgments

This work was supported by the Air Force Office of Scientific Research (FA9550-22-1-0345), the Multi-disciplinary University Research Initiative (MURI) Award by the Army Research Office (W911NF-21-1-0328), and a Schmidt Sciences Polymath Award.

