## Supporting Information for "Storing long-lived memories via molecular error correction: a minimal mathematical model of Crick’s memory switch"

**9 Realistic Crick-like switch** **47**

#### 1 Relevant facts about continuous-time Markov chains

##### 1.1 Motivation

In this paper, we model memory switches as continuous-time Markov processes with finitely many states. We do this for a few reasons. We would like models which have different states, like 'unmarked' and 'marked', and we would also like them to randomly transition between their various states, since we require some mechanism by which information can be lost. While we could have considered models where these transitions happen after a fixed amount of time, deterministically—for example, we could model a protein as losing a phosphate group after exactly one hour—this seems somewhat unrealistic. Models where processes like protein degradation happen randomly have, especially over the past decade, been shown to quantitatively describe available single-cell data well [1–3].

We assume models with finitely many states, as opposed to more complicated models with countably or uncountably infinite state spaces (e.g., protein concentrations), since this class of models is highly amenable to detailed mathematical analysis and allows us to treat Crick-like molecular error correction in detail. As we show towards the end of the main text, one obtains qualitatively similar results by considering a somewhat more biophysically realistic model of Crick's proposal.

##### 1.2 Definitions

A continuous-time Markov chain with $2 \leq S < \infty$ states is defined by a **master equation**

$$\frac{d}{dt}\boldsymbol{p}(t) = \boldsymbol{H}\boldsymbol{p}(t) \tag{1}$$

where $\boldsymbol{p}(t) = (p_0(t), ..., p_{S-1}(t))^T$ is a vector containing the probabilities $p_i(t)$ that the system occupies state $i \in \{0, ..., S-1\}$ at time $t$, and $\boldsymbol{H}$ is an $S \times S$ matrix that we call the **Hamiltonian**. The full behavior of the system is determined by an initial condition $\boldsymbol{p}(0)$, plus the dynamics defined by Eq. 1. Assuming $\boldsymbol{H}$ is time-independent, the solution of Eq. 1 is

$$\boldsymbol{p}(t) = e^{\boldsymbol{H}t}\boldsymbol{p}(0) \ . \tag{2}$$

The initial condition is assumed to be normalized, so that $\sum_{i=0}^{S-1} p_i(0) = 1$. For $\boldsymbol{H}$ to define a valid stochastic process, it must be **infinitesimal stochastic**, which means that $\mathbf{1}^T\boldsymbol{H} = \mathbf{0}^T$ and that $H_{ij} \geq 0$ when $i \neq j$ [4]. The first condition means that the columns of $\boldsymbol{H}$ sum to zero, i.e., $\sum_i H_{ij} = 0$ for all $j$. This condition is important, since it means $\boldsymbol{p}(t)$ is normalized for all $t > 0$ whenever $\boldsymbol{p}(0)$ is normalized. This follows from the observation that

$$\frac{d}{dt}\left[\mathbf{1}^T\boldsymbol{p}(t)\right] = \mathbf{1}^T\frac{d}{dt}\boldsymbol{p}(t) = \mathbf{1}^T\boldsymbol{H}\boldsymbol{p}(t) = \mathbf{0}^T\boldsymbol{p}(t) = 0 \ . \tag{3}$$

##### 1.3 Eigenvalues and eigenvectors of infinitesimal stochastic matrices

Since each stochastic process of interest is defined by a specific choice of $\boldsymbol{H}$, it is perhaps unsurprising that the spectral structure of $\boldsymbol{H}$ characterizes how probabilities evolve in time.

For us, the most important result is the **Perron-Frobenius theorem**, which says that if $\boldsymbol{H}$ is infinitesimal stochastic and *irreducible*:

- The eigenvalue zero has multiplicity one, and the unique (up to scaling) eigenvector $\boldsymbol{\pi}$ corresponding to it can be chosen to have nonnegative real entries. In particular, it can be chosen so that $\pi_i \geq 0$ for all states $i$, and $\sum_{i=0}^{S-1} \pi_i = 1$.
- The other eigenvalues of $\boldsymbol{H}$ have a negative real part.

Hence, if $\boldsymbol{H}$ is diagonalizable, the formal solution of the master equation (Eq. 2) can be written

$$\boldsymbol{p}(t) = \boldsymbol{\pi} - \sum_{i=1}^{S-1} c_i \ \boldsymbol{v}_i \ e^{\lambda_i t} = \boldsymbol{\pi} + \sum_{i=1}^{S-1} [\boldsymbol{w}_i^T \boldsymbol{p}(0)] \ \boldsymbol{v}_i \ e^{\lambda_i t} \tag{4}$$

where $\lambda_1, ..., \lambda_{S-1}$ are the nonzero eigenvalues of $\boldsymbol{H}$, the $\boldsymbol{v}_i$ are the corresponding right eigenvectors, the $\boldsymbol{w}_i$ are the corresponding left eigenvectors, and the $c_i$ are coefficients. Here, we have used the convention that the left eigenvectors are normalized so that $\boldsymbol{w}_i^T \boldsymbol{v}_j = \delta_{ij}$. Since each nonzero eigenvalue has a negative real part, this expression makes it obvious that

$$\lim_{t \to \infty} \boldsymbol{p}(t) = \boldsymbol{\pi} \ , \tag{5}$$

i.e., that $\boldsymbol{\pi}$ corresponds to the unique steady state distribution of the system. Incidentally, the assumption of the Perron-Frobenius theorem—that $\boldsymbol{H}$ is irreducible—is crucial to this result. If our system consisted of at least two disconnected parts, for example, there is not a unique steady state distribution.

A useful fact that we will use throughout these appendices is that the components of right eigenvectors that correspond to nonzero eigenvalues sum to zero. To see this, let $\boldsymbol{v}$ be a right eigenvector corresponding to a nonzero eigenvalue $\lambda$. Then

$$0 = \boldsymbol{0}^T \boldsymbol{v} = \boldsymbol{1}^T \boldsymbol{H} \boldsymbol{v} = \lambda \boldsymbol{1}^T \boldsymbol{v} \quad \Longrightarrow \quad \boldsymbol{1}^T \boldsymbol{v} = 0 \ . \tag{6}$$

##### 1.4 Eigenvector normalization convention

Eigenvectors are only defined up to an overall normalization factor. Throughout, we will consider a normalization which allows us to better interpret how (right) eigenvectors reflect probability flow dynamics: we will use the convention that, if $\boldsymbol{v}_i$ is a right eigenvector of $\boldsymbol{H}$, $\boldsymbol{v}_i$ is normalized so that $\|\boldsymbol{v}_i\|_1 := \sum_{k=0}^{S-1} |v_{ik}| = 2$ (in words: its 1-norm equals two).

Since (by Eq. 6) $\sum_k v_{ik} = 0$, $\boldsymbol{v}_i$ reflects the movement of probability rather than a distribution in its own right. By choosing $\|\boldsymbol{v}_i\|_1 = 2$, we can interpret it as indicating how exactly one unit of probability mass is lost by some set of states, and gained by some other set of states.

We will normalize all left eigenvectors $\boldsymbol{w}_i$ so that $\boldsymbol{w}_i^T \boldsymbol{v}_i = 1$.

##### 1.5 Detailed balance

In our setting, a system satisfies **detailed balance** if

$$H_{ij}\pi_j = H_{ji}\pi_i \tag{7}$$

for all $i \neq j$. Equivalently, a system satisfying detailed balance has

$$\boldsymbol{HD} = \boldsymbol{DH}^T \tag{8}$$

where $\boldsymbol{D}$ denotes the diagonal matrix whose $(i,i)$ entry is $\pi_i$. For us, the significance of detailed balance is that it implies a precise relationship between the right and left eigenvectors of $\boldsymbol{H}$, and in particular implies that we know the left eigenvectors of $\boldsymbol{H}$ given its right eigenvectors.

To see why this is true, assume $\pi_i > 0$ for all $i$ and consider the matrix $\boldsymbol{S} := \boldsymbol{D}^{-1/2}\boldsymbol{H}\boldsymbol{D}^{1/2}$. This matrix is symmetric, since

$$\boldsymbol{S} = \boldsymbol{D}^{-1/2}\boldsymbol{HD}\boldsymbol{D}^{-1/2} = \boldsymbol{D}^{-1/2}\boldsymbol{DH}^T\boldsymbol{D}^{-1/2} = \boldsymbol{D}^{1/2}\boldsymbol{H}^T\boldsymbol{D}^{-1/2} = \boldsymbol{S}^T \ . \tag{9}$$

This implies that $\boldsymbol{S}$ can be diagonalized by an orthogonal matrix, and in particular that $\boldsymbol{S} = \sum_{i=1}^{S} \lambda_i \tilde{\boldsymbol{v}}_i \tilde{\boldsymbol{v}}_i^T$ for eigenvectors $\tilde{\boldsymbol{v}}_i$ with $\tilde{\boldsymbol{v}}_i^T \tilde{\boldsymbol{v}}_j = \delta_{ij}$.

The result we care about follows from the fact that eigenvectors of $\boldsymbol{S}$ correspond to eigenvectors of $\boldsymbol{H}$. In particular, if $\tilde{\boldsymbol{v}}$ is an eigenvector of $\boldsymbol{S}$ with eigenvalue $\lambda$, then

- $\boldsymbol{v} = \boldsymbol{D}^{1/2}\tilde{\boldsymbol{v}}$ is a right eigenvector of $\boldsymbol{H}$ with eigenvalue $\lambda$, and
- $\boldsymbol{w} = \boldsymbol{D}^{-1/2}\tilde{\boldsymbol{v}}$ is a left eigenvector of $\boldsymbol{H}$ with eigenvalue $\lambda$.

The proof is easy:

$$\begin{aligned} \boldsymbol{Hv} &= (\boldsymbol{D}^{1/2}\boldsymbol{S}\boldsymbol{D}^{-1/2})(\boldsymbol{D}^{1/2}\tilde{\boldsymbol{v}}) = \boldsymbol{D}^{1/2}\boldsymbol{S}\tilde{\boldsymbol{v}} = \lambda\boldsymbol{D}^{1/2}\tilde{\boldsymbol{v}} = \lambda\boldsymbol{v} \\ \boldsymbol{H}^T\boldsymbol{w} &= (\boldsymbol{D}^{-1/2}\boldsymbol{S}\boldsymbol{D}^{1/2})(\boldsymbol{D}^{-1/2}\tilde{\boldsymbol{v}}) = \boldsymbol{D}^{-1/2}\boldsymbol{S}\tilde{\boldsymbol{v}} = \lambda\boldsymbol{D}^{-1/2}\tilde{\boldsymbol{v}} = \lambda\boldsymbol{w} \ . \end{aligned} \tag{10}$$

It is also true that left and right eigenvectors defined in this way comprise a biorthogonal set, since

$$\boldsymbol{w}_i^T\boldsymbol{v}_j = \tilde{\boldsymbol{v}}_i^T\boldsymbol{D}^{-1/2}\boldsymbol{D}^{1/2}\tilde{\boldsymbol{v}}_j = \tilde{\boldsymbol{v}}_i^T\tilde{\boldsymbol{v}}_j = \delta_{ij} \ , \tag{11}$$

so we automatically obtain normalization. But the result we really care about is that

$$\boldsymbol{w} = \boldsymbol{D}^{-1/2}\tilde{\boldsymbol{v}} = \boldsymbol{D}^{-1/2}\boldsymbol{D}^{-1/2}\boldsymbol{v} = \boldsymbol{D}^{-1}\boldsymbol{v} \ , \tag{12}$$

which says that one can obtain the $k$-th component of a left eigenvector by dividing the $k$-th component of the corresponding right eigenvector by $\pi_k$. We will use this trick to avoid explicitly computing left eigenvectors.

##### 1.6 Projection

Let $\boldsymbol{H}$ be an $S \times S$ infinitesimal stochastic, diagonalizable Hamiltonian matrix. A strategy we will exploit in Sec. 5 and Sec. 6 is to connect it to an $(S-1)\times(S-1)$ matrix $\tilde{\boldsymbol{H}}$. The benefit of doing this—that is, of projecting the $S$-dimensional problem to an $(S-1)$-dimensional problem—is that it may be easier to solve for objects like eigenvectors.

Let $\tilde{\boldsymbol{H}}$ be the $(S-1)\times(S-1)$ matrix with components

$$\tilde{H}_{ij} = H_{ij} - H_{i,S-1} \tag{13}$$

for all $i, j \in \{0, ..., S-2\}$. (Note that we have reduced $\boldsymbol{H}$ by removing its last row and column. There is no loss of generality in doing this, since we can always relabel states to make some other index last.) Let $\boldsymbol{P}$ denote the $(S-1)\times S$ 'projection' matrix defined by

$$P_{ij} = \begin{cases} \delta_{ij} & \text{if } j < S-1 \\ 0 & \text{if } j = S-1 \end{cases} \tag{14}$$

and $\boldsymbol{Q}$ denote the $S \times (S-1)$ 'expansion' matrix defined by

$$Q_{ij} = \begin{cases} \delta_{ij} & \text{if } i < S-1 \\ -1 & \text{if } i = S-1 \end{cases} . \tag{15}$$

Note that $\tilde{\boldsymbol{H}} = \boldsymbol{PHQ}$, since

$$\sum_{b,c} P_{ab}H_{bc}Q_{cd} = \sum_{b<S-1,c<S-1} \delta_{ab}H_{bc}\delta_{cd} + \sum_{b<S-1,c=S-1} \delta_{ab}H_{bc}(-1) = H_{ad} - H_{a,S-1} = \tilde{H}_{ad} . \tag{16}$$

Also note that $\boldsymbol{PQ} = \boldsymbol{I}$ and that $\boldsymbol{QPx} = \boldsymbol{x}$ for all vectors satisfying $\mathbf{1}^T\boldsymbol{x} = 0$.

We mostly exploit the following facts.

1. If $\boldsymbol{v}$ is a right eigenvector of $\boldsymbol{H}$ whose corresponding eigenvalue is nonzero, then the vector $\tilde{\boldsymbol{v}} := \boldsymbol{Pv}$ (i.e., the vector formed by removing the last component of $\boldsymbol{v}$) is a right eigenvector of $\tilde{\boldsymbol{H}}$ with the same eigenvalue.

2. Similarly, if $\tilde{\boldsymbol{v}}$ is a right eigenvector of $\tilde{\boldsymbol{H}}$, then $\boldsymbol{v} := \boldsymbol{Q}\tilde{\boldsymbol{v}}$ is a right eigenvector of $\boldsymbol{H}$ with the same eigenvalue.

Let us demonstrate these claims.

**First claim.** Suppose $\boldsymbol{v}$ is an eigenvector of $\boldsymbol{H}$ whose corresponding eigenvalue is $\lambda \neq 0$. Then $\tilde{\boldsymbol{v}} := \boldsymbol{Pv}$ is an eigenvector of $\tilde{\boldsymbol{H}}$ with the same eigenvalue, because

$$\tilde{\boldsymbol{H}}\tilde{\boldsymbol{v}} = \boldsymbol{PHQPv} = \boldsymbol{PHv} = \lambda\boldsymbol{Pv} = \lambda\tilde{\boldsymbol{v}} , \tag{17}$$

where we used the fact that $\mathbf{1}^T\boldsymbol{v} = 0$ for all eigenvectors of $\boldsymbol{H}$ with nonzero eigenvalues.

**Second claim.** Suppose $\tilde{\boldsymbol{v}}$ is an eigenvector of $\tilde{\boldsymbol{H}}$. We claim that $\boldsymbol{v} := \boldsymbol{Q}\tilde{\boldsymbol{v}}$ is an eigenvector of $\boldsymbol{H}$ with the same eigenvalue. Note that the vector $\boldsymbol{H}\boldsymbol{v} = \boldsymbol{H}\boldsymbol{Q}\tilde{\boldsymbol{v}}$ sums to zero, since $\mathbf{1}^T\boldsymbol{H} = \mathbf{0}^T$. This means

$$\boldsymbol{H}\boldsymbol{v} = \boldsymbol{H}\boldsymbol{Q}\tilde{\boldsymbol{v}} = \boldsymbol{Q}\boldsymbol{P}\boldsymbol{H}\boldsymbol{Q}\tilde{\boldsymbol{v}} = \boldsymbol{Q}\tilde{\boldsymbol{H}}\tilde{\boldsymbol{v}} = \lambda\boldsymbol{Q}\tilde{\boldsymbol{v}} = \lambda\boldsymbol{v} \ . \tag{18}$$

There are two important consequences of these results. First, the eigenvalues of $\tilde{\boldsymbol{H}}$ are precisely the nonzero eigenvalues of $\boldsymbol{H}$. Second, if $\boldsymbol{\pi}$ denotes the (assumed unique) zero eigenvector of $\boldsymbol{H}$, then expressions of the form

$$\tilde{\boldsymbol{p}}(t) = \boldsymbol{P}\boldsymbol{\pi} - \sum_i c_i \tilde{\boldsymbol{v}}_i e^{\lambda_i t} \tag{19}$$

correspond to solutions

$$\boldsymbol{p}(t) = \boldsymbol{\pi} - \sum_i c_i \boldsymbol{v}_i e^{\lambda_i t} \tag{20}$$

of the master equation. Both facts allow us to study $\boldsymbol{H}$ and its associated master equation by analyzing $\tilde{\boldsymbol{H}}$.

#### 2 Mutual information and memory

We are primarily interested in answering three questions:

- How *long* can a memory switch store a memory? (*memory lifetime*)
- What is the best possible *fidelity* of a stored memory? (*memory capacity*)
- What measure determines whether one switch is better than another? (*memory performance*)

Answering the third question is what will allow us to assess which switch types and parameter settings are 'optimal', and in particular whether and how much Crick-like molecular error correction is useful in this setting.

Assume there is a binary stimulus $u$ that is either present ($u = 1$) or absent ($u = 0$). In the absence of any stimulus, we assume the system is at equilibrium, i.e., that $(d/dt)\boldsymbol{p}(t) = \mathbf{0}$, which implies that $\boldsymbol{p}(t|u = 0) = \boldsymbol{\pi}$. If the stimulus is present, we assume the system has a special initial condition $\boldsymbol{p}(t = 0|u = 1) = \boldsymbol{p}_*$ that is different from $\boldsymbol{\pi}$. Given this setup, measuring memory amounts to quantifying the time-dependent difference between $\boldsymbol{p}(t|u = 1)$ and $\boldsymbol{\pi}$. To ease notation, we will often abbreviate $\boldsymbol{p}(t|u = 1)$ as $\boldsymbol{p}(t)$ in what follows.

Our central idea is to quantify memory via mutual information (MI) [5, 6]. MI has been used to understand memory in neural systems, and is appealing partly because it offers a general way of quantifying memory that is applicable across many scales and substrates [7–9]. Here, we consider the MI between the binary stimulus $u$ and the system's state $s \in \{0, ..., S - 1\}$, which given the joint distribution $p(s, u|t) = p(s|u, t)p(u)$ and marginal $p(s|t) := \sum_{u'} p(s|u', t)p(u')$ is defined as

$$
\begin{aligned}
I(t) &:= \sum_{s,u} p(s|u, t)p(u) \log_2 \left( \frac{p(s|u, t)}{\sum_{u'} p(s|u', t)p(u')} \right) \\
&= 1 - \frac{1}{2} \sum_s \pi_s \log_2 \left( 1 + \frac{p_s(t)}{\pi_s} \right) - \frac{1}{2} \sum_s p_s(t) \log_2 \left( 1 + \frac{\pi_s}{p_s(t)} \right) \ ,
\end{aligned}
\tag{21}
$$

where we have assumed that our two stimulus conditions are equally likely ($p(u = 0) = p(u = 1) = 1/2$). We explicitly write $I(t)$ as time-dependent to emphasize that it is dynamic: by the data-processing inequality [6, 10], it is strictly *decreasing* in time. Moreover, the fact that $\lim_{t\to\infty} \boldsymbol{p}(t) = \boldsymbol{\pi}$ implies that $\lim_{t\to\infty} I(t) = 0$, i.e., that all information is eventually lost. Aside from monotonicity, $I(t)$ has other properties than make it a useful measure of memory. It is bounded between zero (no memory) and one (perfect memory) for all times $t$, is unitless, changes continuously with time, and is invariant to a relabeling of states.

We can use $I(t)$ to define sensible notions of lifetime, capacity, and performance. We will define **memory lifetime** as the amount of time it takes for $I(t)$ to decay to half of its original ($t = 0$) value.[1] This quantity is useful since it roughly captures the time scale on which information is lost, and is always well-defined, since $I(0) > 0$ and $\lim_{t\to\infty} I(t) = 0$.

[1] We obtain similar results by making slightly different choices, like defining lifetime as when $I(t)$ reaches $1/e$ of its original value.

We will define **memory capacity** as the maximum value of $I(t)$, i.e., as $C := I(0)$ since $I(t)$ decreases with time. We would like this number to be close to one, its maximum possible value, since otherwise the system is not even capable of *storing* information about a binary stimulus. In our setting, $C$ is essentially a transformed version of the false-positive probability $p_{\rm FP}$ (the steady state probability of being marked), since

$$\begin{aligned} I(0) &= 1 - \frac{1}{2} p_{\rm FP} \log_2 \left(1 + \frac{1}{p_{\rm FP}}\right) - \frac{1}{2} \log_2 (1 + p_{\rm FP}) \\ &= 1 + \frac{1}{2} p_{\rm FP} \log_2 p_{\rm FP} - \frac{1}{2} (1 + p_{\rm FP}) \log_2 (1 + p_{\rm FP}) \ . \end{aligned} \tag{22}$$

It is a monotonic function of $p_{\rm FP}$, and equals one when $p_{\rm FP} = 0$ and zero when $p_{\rm FP} = 1$.

Finally, we must define performance. At any fixed time $t$, it is always better for $I(t)$ to be higher—but how do we decide which overall *shape* of the $I(t)$ curve is better? Motivated by an analogy with (continuous-time) reinforcement learning [11, 12], if we view mutual information as a kind of reward, it is natural to define **memory performance** as the *discounted* area under the MI curve:

$$J := r \int_0^\infty e^{-rt} I(t) \ dt \leq 1 \tag{23}$$

where $1/r > 0$ is the discounting time scale. The discounting factor $e^{-rt}$ encodes a slight preference for remembering the stimulus better *earlier*; in other words, it is better to remember the stimulus with high fidelity, and then forget it, than to encode it extremely poorly for a longer duration. We say that one kind of memory switch is better than another if it yields a higher value of $J$. For example, given a fixed ambient marking rate $\epsilon > 0$ and discounting rate $r > 0$, there exists an optimal turnover rate $\gamma_*$ for the two-state model.

### 3 Pure turnover model

#### 3.1 Definition

The simplest model we consider is the **pure turnover model**, which has two states and only one possible transition: if the system is in its *marked* (+) state, it may randomly transition to its *unmarked* (−) state. Schematically:

$$- \xleftarrow{\gamma} + \tag{24}$$

where $\gamma > 0$ is the average rate of transition. This model abstracts the idea of a protein which randomly loses a molecular mark (e.g., a phosphate group) via unmarking, or because the marked protein degrades and is replaced by an unmarked protein.

This system is defined by the Hamiltonian

$$\boldsymbol{H} = \begin{pmatrix} 0 & \gamma \\ 0 & -\gamma \end{pmatrix} \ . \tag{25}$$

Let $\boldsymbol{p}(t) := (p_0(t), p_1(t))^T$ denote the time-dependent state occupancy probability distribution, $p_0(t)$ the probability of being in the unmarked state, and $p_1(t)$ the probability of being in the marked state. The master equation reads

$$\begin{aligned} \dot{p}_0(t) &= \gamma p_1(t) \\ \dot{p}_1(t) &= -\gamma p_1(t) \ . \end{aligned} \tag{26}$$

Since $p_0(t) + p_1(t) = 1$ for all $t$, this system's dynamics are completely characterized by the one-dimensional ODE

$$\dot{p}_1(t) = -\gamma p_1(t) \ . \tag{27}$$

#### 3.2 Eigenvalues

The characteristic polynomial associated with $\boldsymbol{H}$ is

$$\det(\boldsymbol{H} - \lambda \boldsymbol{I}) = \lambda(\gamma + \lambda) \tag{28}$$

which implies that the eigenvalues of $\boldsymbol{H}$ are

$$\boxed{\lambda = \lambda_0, \lambda_1 \ = 0, -\gamma} \ . \tag{29}$$

#### 3.3 Steady state probability distribution

The steady state probability distribution $\boldsymbol{\pi}$ (the eigenvector $\boldsymbol{v}_0$ with eigenvalue zero) satisfies

$$\boldsymbol{H} \begin{pmatrix} a \\ b \end{pmatrix} = \begin{pmatrix} 0 & \gamma \\ 0 & -\gamma \end{pmatrix} \begin{pmatrix} a \\ b \end{pmatrix} = \begin{pmatrix} 0 \\ 0 \end{pmatrix} \implies \gamma b = 0 \tag{30}$$

which implies that it is $(a, 0)^T$ up to normalization. Ensuring that its elements sum to one, we have

$$\boxed{\boldsymbol{\pi} = \boldsymbol{v}_0 = \begin{pmatrix} 1 \\ 0 \end{pmatrix}} \, . \tag{31}$$

This makes sense since the unmarked state is a 'sink'.

##### 3.4 Right eigenvectors

Let $\boldsymbol{v}_1$ denote the right eigenvector corresponding to $\lambda_1 = -\gamma$. Its elements must sum to zero (see Sec. 1.3), so $\boldsymbol{v}_1 = c(1, -1)^T$ for some normalization constant $c \neq 0$. We will choose $c = 1$ to adhere to our normalization convention (see Sec. 1.4), so

$$\boxed{\boldsymbol{v}_0 = \begin{pmatrix} 1 \\ 0 \end{pmatrix} \qquad \boldsymbol{v}_1 = \begin{pmatrix} 1 \\ -1 \end{pmatrix}} \, . \tag{32}$$

##### 3.5 Left eigenvectors

The vector $\boldsymbol{w}_0 = (1, 1)^T$ is always a left eigenvector with eigenvalue zero when $\boldsymbol{H}$ is infinitesimal stochastic (see Sec. 1.3). The other left eigenvector, $\boldsymbol{w}_1$, can easily be obtained by orthonormality. Since it is orthogonal to $\boldsymbol{v}_0$, it must equal $\boldsymbol{w}_1 = c(0, 1)^T$ for some nonzero $c$. To ensure $\boldsymbol{w}_1^T \boldsymbol{v}_1 = 1$, we can choose $c = -1$. Hence, the left eigenvectors are

$$\boxed{\boldsymbol{w}_0 = \begin{pmatrix} 1 \\ 1 \end{pmatrix} \qquad \boldsymbol{w}_1 = \begin{pmatrix} 0 \\ -1 \end{pmatrix}} \, . \tag{33}$$

It is easy to see that $\boldsymbol{w}_i^T \boldsymbol{v}_j = \delta_{ij}$.

##### 3.6 Transition probability

The general solution of this system's dynamics (Eq. 27) is

$$\begin{aligned} p_0(t) &= p_0(0) e^{-\gamma t} + (1 - e^{-\gamma t}) \\ p_1(t) &= p_1(0) e^{-\gamma t} \end{aligned} \tag{34}$$

where we have obtained $p_0(t)$ by solving the $p_1$ ODE and then using the fact that $p_0(t) = 1 - p_1(t)$. In the special case that $p_1(0) = 1$,

$$\boxed{\begin{aligned} p_0(t) &= 1 - e^{-\gamma t} \\ p_1(t) &= e^{-\gamma t} \end{aligned}} \, . \tag{35}$$

Hence, the dynamics are extremely simple: probability leaves the marked state exponentially quickly, at a rate $\gamma$. In terms of vectors, the transition probability can be written

$$\boxed{\boldsymbol{p}(t) = \boldsymbol{\pi} - c_1 \boldsymbol{v}_1 e^{\lambda_1 t} = \boldsymbol{\pi} - \boldsymbol{v}_1 e^{-\gamma t}} \, . \tag{36}$$

Note that $c_1 = 1$.

##### 3.7 False negative first-passage time distribution

How long does it take to transition from the marked state to the unmarked state? Equivalently, what is the first-passage time distribution associated with false negatives?

The survival probability is just $S(t) = p_1(t)$, since if the system leaves the marked state, it can never return to it. Hence, the first-passage time distribution is

$$\boxed{f(t) = -\frac{d}{dt}S(t) = \gamma e^{-\gamma t}} \ , \tag{37}$$

i.e., it is exponential, as expected. We immediately learn that the mean and variance are

$$\boxed{\mathbb{E}[t] = \frac{1}{\gamma} \qquad \mathrm{var}(t) = \frac{1}{\gamma^2}} \ . \tag{38}$$

### 4 Two-state model

#### 4.1 Definition

The two-state model abstracts the idea of a memory switch with marking noise. It has two states, *unmarked* ($-$) and *marked* ($+$), and two transitions: from marked to unmarked (turnover), and from unmarked to marked (marking noise). Schematically:

$$- \underset{\gamma}{\overset{\epsilon}{\rightleftarrows}} + \tag{39}$$

where $\gamma > 0$ is the rate of unmarking/turnover, and $\epsilon \geq 0$ is the basal marking rate. We will generally assume that $\gamma \geq \epsilon$, since otherwise it would not make sense to try and store memory by initializing the system in the marked state. Nonzero $\epsilon$ abstracts possibilities like ambient kinase activity. This type of memory switch is strictly worse than the pure turnover model, essentially because nonzero $\epsilon$ makes false positives possible.

Let $\boldsymbol{p}(t) := (p_0(t), p_1(t))^T$ denote the time-dependent state occupancy probability distribution, $p_0(t)$ the probability of being in the unmarked state, and $p_1(t)$ the probability of being in the marked state. This system's Hamiltonian is

$$\boldsymbol{H} = \begin{pmatrix} -\epsilon & \gamma \\ \epsilon & -\gamma \end{pmatrix} \tag{40}$$

and the corresponding master equation is

$$\begin{aligned} \dot{p}_0(t) &= -\epsilon p_0(t) + \gamma p_1(t) \\ \dot{p}_1(t) &= \epsilon p_0(t) - \gamma p_1(t) \ . \end{aligned} \tag{41}$$

Since $p_0(t) + p_1(t) = 1$, dynamics are completely characterized by the one-dimensional ODE

$$\dot{p}_1(t) = \epsilon - (\epsilon + \gamma) p_1(t) \ . \tag{42}$$

#### 4.2 Eigenvalues

The characteristic polynomial associated with $\boldsymbol{H}$ is

$$\det(\boldsymbol{H} - \lambda \boldsymbol{I}) = (\lambda + \epsilon)(\lambda + \gamma) - \gamma\epsilon = \lambda(\lambda + \epsilon + \gamma) \tag{43}$$

which implies that the eigenvalues of $\boldsymbol{H}$ are

$$\boxed{\lambda = \lambda_0, \lambda_1 \ = 0, -(\epsilon + \gamma)} \ . \tag{44}$$

#### 4.3 Steady state probability distribution

The steady state probability distribution $\boldsymbol{\pi}$ (the eigenvector $\boldsymbol{v}_0$ with eigenvalue zero) satisfies

$$\boldsymbol{H} \begin{pmatrix} a \\ b \end{pmatrix} = \begin{pmatrix} -\epsilon & \gamma \\ \epsilon & -\gamma \end{pmatrix} \begin{pmatrix} a \\ b \end{pmatrix} = \begin{pmatrix} 0 \\ 0 \end{pmatrix} \implies \gamma b = \epsilon a \tag{45}$$

which implies that it is $(\gamma, \epsilon)^T$ up to normalization. Ensuring that its elements sum to one,

$$\boxed{\boldsymbol{\pi} = \boldsymbol{v}_0 = \frac{1}{\epsilon + \gamma}\begin{pmatrix}\gamma \\ \epsilon\end{pmatrix}} \ . \tag{46}$$

Note that, even without a stimulus, there is a nonzero probability that the system exists in its marked state.

##### 4.4 Right eigenvectors

Let $\boldsymbol{v}_1$ denote the right eigenvector corresponding to $\lambda_1 = -(\epsilon + \gamma)$. As we argued in the case of the pure turnover model, since the elements of $\boldsymbol{v}_1$ must sum to zero (see Sec. 1.3), it equals $\boldsymbol{v}_1 = c(1, -1)^T$, where $c \neq 0$ is a normalization constant. We again choose $c = 1$ to adhere to our normalization convention (see Sec. 1.4), so

$$\boxed{\boldsymbol{v}_0 = \frac{1}{\epsilon + \gamma}\begin{pmatrix}\gamma \\ \epsilon\end{pmatrix} \qquad \boldsymbol{v}_1 = \begin{pmatrix}1 \\ -1\end{pmatrix}} \ . \tag{47}$$

##### 4.5 Left eigenvectors

The vector $\boldsymbol{w}_0 = (1, 1)^T$ is always a left eigenvector with eigenvalue zero when $\boldsymbol{H}$ is infinitesimal stochastic (see Sec. 1.3). To get the other left eigenvector, $\boldsymbol{w}_1$, we can use a trick. Note that this system satisfies detailed balance, since

$$H_{10}\pi_0 = \epsilon\gamma = H_{01}\pi_1 \ . \tag{48}$$

This implies (see Sec. 1.5) that, up to a normalization constant $c$,

$$\boldsymbol{w}_1 = c \ \mathrm{diag}(\boldsymbol{\pi})^{-1}\boldsymbol{v}_1 = c \ (\epsilon + \gamma)\begin{pmatrix}\frac{1}{\gamma} \\ -\frac{1}{\epsilon}\end{pmatrix} \ . \tag{49}$$

Since

$$\boldsymbol{w}_1^T\boldsymbol{v}_1 = c(\epsilon + \gamma)\left(\frac{1}{\gamma} + \frac{1}{\epsilon}\right) = c\frac{(\epsilon + \gamma)^2}{\epsilon\gamma} \ , \tag{50}$$

to ensure $\boldsymbol{w}_1^T\boldsymbol{v}_1 = 1$ we need $c = (\epsilon\gamma)/(\epsilon + \gamma)^2$, which means the left eigenvectors are

$$\boxed{\boldsymbol{w}_0 = \begin{pmatrix}1 \\ 1\end{pmatrix} \qquad \boldsymbol{w}_1 = \frac{1}{\epsilon + \gamma}\begin{pmatrix}\epsilon \\ -\gamma\end{pmatrix}} \ . \tag{51}$$

It is easy to see that $\boldsymbol{w}_i^T\boldsymbol{v}_j = \delta_{ij}$.

##### 4.6 Transition probability

We can directly solve Eq. 42 to find

$$\begin{aligned} p_0(t) &= p_0(0)e^{-(\epsilon+\gamma)t} + \frac{\gamma}{\epsilon+\gamma}\left[1 - e^{-(\epsilon+\gamma)t}\right] \\ p_1(t) &= p_1(0)e^{-(\epsilon+\gamma)t} + \frac{\epsilon}{\epsilon+\gamma}\left[1 - e^{-(\epsilon+\gamma)t}\right] \ . \end{aligned} \tag{52}$$

In the special case that $p_1(0) = 1$,

$$\boxed{\begin{aligned} p_0(t) &= \frac{\gamma}{\epsilon+\gamma}\left[1 - e^{-(\epsilon+\gamma)t}\right] \\ p_1(t) &= \frac{\epsilon}{\epsilon+\gamma} + \frac{\gamma}{\epsilon+\gamma}e^{-(\epsilon+\gamma)t} \end{aligned}} \ . \tag{53}$$

This result is somewhat more complicated than the analogous result for the pure turnover model, but the high-level behavior is fairly similar: the system approaches its steady state on a time scale $-1/\lambda_1 = 1/(\epsilon + \gamma)$.

In terms of vectors, this transition probability can be written

$$\boxed{\boldsymbol{p}(t) = \boldsymbol{\pi} - c_1 \boldsymbol{v}_1 e^{\lambda_1 t} = \boldsymbol{\pi} - \frac{\gamma}{\epsilon+\gamma}\boldsymbol{v}_1 e^{-(\epsilon+\gamma)t}} \ . \tag{54}$$

##### 4.7 False negative first-passage time distribution

To compute the first-passage time distribution, we can consider an 'amputated' system which has the unmarked to marked transition removed, i.e.,

$$- \ \overset{\gamma}{\leftarrow} \ + \ , \tag{55}$$

which is identical to the pure turnover model. This immediately implies that the survival probability is $S(t) = e^{-\gamma t}$, and hence that

$$\boxed{f(t) = -\frac{d}{dt}S(t) = \gamma e^{-\gamma t}} \tag{56}$$

as in the case of the pure turnover model. Also as before, we have moments

$$\boxed{\mathbb{E}[t] = \frac{1}{\gamma} \qquad \mathrm{var}(t) = \frac{1}{\gamma^2}} \ . \tag{57}$$

### 5 Pair model

#### 5.1 Definition

A natural extension of the two-state model, which represents a single molecular memory switch, is the model which describes a *pair* of independent and identical two-state switches. This abstracts the idea of two proteins which can be independently marked and unmarked, and which utilize the same marking and unmarking mechanisms. This model has three states—*unmarked* (−−), *partially marked* (+−), and *fully marked* (++)—and transitions between states which differ by one sign. Note that there are three states rather than four, since (because the two switches are indistinguishable) the state of the system does not depend on which switch is marked, only how many are marked.

Schematically, we have

$$-- \underset{\gamma}{\overset{2\epsilon}{\rightleftarrows}} +- \underset{2\gamma}{\overset{\epsilon}{\rightleftarrows}} ++ \tag{58}$$

where $\gamma > 0$ is the rate of unmarking/turnover and $\epsilon \geq 0$ is the ambient marking rate.

Let $\boldsymbol{p}(t) := (p_0(t), p_1(t), p_2(t))^T$ denote the time-dependent state occupancy distribution, $p_0(t)$ the probability of being in the unmarked state, $p_1(t)$ the probability of being in the partially marked state, and $p_2(t)$ the probability of being in the fully marked state. This system is defined by the Hamiltonian

$$\boldsymbol{H} = \begin{pmatrix} -2\epsilon & \gamma & 0 \\ 2\epsilon & -(\epsilon+\gamma) & 2\gamma \\ 0 & \epsilon & -2\gamma \end{pmatrix} \tag{59}$$

and the corresponding dynamics are

$$\begin{aligned} \dot{p}_0(t) &= -2\epsilon p_0(t) + \gamma p_1(t) \\ \dot{p}_1(t) &= 2\epsilon p_0(t) - (\epsilon+\gamma) p_1(t) + 2\gamma p_2(t) \\ \dot{p}_2(t) &= \epsilon p_1(t) - 2\gamma p_2(t) \ . \end{aligned} \tag{60}$$

Since $p_0(t) + p_1(t) + p_2(t) = 1$, we can rewrite this as

$$\begin{aligned} \dot{p}_0(t) &= \gamma - (2\epsilon+\gamma) p_0(t) - \gamma p_2(t) \\ \dot{p}_2(t) &= \epsilon - \epsilon p_0(t) - (2\gamma+\epsilon) p_2(t) \ . \end{aligned} \tag{61}$$

Importantly, the dynamics of the reduced probability vector $\tilde{\boldsymbol{p}}(t) := (p_0(t), p_2(t))^T$ can be written in terms of

$$\tilde{\boldsymbol{H}} := \begin{pmatrix} -(2\epsilon+\gamma) & -\gamma \\ -\epsilon & -(2\gamma+\epsilon) \end{pmatrix} \qquad \boldsymbol{b} := \begin{pmatrix} \gamma \\ \epsilon \end{pmatrix} \ , \tag{62}$$

and in particular

$$\frac{d}{dt}\tilde{\boldsymbol{p}}(t) = \tilde{\boldsymbol{H}}\tilde{\boldsymbol{p}}(t) + \boldsymbol{b} \ . \tag{63}$$

This fact is useful since (see Sec. 1.6) the nonzero eigenvalues of $\boldsymbol{H}$ are precisely the eigenvalues of $\tilde{\boldsymbol{H}}$, and the right eigenvectors of $\tilde{\boldsymbol{H}}$ and $\boldsymbol{H}$ can be derived from one another.

##### 5.2 Eigenvalues

The characteristic polynomial associated with $\tilde{\boldsymbol{H}}$ is

$$\begin{aligned}\det\left(\tilde{\boldsymbol{H}} - \lambda \boldsymbol{I}\right) &= (\lambda + 2\epsilon + \gamma)(\lambda + 2\gamma + \epsilon) - \gamma\epsilon \\ &= \lambda^2 + 3(\epsilon + \gamma)\lambda + 2(\epsilon + \gamma)^2 \\ &= (\lambda + \epsilon + \gamma)[\lambda + 2(\epsilon + \gamma)] \ .\end{aligned} \tag{64}$$

Since $\boldsymbol{H}$ is infinitesimal stochastic, one of its eigenvalues is zero. Since its nonzero eigenvalues are those of $\tilde{\boldsymbol{H}}$, the eigenvalues of $\boldsymbol{H}$ are

$$\boxed{\lambda = \lambda_0, \lambda_1, \lambda_2 \ = 0, -(\epsilon + \gamma), -2(\epsilon + \gamma)} \ . \tag{65}$$

Note that we have $\lambda_n = -n(\epsilon + \gamma)$ for $n \in \{0, 1, 2\}$, i.e., that the eigenvalues are integer multiples of $(\epsilon + \gamma)$. This will also be true in the $N$ switch case (see Sec. 7).

##### 5.3 Steady state probability distribution

The steady state probability distribution $\boldsymbol{\pi}$ (the eigenvector $\boldsymbol{v}_0$ with eigenvalue zero) satisfies

$$\boldsymbol{H}\begin{pmatrix} a \\ b \\ c \end{pmatrix} = \begin{pmatrix} -2\epsilon & \gamma & 0 \\ 2\epsilon & -(\epsilon + \gamma) & 2\gamma \\ 0 & \epsilon & -2\gamma \end{pmatrix}\begin{pmatrix} a \\ b \\ c \end{pmatrix} = \begin{pmatrix} 0 \\ 0 \\ 0 \end{pmatrix} \ , \tag{66}$$

which implies that

$$\begin{aligned}\gamma b &= 2\epsilon a \\ \epsilon b &= 2\gamma c \ .\end{aligned} \tag{67}$$

We can satisfy the first constraint by choosing $a \propto \gamma$ and $b \propto 2\epsilon$, and the second by choosing $b \propto 2\gamma$ and $c \propto \epsilon$. This means that both constraints are satisfied by $(\gamma^2, 2\epsilon\gamma, \epsilon^2)^T$. Normalizing this vector, we find

$$\boxed{\boldsymbol{\pi} = \boldsymbol{v}_0 = \frac{1}{(\epsilon + \gamma)^2}\begin{pmatrix} \gamma^2 \\ 2\gamma\epsilon \\ \epsilon^2 \end{pmatrix}} \ . \tag{68}$$

##### 5.4 Right eigenvectors

Let $\boldsymbol{v}_1$ denote the right eigenvector corresponding to $\lambda_1 = -(\epsilon + \gamma)$, and $\boldsymbol{v}_2$ the right eigenvector corresponding to $\lambda_2 = -2(\epsilon + \gamma)$. If $\tilde{\boldsymbol{v}}_1$ and $\tilde{\boldsymbol{v}}_2$ are the analogous right eigenvectors of $\tilde{\boldsymbol{H}}$, the condition $\tilde{\boldsymbol{H}} - \lambda \boldsymbol{I} = \boldsymbol{0}$ implies that their components $(a, b)^T$ satisfy

$$-(2\epsilon + \gamma + \lambda)a - \gamma b = 0 \ . \tag{69}$$

This implies that, up to normalization constants $c_i$,

$$\tilde{\boldsymbol{v}}_i = c_i \begin{pmatrix} \gamma \\ -(2\epsilon + \gamma + \lambda_i) \end{pmatrix} . \tag{70}$$

In particular,

$$\tilde{\boldsymbol{v}}_1 = c_1 \begin{pmatrix} \gamma \\ -\epsilon \end{pmatrix} \qquad \tilde{\boldsymbol{v}}_2 = c_2 \begin{pmatrix} \gamma \\ \gamma \end{pmatrix} . \tag{71}$$

We can convert these into right eigenvectors of $\boldsymbol{H}$ by choosing the last component so that they sum to zero (see Sec. 1.6), and then normalizing them so that their 1-norm equals two (see Sec. 1.4). Hence, the right eigenvectors of $\boldsymbol{H}$ are

$$\boxed{\boldsymbol{v}_0 = \frac{1}{(\epsilon+\gamma)^2} \begin{pmatrix} \gamma^2 \\ 2\gamma\epsilon \\ \epsilon^2 \end{pmatrix} \qquad \boldsymbol{v}_1 = \frac{1}{\gamma} \begin{pmatrix} \gamma \\ -\gamma+\epsilon \\ -\epsilon \end{pmatrix} \qquad \boldsymbol{v}_2 = \begin{pmatrix} \frac{1}{2} \\ -1 \\ \frac{1}{2} \end{pmatrix}} . \tag{72}$$

##### 5.5 Left eigenvectors

Let $\boldsymbol{w}_1$ denote the left eigenvector corresponding to $\lambda_1 = -(\epsilon+\gamma)$, and $\boldsymbol{w}_2$ the left eigenvector corresponding to $\lambda_2 = -2(\epsilon+\gamma)$. Like its single switch analogue, this system satisfies detailed balance since

$$\begin{aligned} H_{01}\pi_1 &= \frac{2\gamma^2\epsilon}{(\epsilon+\gamma)^2} = H_{10}\pi_0 \\ H_{12}\pi_2 &= \frac{2\gamma\epsilon^2}{(\epsilon+\gamma)^2} = H_{21}\pi_1 . \end{aligned} \tag{73}$$

This means that, up to normalization constants $c_i$,

$$\begin{aligned} \boldsymbol{w}_1 &= c_1 \operatorname{diag}(\boldsymbol{\pi})^{-1}\boldsymbol{v}_1 = c_1\frac{(\epsilon+\gamma)^2}{\gamma} \begin{pmatrix} \frac{1}{\gamma} \\ \frac{-\gamma+\epsilon}{2\gamma\epsilon} \\ -\frac{1}{\epsilon} \end{pmatrix} = \frac{c_1(\epsilon+\gamma)^2}{2\gamma^2\epsilon} \begin{pmatrix} 2\epsilon \\ -\gamma+\epsilon \\ -2\gamma \end{pmatrix} = d_1 \begin{pmatrix} 2\epsilon \\ -\gamma+\epsilon \\ -2\gamma \end{pmatrix} \\ \boldsymbol{w}_2 &= c_2 \operatorname{diag}(\boldsymbol{\pi})^{-1}\boldsymbol{v}_2 = c_2(\epsilon+\gamma)^2 \begin{pmatrix} \frac{1}{2\gamma^2} \\ -\frac{1}{2\gamma\epsilon} \\ \frac{1}{2\epsilon^2} \end{pmatrix} = \frac{c_2(\epsilon+\gamma)^2}{2\gamma^2\epsilon^2} \begin{pmatrix} \epsilon^2 \\ -\gamma\epsilon \\ \gamma^2 \end{pmatrix} = d_2 \begin{pmatrix} \epsilon^2 \\ -\gamma\epsilon \\ \gamma^2 \end{pmatrix} \end{aligned} \tag{74}$$

where we defined new normalization constants $d_i$ by combining all of the prefactors. We can choose the constants $d_i$ by insisting that $\boldsymbol{w}_1^T\boldsymbol{v}_1 = \boldsymbol{w}_2^T\boldsymbol{v}_2 = 1$. The first of these dot products is

$$\boldsymbol{w}_1^T\boldsymbol{v}_1 = \frac{d_1}{\gamma}\left[2\epsilon\gamma + (-\gamma+\epsilon)^2 + 2\epsilon\gamma\right] = \frac{d_1}{\gamma}(\epsilon+\gamma)^2 \tag{75}$$

which implies that $d_1 = \gamma/(\epsilon+\gamma)^2$. The second dot product is

$$\boldsymbol{w}_2^T\boldsymbol{v}_2 = \frac{d_2}{2}\left[\epsilon^2 + 2\gamma\epsilon + \gamma^2\right] = \frac{d_2}{2}(\epsilon+\gamma)^2 \tag{76}$$

which implies that $d_2 = 2/(\epsilon+\gamma)^2$. All of this together means that the left eigenvectors are

$$\boxed{\boldsymbol{w}_0 = \begin{pmatrix}1\\1\\1\end{pmatrix} \qquad \boldsymbol{w}_1 = \frac{\gamma}{(\epsilon+\gamma)^2}\begin{pmatrix}2\epsilon\\-\gamma+\epsilon\\-2\gamma\end{pmatrix} \qquad \boldsymbol{w}_2 = \frac{2}{(\epsilon+\gamma)^2}\begin{pmatrix}\epsilon^2\\-\gamma\epsilon\\\gamma^2\end{pmatrix}} \ . \tag{77}$$

Note that $\boldsymbol{w}_i^T\boldsymbol{v}_j = 0$ when $i \neq j$, and $\boldsymbol{w}_i^T\boldsymbol{v}_j = \delta_{ij}$ more generally.

##### 5.6 Transition probability

In principle, for any initial condition $\boldsymbol{p}(0)$ we have the generic expression

$$\boldsymbol{p}(t) = \boldsymbol{\pi} + \boldsymbol{w}_1^T\boldsymbol{p}(0)\ \boldsymbol{v}_1 e^{\lambda_1 t} + \boldsymbol{w}_2^T\boldsymbol{p}(0)\ \boldsymbol{v}_2 e^{\lambda_2 t} \ . \tag{78}$$

Suppose that $\boldsymbol{p}(0) = (0,0,1)^T$. Then

$$\boldsymbol{w}_1^T\boldsymbol{p}(0) = -\frac{2\gamma^2}{(\epsilon+\gamma)^2} \qquad \boldsymbol{w}_2^T\boldsymbol{p}(0) = \frac{2\gamma^2}{(\epsilon+\gamma)^2} \ , \tag{79}$$

which means that if $\boldsymbol{p}(0)$ is concentrated in the fully marked (++) state, the transition probability is

$$\begin{aligned}\boldsymbol{p}(t) &= \frac{1}{(\epsilon+\gamma)^2}\left\{\begin{pmatrix}\gamma^2\\2\gamma\epsilon\\\epsilon^2\end{pmatrix} - 2\gamma\begin{pmatrix}\gamma\\-\gamma+\epsilon\\-\epsilon\end{pmatrix}e^{-(\epsilon+\gamma)t} + \gamma^2\begin{pmatrix}1\\-2\\1\end{pmatrix}e^{-2(\epsilon+\gamma)t}\right\}\\ &= \frac{1}{(\epsilon+\gamma)^2}\begin{pmatrix}\gamma^2 - 2\gamma^2 e^{-(\epsilon+\gamma)t} + \gamma^2 e^{-2(\epsilon+\gamma)t}\\ 2\gamma\epsilon - 2\gamma(-\gamma+\epsilon)e^{-(\epsilon+\gamma)t} - 2\gamma^2 e^{-2(\epsilon+\gamma)t}\\ \epsilon^2 + 2\gamma\epsilon e^{-(\epsilon+\gamma)t} + \gamma^2 e^{-2(\epsilon+\gamma)t}\end{pmatrix}\end{aligned} \tag{80}$$

or equivalently

$$\boxed{\boldsymbol{p}(t) = \frac{1}{(\epsilon+\gamma)^2}\begin{pmatrix}\gamma^2[1-e^{-(\epsilon+\gamma)t}]^2\\ 2\gamma[\epsilon+\gamma e^{-(\epsilon+\gamma)t}][1-e^{-(\epsilon+\gamma)t}]\\ [\epsilon+\gamma e^{-(\epsilon+\gamma)t}]^2\end{pmatrix}} \ . \tag{81}$$

This transition probability can be rewritten in an illuminating way. Define

$$p(t) := \frac{\epsilon}{\epsilon+\gamma} + \frac{\gamma}{\epsilon+\gamma}e^{-(\epsilon+\gamma)t} \ . \tag{82}$$

This quantity is the probability that any *single* switch is marked at time $t$. (Note that it is identical to the two-state model's transition probability, see Sec. 4.) Our above result says that the probability $p(x|t)$ that $x \in \{0,1,2\}$ switches are marked at time $t$ equals

$$p(x|t) = \binom{x}{2} p(t)^x [1-p(t)]^{2-x} \ , \tag{83}$$

i.e., $p(x|t)$ is a binomial distribution with $N = 2$ and 'heads' probability $p(t)$. This makes sense since both switches are assumed to be independent. In fact, since each individual switch is Bernoulli-distributed with 'on' probability $p(t)$, we could have also derived this result by considering the distribution of

$$y := x_1 + x_2 \ , \tag{84}$$

where each $x_i$ is an independent Bernoulli random variable with 'on' probability $p(t)$. When we study $N$ independent switches (Sec. 7), we will see that this result generalizes in the obvious way.

For completeness' sake, the vector form of our result is

$$\boxed{\boldsymbol{p}(t) = \boldsymbol{\pi} - c_1 \boldsymbol{v}_1 e^{\lambda_1 t} - c_2 \boldsymbol{v}_2 e^{\lambda_2 t} = \boldsymbol{\pi} - \frac{2\gamma^2}{(\epsilon+\gamma)^2}\boldsymbol{v}_1 e^{-(\epsilon+\gamma)t} + \frac{2\gamma^2}{(\epsilon+\gamma)^2}\boldsymbol{v}_2 e^{-2(\epsilon+\gamma)t}} \ , \tag{85}$$

so

$$c_1 = \frac{2\gamma^2}{(\epsilon+\gamma)^2} \qquad\qquad c_2 = -\frac{2\gamma^2}{(\epsilon+\gamma)^2} \ . \tag{86}$$

##### 5.7 Transition probability, ODE method

There is an alternative approach to computing the transition probability which does not involve computing eigenvectors. The idea is to start with the reduced system of ODEs (Eq. 61) and differentiate. For the reader's convenience, we reproduce the reduced system here:

$$\begin{aligned} \dot{p}_0(t) &= \gamma - (2\epsilon+\gamma)p_0(t) - \gamma p_2(t) \\ \dot{p}_2(t) &= \epsilon - \epsilon p_0(t) - (2\gamma+\epsilon)p_2(t) \ . \end{aligned} \tag{87}$$

Taking another derivative of $p_2(t)$, we have

$$\begin{aligned} \ddot{p}_2(t) &= -\epsilon\dot{p}_0(t) - (2\gamma+\epsilon)\dot{p}_2(t) \\ &= -\epsilon\gamma + \epsilon(2\epsilon+\gamma)p_0(t) + \epsilon\gamma p_2(t) - (2\gamma+\epsilon)\dot{p}_2(t) \ . \end{aligned} \tag{88}$$

But Eq. 87 tells us that

$$\epsilon p_0(t) = \epsilon - (2\gamma+\epsilon)p_2(t) - \dot{p}_2(t) \ , \tag{89}$$

which implies

$$\ddot{p}_2(t) = -\epsilon\gamma + (2\epsilon+\gamma)\left[\epsilon - (2\gamma+\epsilon)p_2(t) - \dot{p}_2(t)\right] + \epsilon\gamma p_2(t) - (2\gamma+\epsilon)\dot{p}_2(t) \ . \tag{90}$$

This rearrangement is significant since our equation for $p_2(t)$ now does not involve $p_0(t)$. In other words, solving for the transition probability is essentially equivalent to solving a single second-order ODE. Rearranging this to obtain a more standard form,

$$\ddot{p}_2(t) + 3(\epsilon+\gamma)\dot{p}_2(t) + 2(\epsilon+\gamma)^2 p_2(t) = 2\epsilon^2 \ . \tag{91}$$

The characteristic equation associated with this linear ODE is

$$r^2 + 3(\epsilon + \gamma)r + 2(\epsilon + \gamma)^2 = 0 \tag{92}$$

and its solutions are

$$r_\pm = \frac{-3(\epsilon + \gamma) \pm \sqrt{(\epsilon + \gamma)^2}}{2} = -(\epsilon + \gamma), -2(\epsilon + \gamma) \ . \tag{93}$$

Note that these roots correspond exactly to the eigenvalues we found earlier.

We can find a particular solution of Eq. 91 by guessing; if we consider an ansatz $p_2(t) = c$,

$$2(\epsilon + \gamma)^2 c = 2\epsilon^2 \implies c = \frac{\epsilon^2}{(\epsilon + \gamma)^2} \ . \tag{94}$$

Hence, the general solution of Eq. 91 is

$$p_2(t) = \frac{\epsilon^2}{(\epsilon + \gamma)^2} + c_1 e^{-(\epsilon+\gamma)t} + c_2 e^{-2(\epsilon+\gamma)t} \ . \tag{95}$$

We can determine the constants $c_1$ and $c_2$ via initial conditions. We prescribe $p_0(0)$ and $p_2(0)$ for our original system (Eq. 87), and it turns out that this is equivalent to prescribing $p_2(0)$ and $\dot{p}_2(0)$ for Eq. 91. Note that Eq. 87 implies

$$\dot{p}_2(0) = \epsilon - \epsilon p_0(0) - (2\gamma + \epsilon) p_2(0) \ . \tag{96}$$

Imposing these conditions, we find the equations

$$\begin{aligned} p_2(0) &= \frac{\epsilon^2}{(\epsilon + \gamma)^2} + c_1 + c_2 \\ \epsilon - \epsilon p_0(0) - (2\gamma + \epsilon) p_2(0) &= -(\epsilon + \gamma)(c_1 + 2c_2) \ . \end{aligned} \tag{97}$$

We can ease notation somewhat by defining $p := \epsilon/(\epsilon + \gamma)$ and $q := \gamma/(\epsilon + \gamma)$. These equations then become

$$\begin{aligned} p_2(0) &= p^2 + c_1 + c_2 \\ -p[1 - p_0(0)] + (1 + q) p_2(0) &= c_1 + 2c_2 \ . \end{aligned} \tag{98}$$

The first equation implies $c_1 + c_2 = p_2(0) - p^2$, so the second implies

$$c_2 = -p[1 - p_0(0)] + q p_2(0) + p^2 \ . \tag{99}$$

This means

$$\begin{aligned} c_1 &= p[1 - p_0(0)] + p p_2(0) - 2p^2 \\ c_2 &= -p[1 - p_0(0)] + q p_2(0) + p^2 \ . \end{aligned} \tag{100}$$

In summary,

$$p_2(t) = \frac{\epsilon^2}{(\epsilon+\gamma)^2} + \left\{p[1-p_0(0)] + pp_2(0) - 2p^2\right\} e^{-(\epsilon+\gamma)t} + \left\{-p[1-p_0(0)] + qp_2(0) + p^2\right\} e^{-2(\epsilon+\gamma)t} \,. \tag{101}$$

We are still not quite done, since we need to compute $p_0(t)$ and $p_1(t)$. Since $p_0(t)$ can be written in terms of $p_2(t)$ and $\dot{p}_2(t)$,

$$\begin{aligned} pp_0(t) &= p - (1+q)p_2(t) - \frac{1}{\epsilon+\gamma}\dot{p}_2(t) \\ &= p - (1+q)\left[\pi_2 + c_1 e^{-(\epsilon+\gamma)t} + c_2 e^{-2(\epsilon+\gamma)t}\right] + \left[c_1 e^{-(\epsilon+\gamma)t} + 2c_2 e^{-2(\epsilon+\gamma)t}\right] \end{aligned} \tag{102}$$

which implies

$$p_0(t) = \frac{p-(1+q)\pi_2}{p} - \frac{q}{p} c_1 e^{-(\epsilon+\gamma)t} + c_2 e^{-2(\epsilon+\gamma)t} \,. \tag{103}$$

The additive constant is

$$1 - \frac{(2\gamma+\epsilon)\frac{\epsilon^2}{(\epsilon+\gamma)^2}}{\epsilon} = \frac{(\epsilon+\gamma)^2 - \epsilon(2\gamma+\epsilon)}{(\epsilon+\gamma)^2} = \frac{\gamma^2}{(\epsilon+\gamma)^2} \,. \tag{104}$$

The last thing we need is $p_1(t) = 1 - p_0(t) - p_2(t)$. More explicitly,

$$p_1(t) = \pi_1 + \frac{(q-p)}{p} c_1 e^{-(\epsilon+\gamma)t} + (1-2c_2) e^{-2(\epsilon+\gamma)t} \,. \tag{105}$$

Finally, we have

$$\boxed{\begin{aligned} p_0(t) &= \pi_0 - \frac{q}{p} c_1 e^{-(\epsilon+\gamma)t} + c_2 e^{-2(\epsilon+\gamma)t} \\ p_1(t) &= \pi_1 + \frac{(q-p)}{p} c_1 e^{-(\epsilon+\gamma)t} + (1-2c_2) e^{-2(\epsilon+\gamma)t} \\ p_2(t) &= \pi_2 + c_1 e^{-(\epsilon+\gamma)t} + c_2 e^{-2(\epsilon+\gamma)t} \end{aligned}} \tag{106}$$

where

$$\boxed{\begin{aligned} c_1 &= p[1-p_0(0)] + pp_2(0) - 2p^2 \\ c_2 &= -p[1-p_0(0)] + qp_2(0) + p^2 \end{aligned}} \,. \tag{107}$$

In the special case that $p_2(0) = 1$, we have the somewhat simpler coefficients

$$\boxed{\begin{aligned} c_1 &= 2pq = \frac{2\gamma\epsilon}{(\epsilon+\gamma)^2} \\ c_2 &= q - p + p^2 = q - pq = q^2 = \frac{\gamma^2}{(\epsilon+\gamma)^2} \end{aligned}} \tag{108}$$

which imply

$$\boxed{\begin{aligned} p_0(t) &= \frac{\gamma^2}{(\epsilon+\gamma)^2}\left[1-e^{-(\epsilon+\gamma)t}\right]^2 \\ p_1(t) &= \frac{2\gamma}{(\epsilon+\gamma)^2}\left[1-e^{-(\epsilon+\gamma)t}\right]\left[\epsilon+\gamma e^{-(\epsilon+\gamma)t}\right] \\ p_2(t) &= \frac{1}{(\epsilon+\gamma)^2}\left[\epsilon+\gamma e^{-(\epsilon+\gamma)t}\right]^2 \end{aligned}} \ . \tag{109}$$

##### 5.8 False negative first-passage time distribution

To compute the first-passage time distribution, consider the 'amputated' system

$$-- \overset{2\gamma}{\longleftarrow} +- \underset{2\gamma}{\overset{\epsilon}{\rightleftarrows}} ++ \tag{110}$$

where the unmarked state acts as a sink. The corresponding dynamics are

$$\begin{aligned} \dot{p}_1(t) &= -(\gamma+\epsilon)p_1(t) + 2\gamma p_2(t) \\ \dot{p}_2(t) &= \epsilon p_1(t) - 2\gamma p_2(t) \ . \end{aligned} \tag{111}$$

By definition, the first-passage time distribution $f(t)$ equals the negative time derivative of the survival probability (which equals $S(t) = p_1(t) + p_2(t)$), so

$$f(t) = -\dot{S}(t) = -[\dot{p}_1(t) + \dot{p}_2(t)] = \gamma p_1(t) \ . \tag{112}$$

Hence, to compute $f(t)$, it suffices to compute $p_1(t)$. We can compute $p_1(t)$ without computing $p_2(t)$ using the same trick we used in the previous subsection. Differentiate the $p_1(t)$ ODE with respect to time:

$$\begin{aligned} \ddot{p}_1(t) &= -(\gamma+\epsilon)\dot{p}_1(t) + 2\gamma\dot{p}_2(t) \\ &= -(\gamma+\epsilon)\dot{p}_1(t) + 2\gamma[\epsilon p_1(t) - 2\gamma p_2(t)] \ . \end{aligned} \tag{113}$$

Now note that

$$2\gamma p_2(t) = (\gamma+\epsilon)p_1(t) + \dot{p}_1(t) \ , \tag{114}$$

so

$$\begin{aligned} \ddot{p}_1(t) &= -(\gamma+\epsilon)\dot{p}_1(t) + 2\gamma\epsilon p_1(t) - 2\gamma[(\gamma+\epsilon)p_1(t) + \dot{p}_1(t)] \\ &= -(3\gamma+\epsilon)\dot{p}_1(t) - 2\gamma^2 p_1(t) \ . \end{aligned} \tag{115}$$

We conclude that $p_1(t)$ satisfies the linear, second-order ODE

$$\ddot{p}_1(t) + (3\gamma+\epsilon)\dot{p}_1(t) + 2\gamma^2 p_1(t) = 0 \ . \tag{116}$$

The characteristic equation of this ODE is

$$r^2 + (3\gamma+\epsilon)r + 2\gamma^2 = 0 \tag{117}$$

and its solutions are

$$r = \frac{-(3\gamma+\epsilon) \pm \sqrt{(\gamma+\epsilon)^2 + 4\gamma\epsilon}}{2} = -\gamma + \tilde{\Delta}, -2\gamma - \tilde{\Delta} - \epsilon \ , \tag{118}$$

where we define the spectral gap

$$\tilde{\Delta} := \frac{\sqrt{(\epsilon+\gamma)^2 + 4\gamma\epsilon} - (\epsilon+\gamma)}{2} \ . \tag{119}$$

Unlike the spectral gap that we later study in the context of the Crick switch, this gap equals zero when $\epsilon = 0$, and asymptotically becomes

$$\tilde{\Delta} = \frac{\epsilon\left[1 + 3\frac{\gamma}{\epsilon} - 4\frac{\gamma^2}{\epsilon^2}\right] - (\epsilon+\gamma)}{2} + \mathcal{O}(1/\epsilon^2) = \gamma - \frac{2\gamma^2}{\epsilon} + \mathcal{O}(1/\epsilon^2) \tag{120}$$

when $\epsilon$ is large. To ease notation, we will define

$$q_1 = \gamma - \tilde{\Delta} \qquad q_2 := 2\gamma + \tilde{\Delta} + \epsilon \ . \tag{121}$$

Note that (as one can see by inspecting the characteristic equation)

$$q_1 + q_2 = 3\gamma + \epsilon \qquad q_1 q_2 = 2\gamma^2 \ . \tag{122}$$

In any case, the general solution of our second-order ODE is

$$p_1(t) = c_1 e^{-q_1 t} + c_2 e^{-q_2 t} \tag{123}$$

where we must determine $c_1$ and $c_2$ through boundary conditions. To fully determine the original dynamics, we specify $p_1(0)$ and $p_2(0)$; it turns out this is equivalent to specifying $p_1(0)$ and $\dot{p}_1(0)$, since

$$\dot{p}_1(0) = -(\gamma+\epsilon)p_1(0) + 2\gamma p_2(0) \ . \tag{124}$$

Hence, we can determine $c_1$ and $c_2$ from the pair of conditions

$$\begin{aligned} p_1(0) &= c_1 + c_2 \\ -(\gamma+\epsilon)p_1(0) + 2\gamma p_2(0) &= -(q_1 c_1 + q_2 c_2) \ . \end{aligned} \tag{125}$$

Let us specialize to the case we are actually interested in, and choose $p_2(0) = 1$ and $p_1(0) = 0$. We immediately have that $c_1 = -c_2$, and

$$c_1 = \frac{2\gamma}{q_2 - q_1} \ . \tag{126}$$

This means that

$$p_1(t) = \frac{2\gamma}{q_2 - q_1}\left(e^{-q_1 t} - e^{-q_2 t}\right) \ , \tag{127}$$

and hence that the first-passage time distribution is

$$\boxed{f(t) = \frac{2\gamma^2}{q_2 - q_1}\left(e^{-q_1 t} - e^{-q_2 t}\right) = \frac{2\gamma^2}{2\tilde{\Delta} + \epsilon}\left(e^{-(\gamma - \tilde{\Delta})t} - e^{-(2\gamma + \tilde{\Delta} + \epsilon)t}\right)} \; . \tag{128}$$

It is easy to verify that this distribution is normalized, since

$$\int_0^\infty f(t)\; dt = \frac{2\gamma^2}{q_2 - q_1}\left[\frac{1}{q_1} - \frac{1}{q_2}\right] = \frac{2\gamma^2}{q_2 - q_1}\frac{(q_2 - q_1)}{q_1 q_2} = \frac{2\gamma^2}{q_1 q_2} = 1 \; . \tag{129}$$

We can now easily compute the mean and variance of this distribution. The mean is

$$\begin{aligned}
\mathbb{E}[t] &= \frac{2\gamma^2}{q_2 - q_1}\left[\frac{1}{q_1^2} - \frac{1}{q_2^2}\right] \\
&= \frac{2\gamma^2}{q_2 - q_1}\frac{(q_2^2 - q_1^2)}{(q_1 q_2)^2} \\
&= \frac{2\gamma^2}{q_2 - q_1}\frac{(q_2 - q_1)(q_2 + q_1)}{(q_1 q_2)^2} \\
&= \frac{q_2 + q_1}{q_1 q_2} \\
&= \frac{3\gamma + \epsilon}{2\gamma^2} \; .
\end{aligned} \tag{130}$$

The average of $t^2$ is

$$\begin{aligned}
\mathbb{E}[t^2] &= \frac{2\gamma^2}{q_2 - q_1}\left[\frac{2}{q_1^3} - \frac{2}{q_2^3}\right] \\
&= \frac{2q_1 q_2}{q_2 - q_1}\frac{(q_2^3 - q_1^3)}{(q_1 q_2)^3} \\
&= \frac{2q_1 q_2}{q_2 - q_1}\frac{(q_2 - q_1)(q_2^2 + q_1 q_2 + q_1^2)}{(q_1 q_2)^3} \\
&= \frac{2q_1 q_2}{q_2 - q_1}\frac{(q_2 - q_1)[(q_1 + q_2)^2 - q_1 q_2]}{(q_1 q_2)^3} \\
&= \frac{2[(q_1 + q_2)^2 - q_1 q_2]}{(q_1 q_2)^2} \; .
\end{aligned} \tag{131}$$

This implies that the variance of $t$ is

$$\mathrm{var}(t) = \mathbb{E}[t^2] - \mathbb{E}[t]^2 = \frac{[(q_1 + q_2)^2 - 2q_1 q_2]}{(q_1 q_2)^2} = \frac{(3\gamma + \epsilon)^2 - 4\gamma^2}{4\gamma^4} \; . \tag{132}$$

In summary, the low-order moments are

$$\boxed{\mathbb{E}[t] = \frac{3\gamma + \epsilon}{2\gamma^2} \qquad \mathrm{var}(t) = \frac{(3\gamma + \epsilon)^2 - 4\gamma^2}{4\gamma^4}} \; . \tag{133}$$

Note that the average first-passage time increases as $\gamma$ decreases, and in particular that it can be arbitrarily large. This behavior is quite different from the MI-defined lifetime, which goes like $1/\epsilon$ in the $\gamma \to 0$ limit.

Both quantities approach infinity as $\epsilon$ approaches infinity, although the coefficient of variation (i.e., the standard deviation divided by the mean) remains finite. More precisely,

$$\frac{\sqrt{\mathrm{var}(t)}}{\mathbb{E}[t]} = \frac{\sqrt{(3\gamma + \epsilon)^2 - 4\gamma^2}}{3\gamma + \epsilon} \ , \tag{134}$$

and

$$\lim_{\epsilon \to \infty} \frac{\sqrt{\mathrm{var}(t)}}{\mathbb{E}[t]} = 1 \ . \tag{135}$$

Another way of thinking about this fact follows from observing that, in the infinite $\epsilon$ limit, the distribution becomes effectively exponential, since one of the exponential terms dominates the other.

### 6 Crick switch

#### 6.1 Definition

What we call the *Crick switch* represents our attempt to concretely instantiate Crick's proposal for a memory switch with an error correction mechanism. Its definition is almost identical to that of our model of two independent switches (see Sec. 5), but has one crucial difference: we assume that some mechanism (e.g., an enzyme) increases the rate of the partially marked (+−) to fully marked (++) transition beyond its base value. Schematically, we have

$$-- \underset{\gamma}{\overset{2\epsilon}{\rightleftarrows}} +- \underset{2\gamma}{\overset{\epsilon+k}{\rightleftarrows}} ++ \tag{136}$$

where, as before, $\gamma > 0$ is the turnover rate and $\epsilon \geq 0$ is the ambient marking rate. Unlike before, there is a new rate $k \geq 0$ that plays an *error correction* role. In particular, if $k$ is somewhat large, when a stored memory is at risk of being 'lost' because the fully marked state transitions to the partially marked state, the system will tend to quickly return to the fully marked state. It is this tendency to quickly revert to the fully marked state that we are calling "error correction".

This system is defined by the Hamiltonian

$$\boldsymbol{H} = \begin{pmatrix} -2\epsilon & \gamma & 0 \\ 2\epsilon & -(\epsilon+\gamma+k) & 2\gamma \\ 0 & \epsilon+k & -2\gamma \end{pmatrix} \tag{137}$$

which is identical to the one from Sec. 5 when $k = 0$. The corresponding dynamics are

$$\begin{aligned} \dot{p}_0(t) &= -2\epsilon p_0(t) + \gamma p_1(t) \\ \dot{p}_1(t) &= 2\epsilon p_0(t) - (\epsilon+\gamma+k)p_1(t) + 2\gamma p_2(t) \\ \dot{p}_2(t) &= (\epsilon+k)p_1(t) - 2\gamma p_2(t)\ . \end{aligned} \tag{138}$$

Since $p_0(t) + p_1(t) + p_2(t) = 1$, we can rewrite this as

$$\begin{aligned} \dot{p}_0(t) &= \gamma - (2\epsilon+\gamma)p_0(t) - \gamma p_2(t) \\ \dot{p}_2(t) &= \epsilon + k - (\epsilon+k)p_0(t) - (2\gamma+\epsilon+k)p_2(t)\ . \end{aligned} \tag{139}$$

The dynamics of the reduced probability vector $\tilde{\boldsymbol{p}}(t) := (p_0(t), p_2(t))^T$ can be written in terms of

$$\tilde{\boldsymbol{H}} := \begin{pmatrix} -(2\epsilon+\gamma) & -\gamma \\ -(\epsilon+k) & -(2\gamma+\epsilon+k) \end{pmatrix} \qquad \boldsymbol{b} := \begin{pmatrix} \gamma \\ \epsilon+k \end{pmatrix}\ , \tag{140}$$

and in particular

$$\frac{d}{dt}\tilde{\boldsymbol{p}}(t) = \tilde{\boldsymbol{H}}\tilde{\boldsymbol{p}}(t) + \boldsymbol{b}\ . \tag{141}$$

As in Sec. 5, this fact is useful since (see Sec. 1.6) the nonzero eigenvalues of $\boldsymbol{H}$ are precisely the eigenvalues of $\tilde{\boldsymbol{H}}$, and the right eigenvectors of $\tilde{\boldsymbol{H}}$ and $\boldsymbol{H}$ can be derived from one another.

##### 6.2 Eigenvalues

The characteristic polynomial associated with $\tilde{\boldsymbol{H}}$ is

$$\begin{aligned}\det\left(\tilde{\boldsymbol{H}} - \lambda\boldsymbol{I}\right) &= (\lambda + 2\epsilon + \gamma)(\lambda + 2\gamma + \epsilon + k) - \gamma(\epsilon + k) \\ &= \lambda^2 + [3(\epsilon + \gamma) + k]\lambda + 2[(\epsilon + \gamma)^2 + k\epsilon] \ .\end{aligned} \tag{142}$$

We can use the quadratic formula to find that the roots of this polynomial are

$$\begin{aligned}\lambda &= \frac{-3(\epsilon + \gamma) - k \pm \sqrt{[3(\epsilon + \gamma) + k]^2 - 8[(\epsilon + \gamma)^2 + k\epsilon]}}{2} \\ &= \frac{-3(\epsilon + \gamma) - k \pm \sqrt{(\epsilon + \gamma + k)^2 + 4k(\gamma - \epsilon)}}{2} \ .\end{aligned} \tag{143}$$

We will find it convenient to define the *spectral gap*

$$\boxed{\Delta := \frac{\sqrt{(\epsilon + \gamma + k)^2 + 4k(\gamma - \epsilon)} - (\epsilon + \gamma + k)}{2} \geq 0} \tag{144}$$

since it will allow us to write the full set of eigenvalues of $\boldsymbol{H}$ as

$$\boxed{\lambda = \lambda_0, \lambda_1, \lambda_2 \quad = 0, -(\epsilon + \gamma) + \Delta, -2(\epsilon + \gamma) - \Delta - k} \ . \tag{145}$$

When $k = 0$, $\Delta = 0$, and these eigenvalues are precisely those of the pair of two-state models (see Sec. 5). We call $\Delta$ a 'gap' since, for $k > 0$, it pushes the nonzero eigenvalues apart: $\lambda_1$ becomes smaller in magnitude, and $\lambda_2$ becomes larger in magnitude. As discussed in the main text, this separation can help improve memory storage, since it helps create a long time scale.

As we will show in the next subsection, for large $k$, we approximately have

$$\lambda \approx 0, -2\epsilon, -(3\gamma + \epsilon + k) \tag{146}$$

since $\Delta = \gamma - \epsilon + \mathcal{O}(1/k)$ in this limit.

##### 6.3 Interesting properties of the spectral gap

Since the spectral gap $\Delta$ defined in the previous subsection essentially determines what makes the Crick switch interesting relative to its two-state relatives, it is worth briefly studying its properties. Below, we will assume that $\gamma > \epsilon$, since otherwise the architecture of the Crick switch does not make much sense.

**Nonnegativity.** First, $\Delta$ is nonnegative, since

$$\Delta = \frac{\sqrt{(\epsilon + \gamma + k)^2 + 4k(\gamma - \epsilon)} - (\epsilon + \gamma + k)}{2} \geq \frac{\sqrt{(\epsilon + \gamma + k)^2} - (\epsilon + \gamma + k)}{2} = 0 \ . \tag{147}$$

**Large $k$ value.** Next, note that the square root term equals

$$\begin{aligned}\sqrt{(\epsilon+\gamma+k)^2+4k(\gamma-\epsilon)} &= \sqrt{k^2+(6\gamma-2\epsilon)k+(\epsilon+\gamma)^2} \\ &= k\sqrt{1+(6\gamma-2\epsilon)\frac{1}{k}+\frac{(\epsilon+\gamma)^2}{k^2}} \\ &\approx k\left\{1+(3\gamma-\epsilon)\frac{1}{k}+\frac{1}{2}\frac{(\epsilon+\gamma)^2}{k^2}-\frac{1}{8}(6\gamma-2\epsilon)^2\frac{1}{k^2}\right\} \\ &= k+3\gamma-\epsilon-4\gamma(\gamma-\epsilon)\frac{1}{k}\end{aligned} \tag{148}$$

to first order in $1/k$. This means that, when $k$ is large, the gap equals

$$\boxed{\Delta=\gamma-\epsilon-2(\gamma-\epsilon)\frac{\gamma}{k}+\mathcal{O}(1/k^2)}\ . \tag{149}$$

Of particular interest is the infinite $k$ limit

$$\boxed{\lim_{k\to\infty}\Delta(k)=\gamma-\epsilon}\ , \tag{150}$$

since this controls how small the smallest eigenvalue of $\boldsymbol{H}$ can be. In particular, in this limit

$$\begin{aligned}\lambda_1 &\approx -(\epsilon+\gamma)+(\gamma-\epsilon)=-2\epsilon \\ \lambda_2 &\approx -[2(\epsilon+\gamma)+(\gamma-\epsilon)+k]=-(3\gamma+\epsilon+k)\ .\end{aligned} \tag{151}$$

**Polynomial equation.** By the definition of $\Delta$, it is trivial to note that it satisfies the polynomial equation

$$\Delta\left[\Delta+(\epsilon+\gamma+k)\right]=\frac{4k(\gamma-\epsilon)}{4}=k(\gamma-\epsilon)\ . \tag{152}$$

This property is useful for proving other properties of $\Delta$, and also for rationalizing denominators.

**Rationalizing denominators.** For example, we can use the polynomial equation to show that

$$\frac{2\gamma}{2\gamma+k+\Delta}=1-\frac{\Delta}{\gamma-\epsilon}\ . \tag{153}$$

The calculation is straightforward:

$$\begin{aligned}&\left[1-\frac{\Delta}{\gamma-\epsilon}\right][2\gamma+k+\Delta] \\ =\ &\left[1-\frac{\Delta}{\gamma-\epsilon}\right][\gamma-\epsilon+(\epsilon+\gamma+k+\Delta)] \\ =\ &\gamma-\epsilon-\Delta+\epsilon+\gamma+k+\Delta-\frac{1}{\gamma-\epsilon}\Delta(\epsilon+\gamma+k+\Delta) \\ =\ &\gamma-\epsilon-\Delta+\epsilon+\gamma+k+\Delta-k \\ =\ &2\gamma\ .\end{aligned} \tag{154}$$

**Monotonicity.** A final interesting property of $\Delta$ is that it is a monotonically increasing function of $k$. If we use $\Delta'$ to denote the derivative of $\Delta$ with respect to $k$, our polynomial equation (Eq. 152) implies

$$\Delta' \left[2\Delta + (\epsilon + \gamma + k)\right] = (\gamma - \epsilon) \tag{155}$$

or equivalently that

$$\Delta' = \frac{\gamma - \epsilon}{2\Delta + (\epsilon + \gamma + k)} > 0 \ , \tag{156}$$

where we have used the assumption that $\gamma > \epsilon$. Note that if $\gamma = \epsilon$, $\Delta = 0$, i.e., there is no eigenvalue gap. Since $\Delta$ is a monotonic function of $k$, its values at $k = 0$ and $k = \infty$ are lower and upper bounds:

$$0 \leq \Delta \leq \gamma - \epsilon \ . \tag{157}$$

##### 6.4 Steady state probability distribution

The steady state probability distribution $\boldsymbol{\pi}$ (the eigenvector $\boldsymbol{v}_0$ with eigenvalue zero) satisfies

$$\boldsymbol{H} \begin{pmatrix} a \\ b \\ c \end{pmatrix} = \begin{pmatrix} -2\epsilon & \gamma & 0 \\ 2\epsilon & -(\epsilon + \gamma + k) & 2\gamma \\ 0 & \epsilon + k & -2\gamma \end{pmatrix} \begin{pmatrix} a \\ b \\ c \end{pmatrix} = \begin{pmatrix} 0 \\ 0 \\ 0 \end{pmatrix} \ , \tag{158}$$

which implies that

$$\begin{aligned} \gamma b &= 2\epsilon a \\ (\epsilon + k) b &= 2\gamma c \ . \end{aligned} \tag{159}$$

We can satisfy the first constraint by choosing $a \propto \gamma$ and $b \propto 2\epsilon$, and the second by choosing $b \propto 2\gamma$ and $c \propto \epsilon + k$. This means that both constraints are satisfied by $(\gamma^2, 2\epsilon\gamma, \epsilon^2 + k\epsilon)^T$. Normalizing this vector, we find

$$\boxed{\boldsymbol{\pi} = \boldsymbol{v}_0 = \frac{1}{(\epsilon + \gamma)^2 + k\epsilon} \begin{pmatrix} \gamma^2 \\ 2\gamma\epsilon \\ \epsilon^2 + k\epsilon \end{pmatrix}} \ . \tag{160}$$

##### 6.5 Right eigenvectors

Let $\boldsymbol{v}_1$ denote the right eigenvector corresponding to $\lambda_1 = -(\epsilon + \gamma) + \Delta$, and $\boldsymbol{v}_2$ the right eigenvector corresponding to $\lambda_2 = -2(\epsilon + \gamma) - \Delta - k$. If $\tilde{\boldsymbol{v}}_1$ and $\tilde{\boldsymbol{v}}_2$ are the analogous right eigenvectors of $\tilde{\boldsymbol{H}}$, the condition

$$(\tilde{\boldsymbol{H}} - \lambda \boldsymbol{I}) \begin{pmatrix} a \\ b \end{pmatrix} = \begin{pmatrix} -(2\epsilon + \gamma + \lambda) & -\gamma \\ -(\epsilon + k) & -(2\gamma + \epsilon + k + \lambda) \end{pmatrix} \begin{pmatrix} a \\ b \end{pmatrix} = \begin{pmatrix} 0 \\ 0 \end{pmatrix} \tag{161}$$

implies that

$$-(2\epsilon + \gamma + \lambda) a - \gamma b = 0 \ . \tag{162}$$

This implies that, up to normalization constants $c_i$,

$$\tilde{\boldsymbol{v}}_i = c_i \begin{pmatrix} \gamma \\ -(2\epsilon + \gamma + \lambda_i) \end{pmatrix} \ . \tag{163}$$

In particular,

$$\tilde{\boldsymbol{v}}_1 = c_1 \begin{pmatrix} \gamma \\ -\epsilon - \Delta \end{pmatrix} \qquad \tilde{\boldsymbol{v}}_2 = c_2 \begin{pmatrix} \gamma \\ \gamma + \Delta + k \end{pmatrix} \ . \tag{164}$$

Since eigenvectors of $\tilde{\boldsymbol{H}}$ correspond to eigenvectors of $\boldsymbol{H}$ (see Sec. 1.6), up to normalization,

$$\boldsymbol{v}_1 = c_1 \begin{pmatrix} \gamma \\ -\gamma + \epsilon + \Delta \\ -\epsilon - \Delta \end{pmatrix} \qquad \boldsymbol{v}_2 = c_2 \begin{pmatrix} \gamma \\ -2\gamma - \Delta - k \\ \gamma + \Delta + k \end{pmatrix} \ . \tag{165}$$

To adhere to our normalization convention (see Sec. 1.4), we will choose

$$c_1 = \frac{1}{\gamma} \qquad c_2 = \frac{1}{2\gamma + \Delta + k} \tag{166}$$

since

$$\begin{aligned} \|\boldsymbol{v}_1\|_1 &= |c_1| \ [\gamma + (\gamma - \epsilon - \Delta) + (\epsilon + \Delta)] = |c_1| \ 2\gamma \\ \|\boldsymbol{v}_2\|_1 &= |c_2| \ [\gamma + (2\gamma + \Delta + k) + (\gamma + \Delta + k)] = |c_2| \ 2(2\gamma + \Delta + k) \ . \end{aligned} \tag{167}$$

Hence, the right eigenvectors of $\boldsymbol{H}$ are

$$\boxed{\boldsymbol{v}_0 = \frac{1}{(\epsilon+\gamma)^2 + k\epsilon} \begin{pmatrix} \gamma^2 \\ 2\gamma\epsilon \\ \epsilon^2 + k\epsilon \end{pmatrix} \quad \boldsymbol{v}_1 = \frac{1}{\gamma} \begin{pmatrix} \gamma \\ -\gamma + \epsilon + \Delta \\ -\epsilon - \Delta \end{pmatrix} \quad \boldsymbol{v}_2 = \frac{1}{2\gamma + \Delta + k} \begin{pmatrix} \gamma \\ -2\gamma - \Delta - k \\ \gamma + \Delta + k \end{pmatrix}} \ . \tag{168}$$

We can somewhat simplify $\boldsymbol{v}_2$ by rewriting it so that $\Delta$ does not appear in any denominator, and that no higher powers of $\Delta$ appear. (This can always be done, since $\Delta$ satisfies Eq. 152.) To do this, we can use an identity we derived earlier (Eq. 153) to write

$$\boldsymbol{v}_2 = \frac{1}{2\gamma} \left[1 - \frac{\Delta}{\gamma - \epsilon}\right] \begin{pmatrix} \gamma \\ -2\gamma - \Delta - k \\ \gamma + \Delta + k \end{pmatrix} \ . \tag{169}$$

A calculation similar to the one we used to derive Eq. 153 tell us that

$$\left[1 - \frac{\Delta}{\gamma - \epsilon}\right] (\gamma + k + \Delta) = \gamma \left[1 + \frac{\Delta}{\gamma - \epsilon}\right] \ . \tag{170}$$

We conclude that

$$\boldsymbol{v}_2 = \begin{pmatrix} \frac{1}{2}\left[1 - \frac{\Delta}{\gamma-\epsilon}\right] \\ -1 \\ \frac{1}{2}\left[1 + \frac{\Delta}{\gamma-\epsilon}\right] \end{pmatrix} \ . \tag{171}$$

Note that it is now obvious that these eigenvectors reduce to those of the pair model when $k = 0$, and that in the $k \to \infty$ limit they effectively reduce to those of the two-state model:

$$\lim_{k\to\infty} \boldsymbol{v}_1 = \frac{1}{\gamma}\begin{pmatrix}\gamma\\0\\-\gamma\end{pmatrix} = \begin{pmatrix}1\\0\\-1\end{pmatrix} \qquad \lim_{k\to\infty} \boldsymbol{v}_2 = \begin{pmatrix}0\\-1\\1\end{pmatrix} . \tag{172}$$

Technically, these eigenvectors correspond to something like a pair of two-state models, each operating at a different time scale. $\boldsymbol{v}_2$ operates on a fast time scale, and corresponds to error correction, while $\boldsymbol{v}_1$ operates on a slow time scale, and corresponds to information loss.

For ease of reference, we box the final form of the right eigenvectors:

$$\boxed{\boldsymbol{v}_0 = \frac{1}{(\epsilon+\gamma)^2 + k\epsilon}\begin{pmatrix}\gamma^2\\2\gamma\epsilon\\\epsilon^2 + k\epsilon\end{pmatrix} \quad \boldsymbol{v}_1 = \frac{1}{\gamma}\begin{pmatrix}\gamma\\-\gamma+\epsilon+\Delta\\-\epsilon-\Delta\end{pmatrix} \quad \boldsymbol{v}_2 = \begin{pmatrix}\frac{1}{2}\left[1 - \frac{\Delta}{\gamma-\epsilon}\right]\\-1\\\frac{1}{2}\left[1 + \frac{\Delta}{\gamma-\epsilon}\right]\end{pmatrix}} . \tag{173}$$

##### 6.6 Left eigenvectors

Let $\boldsymbol{w}_1$ denote the left eigenvector corresponding to $\lambda_1 = -(\epsilon+\gamma) + \Delta$, and $\boldsymbol{w}_2$ the left eigenvector corresponding to $\lambda_2 = -2(\epsilon+\gamma) - \Delta - k$. This system satisfies detailed balance, since

$$\begin{aligned} H_{01}\pi_1 &= \frac{2\gamma^2\epsilon}{(\epsilon+\gamma)^2 + k\epsilon} = H_{10}\pi_0 \\ H_{12}\pi_2 &= \frac{2\gamma\epsilon(\epsilon+k)}{(\epsilon+\gamma)^2 + k\epsilon} = H_{21}\pi_1 \ . \end{aligned} \tag{174}$$

This means that, up to normalization constants $c_i$,

$$\begin{aligned} \boldsymbol{w}_1 &= c_1 \mathrm{diag}(\boldsymbol{\pi})^{-1}\boldsymbol{v}_1 = c_1\frac{[(\epsilon+\gamma)^2 + k\epsilon]}{\gamma}\begin{pmatrix}\frac{1}{\gamma}\\\frac{-\gamma+\epsilon+\Delta}{2\gamma\epsilon}\\-\frac{(\epsilon+\Delta)}{\epsilon(\epsilon+k)}\end{pmatrix} = d_1\begin{pmatrix}2\epsilon(\epsilon+k)\\(-\gamma+\epsilon+\Delta)(\epsilon+k)\\-2\gamma(\epsilon+\Delta)\end{pmatrix} \\ \boldsymbol{w}_2 &= c_2 \mathrm{diag}(\boldsymbol{\pi})^{-1}\boldsymbol{v}_2 = c_2[(\epsilon+\gamma)^2 + k\epsilon]\begin{pmatrix}\frac{1}{2\gamma^2}\left[1 - \frac{\Delta}{\gamma-\epsilon}\right]\\-\frac{1}{2\gamma\epsilon}\\\frac{1}{2\epsilon(\epsilon+k)}\left[1 + \frac{\Delta}{\gamma-\epsilon}\right]\end{pmatrix} = d_2\begin{pmatrix}\epsilon(\epsilon+k)\left[1 - \frac{\Delta}{\gamma-\epsilon}\right]\\-\gamma(\epsilon+k)\\\gamma^2\left[1 + \frac{\Delta}{\gamma-\epsilon}\right]\end{pmatrix} \end{aligned}$$

where we defined new normalization constants $d_i$ by combining all of the prefactors. Note,

$$\boldsymbol{w}_1^T\boldsymbol{v}_1 = \frac{d_1}{\gamma}\left[2\gamma\epsilon(\epsilon+k) + (-\gamma+\epsilon+\Delta)^2(\epsilon+k) + 2\gamma(\epsilon+\Delta)^2\right] . \tag{175}$$

This expression does not appear to simplify. Hence, if we want $\boldsymbol{w}_1^T \boldsymbol{v}_1 = 1$, we need

$$d_1 = \frac{\gamma}{2\gamma\epsilon(\epsilon+k) + (-\gamma+\epsilon+\Delta)^2(\epsilon+k) + 2\gamma(\epsilon+\Delta)^2} \ . \tag{176}$$

Similarly,

$$\boldsymbol{w}_2^T \boldsymbol{v}_2 = \frac{d_2}{2}\left[\epsilon(\epsilon+k)\left(1 - \frac{\Delta}{\gamma-\epsilon}\right)^2 + 2\gamma(\epsilon+k) + \gamma^2\left(1 + \frac{\Delta}{\gamma-\epsilon}\right)^2\right] \ . \tag{177}$$

which means $\boldsymbol{w}_2^T \boldsymbol{v}_2 = 1$ forces

$$d_2 = \frac{2}{\epsilon(\epsilon+k)\left(1 - \frac{\Delta}{\gamma-\epsilon}\right)^2 + 2\gamma(\epsilon+k) + \gamma^2\left(1 + \frac{\Delta}{\gamma-\epsilon}\right)^2} \ . \tag{178}$$

We conclude that the left eigenvectors of $\boldsymbol{H}$ are

$$\boxed{\begin{aligned} \boldsymbol{w}_0 &= \begin{pmatrix} 1 \\ 1 \\ 1 \end{pmatrix} \\ \boldsymbol{w}_1 &= \frac{\gamma}{2\gamma\epsilon(\epsilon+k) + (-\gamma+\epsilon+\Delta)^2(\epsilon+k) + 2\gamma(\epsilon+\Delta)^2} \begin{pmatrix} 2\epsilon(\epsilon+k) \\ (-\gamma+\epsilon+\Delta)(\epsilon+k) \\ -2\gamma(\epsilon+\Delta) \end{pmatrix} \\ \boldsymbol{w}_2 &= \frac{2}{\epsilon(\epsilon+k)\left(1 - \frac{\Delta}{\gamma-\epsilon}\right)^2 + 2\gamma(\epsilon+k) + \gamma^2\left(1 + \frac{\Delta}{\gamma-\epsilon}\right)^2} \begin{pmatrix} \epsilon(\epsilon+k)\left[1 - \frac{\Delta}{\gamma-\epsilon}\right] \\ -\gamma(\epsilon+k) \\ \gamma^2\left[1 + \frac{\Delta}{\gamma-\epsilon}\right] \end{pmatrix} \end{aligned}} \ . \tag{179}$$

Because these normalization factors are cumbersome, we will not use them in the next two sections to compute the transition probability. Instead, we will exploit a certain trick.

##### 6.7 Eigenvectors of reduced Hamiltonian

Since the reduced Hamiltonian $\tilde{\boldsymbol{H}}$ is $2 \times 2$, and hence much easier to work with than $\boldsymbol{H}$, it is of interest to completely characterize its right and left eigenvectors. We can use these eigenvectors, instead of the left and right eigenvectors of $\boldsymbol{H}$, to more cleanly derive a formula for the transition probability.

We already derived the right eigenvectors of $\tilde{\boldsymbol{H}}$, since they correspond to those of $\boldsymbol{H}$:

$$\boxed{\tilde{\boldsymbol{v}}_1 = \frac{1}{\gamma}\begin{pmatrix} \gamma \\ -\epsilon-\Delta \end{pmatrix} \qquad \tilde{\boldsymbol{v}}_2 = \frac{1}{2\gamma+\Delta+k}\begin{pmatrix} \gamma \\ \gamma+\Delta+k \end{pmatrix}} \ . \tag{180}$$

The left eigenvectors of $\tilde{\boldsymbol{H}}$, $\tilde{\boldsymbol{w}}_1$ and $\tilde{\boldsymbol{w}}_2$, are less obviously related to those of $\boldsymbol{H}$, so we will derive them from scratch. Since the left eigenvectors satisfy

$$(\tilde{\boldsymbol{H}}^T - \lambda \boldsymbol{I})\begin{pmatrix} a \\ b \end{pmatrix} = \begin{pmatrix} -(2\epsilon+\gamma+\lambda) & -(\epsilon+k) \\ -\gamma & -(2\gamma+\epsilon+k+\lambda) \end{pmatrix}\begin{pmatrix} a \\ b \end{pmatrix} = \begin{pmatrix} 0 \\ 0 \end{pmatrix} \tag{181}$$

implies that

$$-(2\epsilon+\gamma+\lambda)a-(\epsilon+k)b=0\ . \tag{182}$$

This implies that, up to normalization constants $c_i$,

$$\tilde{\boldsymbol{w}}_i = c_i \begin{pmatrix} \epsilon+k \\ -(2\epsilon+\gamma+\lambda_i) \end{pmatrix}\ . \tag{183}$$

In particular,

$$\tilde{\boldsymbol{w}}_1 = c_1 \begin{pmatrix} \epsilon+k \\ -\epsilon-\Delta \end{pmatrix} \qquad \tilde{\boldsymbol{w}}_2 = c_2 \begin{pmatrix} \epsilon+k \\ \gamma+\Delta+k \end{pmatrix}\ . \tag{184}$$

To determine $c_1$ and $c_2$, the relevant dot products are

$$\begin{aligned} \tilde{\boldsymbol{w}}_1^T \tilde{\boldsymbol{v}}_1 &= \frac{c_1}{\gamma}\left[\gamma(\epsilon+k)+(\epsilon+\Delta)^2\right] \\ \tilde{\boldsymbol{w}}_2^T \tilde{\boldsymbol{v}}_2 &= \frac{c_2}{2\gamma+\Delta+k}\left[\gamma(\epsilon+k)+(\gamma+\Delta+k)^2\right]\ . \end{aligned} \tag{185}$$

If we want both of these to equal one, we find

$$\boxed{\tilde{\boldsymbol{w}}_1 = \frac{\gamma}{\gamma(\epsilon+k)+(\epsilon+\Delta)^2}\begin{pmatrix} \epsilon+k \\ -\epsilon-\Delta \end{pmatrix} \qquad \tilde{\boldsymbol{w}}_2 = \frac{2\gamma+\Delta+k}{\gamma(\epsilon+k)+(\gamma+\Delta+k)^2}\begin{pmatrix} \epsilon+k \\ \gamma+\Delta+k \end{pmatrix}}\ . \tag{186}$$

We will use the fact that $\tilde{\boldsymbol{w}}_i^T \tilde{\boldsymbol{v}}_j = \delta_{ij}$ in the next subsection. Note that we can use identities satisfied by $\Delta$ to slightly rewrite these left eigenvectors. It is true that

$$(\epsilon+\Delta)(\epsilon+\gamma+2\Delta+k) = \gamma(\epsilon+k)+(\epsilon+\Delta)^2 \tag{187}$$

and

$$(\gamma+\Delta+k)(\epsilon+\gamma+2\Delta+k) = \gamma(\epsilon+k)+(\gamma+\Delta+k)^2\ . \tag{188}$$

Also,

$$(\Delta+\epsilon)(\Delta+\gamma+k) = \gamma(\epsilon+k)\ . \tag{189}$$

We can establish (through straightforward algebra) these identities in the same way that we proved Eq. 153. They imply that

$$\boxed{\tilde{\boldsymbol{w}}_1 = \frac{\gamma}{(\epsilon+\Delta)(\epsilon+\gamma+2\Delta+k)}\begin{pmatrix} \epsilon+k \\ -\epsilon-\Delta \end{pmatrix} \quad \tilde{\boldsymbol{w}}_2 = \frac{2\gamma+\Delta+k}{(\gamma+\Delta+k)(\epsilon+\gamma+2\Delta+k)}\begin{pmatrix} \epsilon+k \\ \gamma+\Delta+k \end{pmatrix}}\ . \tag{190}$$

##### 6.8 Transition probability

The solution of the reduced dynamics (Eq. 141) is

$$\tilde{\boldsymbol{p}}(t) = \tilde{\boldsymbol{\pi}} - c_1\tilde{\boldsymbol{v}}_1 e^{\lambda_1 t} - c_2\tilde{\boldsymbol{v}}_2 e^{\lambda_2 t} = \tilde{\boldsymbol{\pi}} + [\tilde{\boldsymbol{w}}_1^T(\tilde{\boldsymbol{p}}(0)-\tilde{\boldsymbol{\pi}})]\tilde{\boldsymbol{v}}_1 e^{\lambda_1 t} + [\tilde{\boldsymbol{w}}_2^T(\tilde{\boldsymbol{p}}(0)-\tilde{\boldsymbol{\pi}})]\tilde{\boldsymbol{v}}_2 e^{\lambda_2 t} \tag{191}$$

where we define $\tilde{\boldsymbol{\pi}} := (\pi_0, \pi_2)^T$ as the projection of the steady state distribution of $\boldsymbol{H}$. We are especially interested in the case that $\tilde{\boldsymbol{p}}(0) = (0, 1)^T$, so we will compute the corresponding transition probability explicitly. Note,

$$\tilde{\boldsymbol{p}}(0) - \tilde{\boldsymbol{\pi}} = \frac{1}{(\epsilon+\gamma)^2 + k\epsilon} \begin{pmatrix} -\gamma^2 \\ \gamma^2 + 2\gamma\epsilon \end{pmatrix} \tag{192}$$

and, ignoring normalizations,

$$\begin{aligned} \boldsymbol{w}_1^T(\tilde{\boldsymbol{p}}(0) - \tilde{\boldsymbol{\pi}}) &\propto -(\epsilon+k)\gamma^2 - (\epsilon+\Delta)\gamma(\gamma+2\epsilon) = -\gamma\left[\gamma(\epsilon+k) + (\epsilon+\Delta)(\gamma+2\epsilon)\right] \\ \boldsymbol{w}_2^T(\tilde{\boldsymbol{p}}(0) - \tilde{\boldsymbol{\pi}}) &\propto -(\epsilon+k)\gamma^2 + (\gamma+k+\Delta)\gamma(\gamma+2\epsilon) = \gamma\left[-\gamma(\epsilon+k) + (\gamma+k+\Delta)(\gamma+2\epsilon)\right] \ . \end{aligned}$$

Overall, including normalizations,

$$\begin{aligned} \boldsymbol{w}_1^T(\tilde{\boldsymbol{p}}(0) - \tilde{\boldsymbol{\pi}}) &= -\frac{\gamma^2\left[\gamma(\epsilon+k) + (\epsilon+\Delta)(\gamma+2\epsilon)\right]}{(\epsilon+\Delta)(\epsilon+\gamma+2\Delta+k)[(\epsilon+\gamma)^2+k\epsilon]} \\ \boldsymbol{w}_2^T(\tilde{\boldsymbol{p}}(0) - \tilde{\boldsymbol{\pi}}) &= \frac{\gamma(2\gamma+\Delta+k)\left[-\gamma(\epsilon+k) + (\gamma+k+\Delta)(\gamma+2\epsilon)\right]}{(\gamma+\Delta+k)(\epsilon+\gamma+2\Delta+k)[(\epsilon+\gamma)^2+k\epsilon]} \ . \end{aligned}$$

Hence, the transition probability equals

$$\tilde{\boldsymbol{p}}(t) = \tilde{\boldsymbol{\pi}} - c_1\tilde{\boldsymbol{v}}_1 e^{\lambda_1 t} - c_2\tilde{\boldsymbol{v}}_2 e^{\lambda_2 t} \tag{193}$$

where

$$\begin{aligned} c_1 &= \frac{\gamma^2}{(\epsilon+\gamma+2\Delta+k)[(\epsilon+\gamma)^2+k\epsilon]} \cdot \frac{\left[\gamma(\epsilon+k) + (\epsilon+\Delta)(\gamma+2\epsilon)\right]}{\epsilon+\Delta} \\ c_2 &= -\frac{\gamma(2\gamma+\Delta+k)}{(\epsilon+\gamma+2\Delta+k)[(\epsilon+\gamma)^2+k\epsilon]} \cdot \frac{\left[-\gamma(\epsilon+k) + (\gamma+k+\Delta)(\gamma+2\epsilon)\right]}{\gamma+\Delta+k} \ . \end{aligned} \tag{194}$$

We can slightly simplify these coefficients by noting that

$$\frac{\left[\gamma(\epsilon+k) + (\epsilon+\Delta)(\gamma+2\epsilon)\right]}{\epsilon+\Delta} = \Delta+\gamma+k+\gamma+2\epsilon = \Delta + 2(\epsilon+\gamma) + k \tag{195}$$

and

$$\frac{\left[-\gamma(\epsilon+k) + (\gamma+k+\Delta)(\gamma+2\epsilon)\right]}{\gamma+\Delta+k} = -(\epsilon+\Delta)+\gamma+2\epsilon = \epsilon+\gamma-\Delta \ . \tag{196}$$

This means

$$\begin{aligned} c_1 &= \frac{\gamma^2}{(\epsilon+\gamma+2\Delta+k)[(\epsilon+\gamma)^2+k\epsilon]} \ \left[2(\epsilon+\gamma)+k+\Delta\right] \\ c_2 &= -\frac{\gamma(2\gamma+\Delta+k)}{(\epsilon+\gamma+2\Delta+k)[(\epsilon+\gamma)^2+k\epsilon]} \ \left[\epsilon+\gamma-\Delta\right] \ . \end{aligned} \tag{197}$$

Finally, since moving from the reduced system to the full system just amounts to the replacements $\tilde{\boldsymbol{\pi}} \to \boldsymbol{\pi}$, $\tilde{\boldsymbol{v}}_1 \to \boldsymbol{v}_1$, and $\tilde{\boldsymbol{v}}_2 \to \boldsymbol{v}_2$,

$$\boxed{\boldsymbol{p}(t) = \boldsymbol{\pi} - c_1 \boldsymbol{v}_1 e^{\lambda_1 t} - c_2 \boldsymbol{v}_2 e^{\lambda_2 t}} \tag{198}$$

with the above $c_1$ and $c_2$.

As a sanity check, consider the $k \to 0$ limit. We have

$$\begin{aligned} c_1 &= \frac{2\gamma^2}{(\epsilon+\gamma)^2} \\ c_2 &= -\frac{2\gamma^2}{(\epsilon+\gamma)^2} \ , \end{aligned} \tag{199}$$

which matches what we found for the pair model (see Sec. 5). On the other hand, in the $k \to \infty$ limit,

$$\begin{aligned} c_1 &\to \frac{\gamma^2}{(3\gamma - \epsilon + k)[(\epsilon+\gamma)^2 + k\epsilon]} \ (3\gamma + \epsilon + k) \approx \frac{\gamma^2}{(\epsilon+\gamma)^2 + k\epsilon} \\ c_2 &\to -\frac{\gamma(3\gamma - \epsilon + k)}{(3\gamma - \epsilon + k)[(\epsilon+\gamma)^2 + k\epsilon]} \ 2\epsilon = -\frac{2\epsilon\gamma}{(\epsilon+\gamma)^2 + k\epsilon} \ . \end{aligned} \tag{200}$$

##### 6.9 Transition probability, ODE method

It is also worth considering an alternate approach to computing the transition probability which does not involve computing eigenvectors. Here we largely repeat the argument from Sec. 5, but for the Crick switch. Our goal is to write down a second-order ODE for $p_2(t)$ that does not involve $p_0(t)$ and $p_1(t)$, which can then be solved in the usual way.

**Obtaining a second-order ODE in one variable.** We begin with the reduced dynamics of the Crick switch,

$$\begin{aligned} \dot{p}_0(t) &= \gamma - (2\epsilon + \gamma) p_0(t) - \gamma p_2(t) \\ \dot{p}_2(t) &= \epsilon + k - (\epsilon + k) p_0(t) - (2\gamma + \epsilon + k) p_2(t) \ . \end{aligned} \tag{201}$$

Taking another derivative of $p_2(t)$, we have

$$\begin{aligned} \ddot{p}_2(t) &= -(\epsilon + k)\dot{p}_0(t) - (2\gamma + \epsilon + k)\dot{p}_2(t) \\ &= -(\epsilon + k)\gamma + (\epsilon + k)(2\epsilon + \gamma) p_0(t) + (\epsilon + k)\gamma p_2(t) - (2\gamma + \epsilon + k)\dot{p}_2(t) \ . \end{aligned} \tag{202}$$

But, using Eq. 201, we can write $p_0(t)$ in terms of $p_2(t)$ and $\dot{p}_2(t)$:

$$(\epsilon + k) p_0(t) = (\epsilon + k) - (2\gamma + \epsilon + k) p_2(t) - \dot{p}_2(t) \ . \tag{203}$$

This implies

$$\ddot{p}_2(t) = -(\epsilon+k)\gamma + (2\epsilon+\gamma)\left[\epsilon + k - (2\gamma + \epsilon + k) p_2(t) - \dot{p}_2(t)\right] + (\epsilon+k)\gamma p_2(t) - (2\gamma+\epsilon+k)\dot{p}_2(t) \ . \tag{204}$$

This rearrangement is significant since our equation for $p_2(t)$ now does not involve $p_0(t)$. In other words, solving for the transition probability is essentially equivalent to solving a single second-order ODE. Rearranging this to obtain a more standard form,

$$\ddot{p}_2(t) + [3(\epsilon+\gamma) + k]\dot{p}_2(t) + 2[(\epsilon+\gamma)^2 + k\epsilon]p_2(t) = 2\epsilon(\epsilon+k) \ . \tag{205}$$

**Solving the second-order ODE.** The characteristic equation associated with this linear ODE is

$$r^2 + [3(\epsilon+\gamma) + k]r + 2[(\epsilon+\gamma)^2 + k\epsilon] = 0 \tag{206}$$

and its solutions are

$$r_\pm = \frac{-3(\epsilon+\gamma) - k \pm \sqrt{(\epsilon+\gamma+k)^2 + 4k(\gamma-\epsilon)}}{2} = -(\epsilon+\gamma)+\Delta, -2(\epsilon+\gamma)-\Delta-k \ . \tag{207}$$

These roots are, of course, the same as the eigenvalues we found before.

We can find a particular solution of Eq. 205 by considering an ansatz $p_2(t) = c$:

$$2[(\epsilon+\gamma)^2 + k\epsilon]c = 2\epsilon(\epsilon+k) \implies c = \frac{\epsilon(\epsilon+k)}{(\epsilon+\gamma)^2 + k\epsilon} \ . \tag{208}$$

Hence, the general solution of Eq. 205 is

$$p_2(t) = \frac{\epsilon(\epsilon+k)}{(\epsilon+\gamma)^2 + k\epsilon} + c_1 e^{-[(\epsilon+\gamma)-\Delta]t} + c_2 e^{-[2(\epsilon+\gamma)+\Delta+k]t} \ . \tag{209}$$

We can determine the constants $c_1$ and $c_2$ via initial conditions. We prescribe $p_0(0)$ and $p_2(0)$ for our original system (Eq. 201), and it turns out that this is equivalent to prescribing $p_2(0)$ and $\dot{p}_2(0)$ for Eq. 205. Note that Eq. 201 implies

$$\dot{p}_2(0) = \epsilon + k - (\epsilon+k)p_0(0) - (2\gamma+\epsilon+k)p_2(0) \ . \tag{210}$$

Imposing these conditions, we find the equations

$$\begin{aligned} p_2(0) &= \pi_2 + c_1 + c_2 \\ \epsilon + k - (\epsilon+k)p_0(0) - (2\gamma+\epsilon+k)p_2(0) &= -\left\{(\epsilon+\gamma-\Delta)c_1 + [2(\epsilon+\gamma)+\Delta+k]c_2\right\} \end{aligned} \tag{211}$$

Using the first equation, the second equation becomes

$$\begin{aligned} &\epsilon + k - (\epsilon+k)p_0(0) - (2\gamma+\epsilon+k)p_2(0) = -\left\{(\epsilon+\gamma-\Delta)[p_2(0)-\pi_2] + [\epsilon+\gamma+2\Delta+k]c_2\right\} \\ &\implies \epsilon + k - (\epsilon+k)p_0(0) - (\gamma+k+\Delta)p_2(0) - (\epsilon+\gamma-\Delta)\pi_2 = -(\epsilon+\gamma+2\Delta+k)c_2 \end{aligned}$$

which implies

$$c_2 = \frac{-(\epsilon+k)[1-p_0(0)] + (\gamma+k+\Delta)p_2(0) + (\epsilon+\gamma-\Delta)\pi_2}{\epsilon+\gamma+2\Delta+k} \ . \tag{212}$$

This means

$$c_1 = \frac{(\epsilon+k)[1-p_0(0)] + (\epsilon+\Delta)p_2(0) - [2(\epsilon+\gamma)+\Delta+k]\pi_2}{\epsilon+\gamma+2\Delta+k} \ . \tag{213}$$

**Solving for the other components of $\boldsymbol{p}(t)$.** Now we need to compute $p_0(t)$ and $p_1(t)$. Using the relationship between $p_0(t)$, $p_2(t)$, and $\dot{p}_2(t)$,

$$\begin{aligned}(\epsilon+k)p_0(t) &= (\epsilon+k) - (2\gamma+\epsilon+k)p_2(t) - \dot{p}_2(t) \\ &= (\epsilon+k) - (2\gamma+\epsilon+k)\left[\pi_2 + c_1 e^{-[(\epsilon+\gamma)-\Delta]t} + c_2 e^{-[2(\epsilon+\gamma)+\Delta+k]t}\right] + \left\{(\epsilon+\gamma-\Delta)c_1 e^{-[(\epsilon+\gamma)-\Delta]t}\right. \\ &\quad \left. + [2(\epsilon+\gamma)+\Delta+k]c_2 e^{-[2(\epsilon+\gamma)+\Delta+k]t}\right\} .\end{aligned} \tag{214}$$

This implies

$$\begin{aligned}p_0(t) &= \pi_0 + \frac{[\epsilon+\gamma-\Delta-(2\gamma+\epsilon+k)]}{\epsilon+k} c_1 e^{-[(\epsilon+\gamma)-\Delta]t} + \frac{[2(\epsilon+\gamma)+\Delta+k-(2\gamma+\epsilon+k)]}{\epsilon+k} c_2 e^{-[2(\epsilon+\gamma)+\Delta+k]t} \\ &= \pi_0 - \frac{(\gamma+\Delta+k)}{\epsilon+k} c_1 e^{-[(\epsilon+\gamma)-\Delta]t} + \frac{(\epsilon+\Delta)}{\epsilon+k} c_2 e^{-[2(\epsilon+\gamma)+\Delta+k]t} .\end{aligned} \tag{215}$$

The last thing we need is $p_1(t) = 1 - p_0(t) - p_2(t)$, which is

$$p_1(t) = \pi_1 + \left[\frac{(\gamma+\Delta+k)}{\epsilon+k} - 1\right] c_1 e^{-[(\epsilon+\gamma)-\Delta]t} - \left[\frac{(\epsilon+\Delta)}{\epsilon+k} + 1\right] c_2 e^{-[2(\epsilon+\gamma)+\Delta+k]t} . \tag{216}$$

Our final answer is

$$\boxed{\begin{aligned}p_0(t) &= \pi_0 - \frac{(\gamma+\Delta+k)}{\epsilon+k} c_1 e^{-[(\epsilon+\gamma)-\Delta]t} + \frac{(\epsilon+\Delta)}{\epsilon+k} c_2 e^{-[2(\epsilon+\gamma)+\Delta+k]t} \\ p_1(t) &= \pi_1 + \left[\frac{(\gamma+\Delta+k)}{\epsilon+k} - 1\right] c_1 e^{-[(\epsilon+\gamma)-\Delta]t} - \left[\frac{(\epsilon+\Delta)}{\epsilon+k} + 1\right] c_2 e^{-[2(\epsilon+\gamma)+\Delta+k]t} \\ p_2(t) &= \pi_2 + c_1 e^{-[(\epsilon+\gamma)-\Delta]t} + c_2 e^{-[2(\epsilon+\gamma)+\Delta+k]t}\end{aligned}} \tag{217}$$

where

$$\boxed{\begin{aligned}c_1 &= \frac{(\epsilon+k)[1-p_0(0)] + (\epsilon+\Delta)p_2(0) - [2(\epsilon+\gamma)+\Delta+k]\pi_2}{\epsilon+\gamma+2\Delta+k} \\ c_2 &= \frac{-(\epsilon+k)[1-p_0(0)] + (\gamma+k+\Delta)p_2(0) + (\epsilon+\gamma-\Delta)\pi_2}{\epsilon+\gamma+2\Delta+k}\end{aligned}} . \tag{218}$$

In the special case that $p_2(0) = 1$, the coefficients simplify to

$$\boxed{\begin{aligned}c_1 &= \frac{(2\epsilon+\Delta+k) - [2(\epsilon+\gamma)+\Delta+k]\pi_2}{\epsilon+\gamma+2\Delta+k} = \frac{\gamma[2\epsilon(\epsilon+\gamma) + (\gamma+2\epsilon)\Delta + k\gamma]}{[(\epsilon+\gamma)^2 + k\epsilon][\epsilon+\gamma+2\Delta+k]} \\ c_2 &= \frac{\gamma-\epsilon+\Delta+(\epsilon+\gamma-\Delta)\pi_2}{\epsilon+\gamma+2\Delta+k} = \frac{\gamma[\gamma(\epsilon+\gamma) + (\gamma+2\epsilon)\Delta + 2k\epsilon]}{[(\epsilon+\gamma)^2 + k\epsilon][\epsilon+\gamma+2\Delta+k]}\end{aligned}} . \tag{219}$$

The expressions are now more complicated, so it seems unhelpful to write them out in full. But to illustrate this point, one has

$$\boxed{p_2(t) = \pi_2 + \frac{\gamma\left\{[2\epsilon(\epsilon+\gamma) + k\gamma + (\gamma+2\epsilon)\Delta]e^{-[\epsilon+\gamma-\Delta]t} + [\gamma(\epsilon+\gamma) + (\gamma+2\epsilon)\Delta + 2k\epsilon]e^{-[2(\epsilon+\gamma)+\Delta+k]t}\right\}}{[\epsilon+\gamma+2\Delta+k][(\epsilon+\gamma)^2 + k\epsilon]}} . \tag{220}$$

One can show, through tedious algebra involving spectral gap identities, that the result obtained using this method is equivalent to the one we obtained using the eigenvector method. The apparent difference here is due to how the eigenvectors that implicitly appear are normalized. For example, a scaled version of $\boldsymbol{v}_1$ appears if one inspects the coefficients of $e^{\lambda_1 t}$:

$$\begin{pmatrix} -\frac{(\gamma+\Delta+k)}{\epsilon+k} \\ \left[\frac{(\gamma+\Delta+k)}{\epsilon+k} - 1\right] \\ 1 \end{pmatrix} . \tag{221}$$

Its normalization is (implicitly) chosen so that the last component equals one.

##### 6.10 False negative first-passage time distribution

To compute the first-passage time distribution, we must consider the 'amputated' system

$$-- \overset{2\gamma}{\longleftarrow} +- \underset{2\gamma}{\overset{\epsilon+k}{\rightleftarrows}} ++ \tag{222}$$

where the unmarked state acts as a sink. Fortunately for us, this is exactly the same system we considered when computing the first-passage time distribution in Sec. 5 (for the pair of two-state models), with the only difference being that the partially marked to fully marked transition was labeled with a rate $\epsilon$ instead of $\epsilon + k$. Hence, we can reuse the result from that section, which is that the first-passage time distribution $f(t)$ is

$$\boxed{f(t) = \frac{2\gamma^2}{q_2 - q_1}\left(e^{-q_1 t} - e^{-q_2 t}\right) = \frac{2\gamma^2}{2\tilde{\Delta} + \epsilon + k}\left(e^{-(\gamma-\tilde{\Delta})t} - e^{-(2\gamma+\tilde{\Delta}+\epsilon+k)t}\right)} , \tag{223}$$

where we must define the rates

$$q_1 = \gamma - \tilde{\Delta} \qquad q_2 := 2\gamma + \tilde{\Delta} + \epsilon + k \tag{224}$$

and spectral gap

$$\tilde{\Delta} := \frac{\sqrt{(\epsilon + k + \gamma)^2 + 4\gamma(\epsilon + k)} - (\epsilon + k + \gamma)}{2} . \tag{225}$$

Moreover, the mean and variance of this distribution are

$$\boxed{\mathbb{E}[t] = \frac{3\gamma + \epsilon + k}{2\gamma^2} \qquad \text{var}(t) = \frac{(3\gamma + \epsilon + k)^2 - 4\gamma^2}{4\gamma^4}} . \tag{226}$$

In the large $k$ limit, note that

$$\mathbb{E}[t] \approx \frac{k}{2\gamma^2} \qquad \text{var}(t) \approx \frac{k^2}{4\gamma^4} , \tag{227}$$

which reflects the fact that $f(t)$ becomes approximately exponential in this limit. This happens since $f(t)$ becomes dominated by the slow time scale, i.e.,

$$f(t) \approx \frac{2\gamma^2}{k} e^{-2\frac{\gamma^2}{k}t} \ , \tag{228}$$

where we used the asymptotic result

$$\tilde{\Delta} = \gamma - \frac{2\gamma^2}{\epsilon + k} + \mathcal{O}(1/(\epsilon + k)^2) \ . \tag{229}$$

### 7 Many independent and indistinguishable two-state models

#### 7.1 Definition

In Sec. 5, we generalized the two-state model by considering a pair of independent and identical two-state models. There is an obvious generalization of the pair model, and more broadly of any molecular memory switch: the model which consists of $N \geq 1$ independent and indistinguishable copies of that switch. We expect that $N$ copies of a switch are strictly better at storing information than a single copy, and in the main text show that this is true in the case of the two-state model and Crick switch. In this section and the next, our goal is to show that the solutions of the single switch models can be used to straightforwardly solve their $N$-switch generalizations.

The two-state model has (as its name suggests) 2 states, unmarked and marked. Its $N$-switch generalization has $N+1$ states: 0 switches are marked, 1 switch is marked, and so on, all the way up to $N$ switches being marked. To describe this system in more technical detail, it is useful to invoke the mathematical formalism associated with the chemical master equation (CME). The CME description of this system follows from assuming two possible chemical reactions:

$$- \xrightarrow{\epsilon} + \qquad\qquad + \xrightarrow{\gamma} - \tag{230}$$

and elementary propensity functions (i.e., the forward rate is proportional to the number of unmarked switches, and the backward rate is proportional to the number of marked switches). The CME that describes the time evolution of the probability $p(x|t)$ that $x \in \{0, 1, ..., N\}$ switches are marked at time $t \geq 0$ is

$$\frac{\partial p(x|t)}{\partial t} = \epsilon \left[ (N-x+1)p(x-1|t) - (N-x)p(x|t) \right] + \gamma \left[ (x+1)p(x+1|t) - xp(x|t) \right] \ . \tag{231}$$

As a boundary condition, we define $p(x|t) = 0$ for $x > N$ and $x < 0$. We are specifically interested in the initial condition $p(x|0) = \delta_{x,N}$, i.e., with all switches initially marked.

As before, we can also write this CME as

$$\frac{\partial \boldsymbol{p}(t)}{\partial t} = \boldsymbol{H}\boldsymbol{p}(t) \tag{232}$$

where $\boldsymbol{p}(t) := (p_0(t), ..., p_N(t))^T$ and the Hamiltonian $\boldsymbol{H}$ has size $(N+1) \times (N+1)$.

#### 7.2 Transition probability

A direct way to solve the system consisting of $N$ two-state models is to find the eigenvalues and eigenvectors associated with the Hamiltonian $\boldsymbol{H}$. But this is both generally difficult and unnecessary. We can instead exploit the fact that we have already solved the (single protein) two-state model, and that this system consists of many two-state models.

The idea is as follows. Let $x_i$ be a random variable with two possible states, which we label as 0 and 1. Assume it takes a value of 1 with a probability $p_1(t)$ that corresponds to the marked probability of the two-state model:

$$p_1(t) = \frac{\epsilon}{\epsilon+\gamma} + \frac{\gamma}{\epsilon+\gamma} e^{-(\epsilon+\gamma)t} \ . \tag{233}$$

Let $z_1, ..., z_N$ be independent binary random variables like this, and consider $x := z_1 + \cdots z_N$. By definition, $x$ takes values between 0 and $N$, and moreover tracks the overall number of marked switches. Since the distribution of a sum of $N$ independent and identically distributed Bernoulli random variables is binomial, we have

$$\boxed{p(x|t) = \binom{N}{x} p_1(t)^x p_0(t)^{N-x}} \ . \tag{234}$$

This distribution satisfies the desired initial condition, since $p(x|0)$ is zero unless $x = N$. Moreover, it is easy to show that this distribution satisfies Eq. 231 for all times $t$. Note that

$$\begin{aligned} p(x-1|t) &= \frac{x}{N-x+1} \frac{p_0(t)}{p_1(t)} \, p(x|t) \\ p(x+1|t) &= \frac{(N-x)}{x+1} \frac{p_1(t)}{p_0(t)} \, p(x|t) \end{aligned} \tag{235}$$

and that

$$\begin{aligned} \frac{\partial p(x|t)}{\partial t} &= \binom{N}{x} \left\{ x \frac{\dot{p}_1(t)}{p_1(t)} + (N-x) \frac{\dot{p}_0(t)}{p_0(t)} \right\} p_1(t)^x p_0(t)^{N-x} \\ &= \left\{ x \left( \epsilon \frac{p_0(t)}{p_1(t)} - \gamma \right) + (N-x) \left( \gamma \frac{p_1(t)}{p_0(t)} - \epsilon \right) \right\} p(x|t) \ . \end{aligned} \tag{236}$$

Putting these facts together, we have

$$\frac{\partial p(x|t)}{\partial t} = \{ \epsilon(N-x+1) p(x-1|t) - \gamma x p(x|t) + \gamma(x+1) p(x+1|t) - \epsilon(N-x) p(x|t) \} \ . \tag{237}$$

But this is just Eq. 231, so we are done. It is worth mentioning that this strategy works regardless of the initial condition we use, since it only depends on the single protein master equation. Here, we assumed the system begins in its marked state with probability one, but we could make a different assumption and still exactly solve the corresponding multi-switch model.

##### 7.3 Moments

We can easily compute the mean and variance of $p(x|t)$, since it is a binomial distribution:

$$\begin{aligned} \mathbb{E}[x] &= p_1(t) \\ \text{var}(x) &= N p_1(t) [1 - p_1(t)] \ . \end{aligned} \tag{238}$$

Note that the moments of the scaled variable $y := x/N$, which measures the *fraction* rather than the *number* of marked switches, follow straightforwardly from these:

$$\begin{aligned} \mathbb{E}[y] &= p_1(t) \\ \mathrm{var}(y) &= \frac{1}{N} p_1(t)[1 - p_1(t)] \ . \end{aligned} \tag{239}$$

#### 7.4 Eigenvalues

Usefully, the transition probability implies the eigenvalues of the Hamiltonian, because

$$p(x|t) = \binom{N}{x} p_1(t)^x p_0(t)^{N-x} = \binom{N}{x} \left[ \frac{\epsilon}{\epsilon+\gamma} + \frac{\gamma}{\epsilon+\gamma} e^{-(\epsilon+\gamma)t} \right]^x \left[ \frac{\gamma}{\epsilon+\gamma} \left( 1 - e^{-(\epsilon+\gamma)t} \right) \right]^{N-x} \tag{240}$$

can be written in the form

$$p(x|t) = \pi(x) - \sum_i c_i v_i(x) e^{\lambda_i t} \ , \tag{241}$$

i.e., as a sum of terms each associated with a different exponential. Because there exists only one such expansion of $p(x|t)$, the arguments of those exponentials must correspond to the eigenvalues of $\boldsymbol{H}$. (We could be more careful, and establish this claim more rigorously, but we will not pursue that here.)

From the form of the transition probability, it is clear that only powers of $e^{-(\epsilon+\gamma)t}$ will appear. It must be the case, then, that the eigenvalues of $\boldsymbol{H}$ are

$$\boxed{\lambda = 0, -(\epsilon+\gamma), -2(\epsilon+\gamma), ..., -N(\epsilon+\gamma)} \ . \tag{242}$$

This is consistent with what we know of the two-state model (Sec. 4) and pair model (Sec. 5), which correspond to the $N = 1$ and $N = 2$ cases.

### 8 Many independent and indistinguishable Crick switches

#### 8.1 Definition

We can also consider the $N$-switch generalization of the Crick switch, and use a strategy similar to the one we used in the previous section to exactly solve it.

The Crick switch has 3 states (unmarked, partially marked, and fully marked), so its $N$-switch generalization has $\binom{N+2}{2} = (N+1)(N+2)/2$ states. This follows from a basic combinatorics argument. One can divide $N$ identical objects into 3 groups by lining them up and placing 2 walls somewhere in the sequence of objects. From a 'stars and bars' perspective, we can imagine $N+2$ objects lined up, and choose 2 of them to be walls.

The CME description of this system follows from assuming four possible reactions:

$$-- \xrightarrow{2\epsilon} +- \qquad +- \xrightarrow{\gamma} -- \qquad +- \xrightarrow{\epsilon+k} ++ \qquad ++ \xrightarrow{2\gamma} +- \tag{243}$$

and elementary propensity functions. Let $p(x_0, x_1, x_2|t)$ denote the probability that $x_0$ switches are unmarked, $x_1$ are partially marked, and $x_2$ fully marked. The CME that describes how it evolves in time is

$$\begin{aligned}\frac{\partial p(\boldsymbol{x}|t)}{\partial t} &= 2\epsilon\left[(x_0+1)p(x_0+1, x_1-1, x_2|t) - x_0 p(\boldsymbol{x}|t)\right] \\ &+ \gamma\left[(x_1+1)p(x_0-1, x_1+1, x_2|t) - x_1 p(\boldsymbol{x}|t)\right] \\ &+ (\epsilon+k)\left[(x_1+1)p(x_0, x_1+1, x_2-1|t) - x_1 p(\boldsymbol{x}|t)\right] \\ &+ 2\gamma\left[(x_2+1)p(x_0, x_1-1, x_2+1|t) - x_2 p(\boldsymbol{x}|t)\right] \ .\end{aligned} \tag{244}$$

We define $p(\boldsymbol{x}|t) = 0$ for any $\boldsymbol{x} := (x_0, x_1, x_2)^T$ such that $x_0+x_1+x_2 \neq N$. We are specifically interested in the case where $p(\boldsymbol{x}|0) = \delta_{x_0,0}\delta_{x_1,0}\delta_{x_2,N}$, i.e., where initially all switches are fully marked with probability one.

#### 8.2 Transition probability

We can solve the $N$ Crick switch model the same way we solved the system consisting of $N$ two-state switches. Consider a random variable $\boldsymbol{z} = (z_1, z_2, z_3)^T$ which can take one of three possible one-hot values: $(1,0,0)^T$, $(0,1,0)^T$, or $(0,0,1)^T$. Assume that $\boldsymbol{z}$ takes these values with probabilities $p_0(t)$, $p_1(t)$, and $p_2(t)$ that match those of the Crick switch. (The first represents the unmarked probability, the second the partially marked probability, and the third the fully marked probability).

Consider $N$ independent and identically distributed variables of this kind, and the combination $\boldsymbol{x} := \boldsymbol{z}_1 + \cdots + \boldsymbol{z}_N$. Because $\boldsymbol{x}$ is a sum of independent and identically distributed variables with a categorical distribution, $\boldsymbol{x}$ has a multinomial distribution:

$$\boxed{p(\boldsymbol{x}|t) = \binom{N}{x_0, x_1, x_2} p_0(t)^{x_0} p_1(t)^{x_1} p_2(t)^{x_2}} \ . \tag{245}$$

Moreover, it satisfies the desired initial condition since $p(\boldsymbol{x}|0)$ equals zero except when $x_0 = x_1 = 0$ and $x_2 = N$. As before, we can show directly that this distribution satisfies the mult-switch CME (Eq. 244). Note,

$$\begin{aligned}
p(x_0 - 1, x_1 + 1, x_2|t) &= \frac{x_0}{x_1 + 1}\,\frac{p_1(t)}{p_0(t)}\;p(\boldsymbol{x}|t) \\
p(x_0 + 1, x_1 - 1, x_2|t) &= \frac{x_1}{x_0 + 1}\,\frac{p_0(t)}{p_1(t)}\;p(\boldsymbol{x}|t) \\
p(x_0, x_1 + 1, x_2 - 1|t) &= \frac{x_2}{x_1 + 1}\,\frac{p_1(t)}{p_2(t)}\;p(\boldsymbol{x}|t) \\
p(x_0, x_1 - 1, x_2 + 1|t) &= \frac{x_1}{x_2 + 1}\,\frac{p_2(t)}{p_1(t)}\;p(\boldsymbol{x}|t)
\end{aligned} \tag{246}$$

and

$$\begin{aligned}
\frac{\partial p(\boldsymbol{x}|t)}{\partial t} &= \binom{N}{x_0, x_1, x_2}\left\{x_0\frac{\dot{p}_0(t)}{p_0(t)} + x_1\frac{\dot{p}_1(t)}{p_1(t)} + x_2\frac{\dot{p}_2(t)}{p_2(t)}\right\} p_0(t)^{x_0} p_1(t)^{x_1} p_2(t)^{x_2} \\
&= \left\{\frac{x_0}{p_0(t)}\left[-2\epsilon p_0(t) + \gamma p_1(t)\right] + \frac{x_1}{p_1(t)}\left[2\epsilon p_0(t) - (\epsilon + \gamma + k)p_1(t) + 2\gamma p_2(t)\right]\right. \\
&\quad \left. + \frac{x_2}{p_2(t)}\left[(\epsilon + k)p_1(t) - 2\gamma p_2(t)\right]\right\} p(\boldsymbol{x}|t)\ .
\end{aligned} \tag{247}$$

Notice that several terms on the right-hand side are already present in Eq. 244:

$$\left[-2\epsilon x_0 - (\epsilon + \gamma + k)x_1 - 2\gamma x_2\right]\, p(\boldsymbol{x}|t)\ . \tag{248}$$

All we need to show is that the remaining terms in Eq. 247 correspond to the remaining terms in Eq. 244. This is easy to show:

$$\begin{aligned}
\gamma x_0 \frac{p_1(t)}{p_0(t)} p(\boldsymbol{x}|t) &= \gamma(x_1 + 1)p(x_0 - 1, x_1 + 1, x_2|t) \\
2\epsilon x_1 \frac{p_0(t)}{p_1(t)} p(\boldsymbol{x}|t) &= 2\epsilon(x_0 + 1)p(x_0 + 1, x_1 - 1, x_2|t) \\
2\gamma x_1 \frac{p_2(t)}{p_1(t)} p(\boldsymbol{x}|t) &= 2\gamma(x_2 + 1)p(x_0, x_1 - 1, x_2 + 1|t) \\
(\epsilon + k)x_2 \frac{p_1(t)}{p_2(t)} p(\boldsymbol{x}|t) &= (\epsilon + k)(x_1 + 1)p(x_0, x_1 + 1, x_2 - 1|t)\ .
\end{aligned} \tag{249}$$

Hence, our multinomial $p(\boldsymbol{x}|t)$ solves Eq. 244 with the correct initial condition.

#### 8.3 Moments

Since $p(\boldsymbol{x}|t)$ is multinomial, its low-order moments are easy to compute. We have

$$\begin{aligned}
\mathbb{E}[x_i] &= p_i(t) \\
\text{var}(x_i) &= N p_i(t)[1 - p_i(t)] \\
\text{Cov}(x_i, x_j) &= -N p_i(t) p_j(t) \qquad\qquad (i \neq j)\ .
\end{aligned} \tag{250}$$

##### 8.4 Eigenvalues

As we noted in Sec. 8, knowing the transition probability allows us to deduce the eigenvalues of the multi-switch model's Hamiltonian. In short, we know that

$$p(\boldsymbol{x}|t) = \binom{N}{x_0, x_1, x_2} p_0(t)^{x_0} p_1(t)^{x_1} p_2(t)^{x_2} \tag{251}$$

and that each $p_i(t)$ has the form (see Sec. 6)

$$p_i(t) = \pi_i - c_1 v_{1i} e^{\lambda_1 t} - c_2 v_{2i} e^{\lambda_2 t} \ . \tag{252}$$

We can expand the transition probability to write it as a sum of terms each associated with a different exponential. The exponentials that appear in this expansion have the form

$$e^{(n_1 \lambda_1 + n_2 \lambda_2) t} \tag{253}$$

where $n_1$ and $n_2$ are nonnegative integers with $n_1 + n_2 \leq N$. Hence, the eigenvalues of this multi-switch model are

$$\boxed{\lambda_{n_1, n_2} = n_1 \lambda_1 + n_2 \lambda_2} \tag{254}$$

for all $n_1, n_2 \in \{0, 1, ..., N\}$ such that $n_1 + n_2 \leq N$. There are $\binom{N+2}{2}$ such pairs of nonnegative integers, which is precisely the number of eigenvalues we expect $\boldsymbol{H}$ to have.

#### 9 Realistic Crick-like switch

In the main text, we consider a more realistic model of a Crick-like switch that explicitly includes both (i) the binding and unbinding of monomers, and (ii) monomer gain and loss. This model has six distinct chemical species: three types of monomers ($X_0$, $X_1$, and $Z$) and three types of dimers ($C_{00}$, $C_{10}$ and $C_{11}$).

- $X_0$ is an unmarked monomer and $X_1$ is a marked monomer. The transitions between these two species recapitulate those of the two-state model.
- $C_{00}$ is an unmarked dimer, $C_{10}$ is a partially marked dimer, and $C_{11}$ is a fully marked dimer. The transitions between these three species recapitulate those of the Crick switch.
- $Z$ is a reservoir species which accounts for the fact, in biophysically realistic systems, the number of switches may not be conserved across time, but can fluctuate. Both $X_0$ and $X_1$ can transition to $Z$, in which case we say that a monomer has been 'lost'. $Z$ can transition to $X_0$, in which case we say that a monomer has been 'gained'.

The full reaction list of the model is as follows:

$$\begin{aligned}
Z &\xrightarrow{\alpha} X_0 \\
X_0 &\xrightarrow{\beta} Z \\
X_1 &\xrightarrow{\beta} Z \\
X_0 &\xrightarrow{\epsilon} X_1 \\
X_1 &\xrightarrow{\gamma} X_0 \\
X_0 + X_0 &\xrightarrow{b_f} C_{00} \\
X_0 + X_1 &\xrightarrow{b_f} C_{10} \\
X_1 + X_1 &\xrightarrow{b_f} C_{11} \\
C_{00} &\xrightarrow{b_r} X_0 + X_0 \\
C_{10} &\xrightarrow{b_r} X_1 + X_0 \\
C_{11} &\xrightarrow{b_r} X_1 + X_1 \\
C_{00} &\xrightarrow{2\epsilon} C_{10} \\
C_{10} &\xrightarrow{\epsilon+k} C_{11} \\
C_{11} &\xrightarrow{2\gamma} C_{10} \\
C_{10} &\xrightarrow{\gamma} C_{00}
\end{aligned} \tag{255}$$

The parameters $\epsilon$, $\gamma$, and $k$ have the same meaning as before: they correspond to the rates of marking noise, (mark) turnover, and catalysis, respectively.

The parameters $b_f$ and $b_r$ correspond to the rates of monomer-monomer binding and unbinding. In our simulations, we assume $b_f$ is much larger than $b_r$, since otherwise the number of monomers would tend to be larger than the number of dimers. We assume elementary propensity functions; for example, the $X_0$-$X_0$ binding rate is $(b_f/2)x_0(x_0 - 1)$.

The parameters $\alpha$ and $\beta$ correspond to the rates of monomer gain and loss. In our simulations, we assume $\alpha$ is much larger than $\beta$, since otherwise the number of reservoir molecules would tend to be larger than the number of other molecules.

Generally, we expect that this model reduces to the Crick switch when

- $b_f$ and $b_r$ are both small and $b_f \gg b_r$ (so that dimers tend to be stable, and binding dynamics is irrelevant on the time scales of marking and unmarking dynamics); and
- $\alpha$ and $\beta$ are both small and $\alpha \gg \beta$ (so that the number of switch-related molecules is roughly constant and close to the total number of molecules, and does not fluctuate on the time scales of marking and unmarking dynamics).

This model's bimolecular reactions (i.e., all reactions of the form $X_i + X_j \to C_{ij}$) make it not analytically tractable, so we study this model in the main text by numerically solving its master equation.

Finally, it is worth noting that this model's reaction list implies a conservation law. If we let $x_0$, $x_1$, $z$, $c_{00}$, $c_{10}$, and $c_{11}$ denote the number of each type of species, the sum

$$z + x_0 + x_1 + 2(c_{00} + c_{10} + c_{11}) = \text{const.} \tag{256}$$

remains the same across time. Intuitively, this means that the total number of molecules (with each dimer counted as two molecules) remains constant. One implication of this conservation law is that the number of possible states is finite, and depends on the model's initial condition (since it effectively sets the total number of molecules).
